# Loss of Nance-Horan Syndrome b (*nhsb*) prevents expansion growth of retinal progenitor cells by selective up-regulation of *Δ113p53*

**DOI:** 10.1101/775171

**Authors:** Paul J. Vorster, John Ojumu, Amanda J. G. Dickinson, Gregory S. Walsh

**Affiliations:** Department of Biology, Virginia Commonwealth University, 1000 W. Cary Street, Richmond, VA 23284, USA

**Author notes:** email addresses: Paul Vorster, John Ojumu, Amanda Dickinson, Gregory Walsh.

**Keywords:** Nance-Horan Syndrome, nhsb, Δ113p53, p53, retinogenesis, zebrafish

## Abstract

The regulation of cell cycle progression and differentiation in retinal progenitor cells is a fundamental feature controlling organ size of the eye in vertebrates. Nance-Horan Syndrome (NHS) is a rare X-linked disorder caused by mutations in the *NHS* gene. Dysmorphic features of NHS include severe congenital cataracts, micropthalmia, facial dysmorphisms, and visual impairment. In this study we report an evolutionarily conserved role for NHS in vertebrate retinogenesis. Loss of function of *nhs* leads to small eye size in both zebrafish and *Xenopus tropicalis,* marked by reduced proliferation but not cell death. Transcriptome analysis of *nhs* morphant zebrafish eyes revealed a marked upregulation in *Δ113p53,* an isoform of *p53*, concomitant with a selective upregulation of p53 responsive genes that inhibit cell cycle progression but not apoptosis. Our data supports a model where Nhs is a negative regulator of *Δ113p53* expression and exerts its function through regulation of the p53 pathway to promote expansive growth of retinal progenitor cells prior to differentiation.

## Introduction

A fundamental question in biology is how organ size is controlled during development. Organogenesis is regulated by a balance between progenitor cell cycle progression, self-renewal, differentiation, and programmed cell death. The neural retina serves as a model system that illustrates this general concept. The retina develops as an outpouching of the neuroepithelium and grows through symmetric proliferative divisions of retinal progenitor cells (RPCs) followed by a switch to asymmetric neurogenic divisions (Fadool and Dowling 2008; Willardsen and Link 2011; He et al. 2012; Boije et al. 2014). In the zebrafish retina, early RPCs undergoing symmetric cell divisions progress through the cell cycle slowly but then switch to a rapidly proliferating phase in which the cell cycle duration markedly decreases (Hu and Easter 1999; Li et al. 2000; He et al. 2012). This serves to rapidly increase the progenitor pool in advance of asymmetric neurogenic divisions in which some daughter cells exit the cell cycle to differentiate into neurons (He et al. 2012). Thus, coordinated cell proliferation and differentiation result in the correct number of cells that collectively contribute to the volume and size of the retina (Joseph and Hermanson 2010; Hindley and Philpott 2012). However, the molecular mechanisms that regulate cell cycle progression in retinal progenitor cells are not completely understood (He et al. 2012; Boije et al. 2014).

Nance-Horan Syndrome (NHS) is an X-linked developmental disorder characterized by severe congenital cataracts, dental anomalies, dysmorphic facial features and mental retardation (Brooks et al. 2004; Ramprasad et al. 2005; Coccia et al. 2009). Additional features of the disease include micropthalmia and microcornea (Florijn et al. 2006; Huang et al. 2007). NHS is caused by mutations in the *NHS* gene found at the distal end of the X chromosome. The most frequent mutations associated with NHS are nonsense mutations predicted to result in a truncated protein (Burdon et al. 2003; Ramprasad et al. 2005; Florijn et al. 2006; Huang et al. 2007; Sharma et al. 2008; Coccia et al. 2009; Chograni et al. 2011; Tug et al. 2013; Hong et al. 2014; Shoshany et al. 2017; Tian et al. 2017), but insertions, small deletions, and a missense mutations have also been reported (Accogli et al., 2017; Brooks et al., 2004; Burdon et al., 2003; Chograni et al., 2011; Florijn et al., 2006; Li et al., 2015). Affected males have severe bilateral congenital dense nuclear cataracts that lead to profound vision loss and usually require surgery at an early age (Burdon et al. 2003; Huang et al. 2007; Sharma et al. 2008; Coccia et al. 2009). Despite cataract surgery during infancy, many NHS patients continue to exhibit visual impairment that may be caused by an underlying retinopathy (Mathys et al. 2007; Ding et al. 2009). It has been demonstrated that NHS protein is expressed in midbrain, lens, tooth and retina (Burdon et al. 2003; Sharma et al. 2006).

The p53 tumor suppressor protein has an established role as a cellular stress sensor, responding to a variety of insults such as DNA damage, nutrient starvation, nucleolar and ribosomal stress, and hypoxia (Vousden and Lane 2007; McCubrey et al. 2017). Evidence suggests that p53 functions to arrest cell cycle progression allowing time for DNA repair machineries to restore genome integrity before replication. If deleterious conditions are too severe, p53 activates cell death pathways. p53 performs many of its functions though downstream transcriptional activation of target genes, such as *cdkn1a*/*p21^CIP1/WAF1^* (p21) to halt cell cycle at the G1 phase and *bax* to execute programmed cell death pathways (Vogelstein et al. 2000; Harris and Levine 2005).

In addition to the full-length p53 protein, the *p53* gene locus encodes for several isoforms generated by alternative promoter usage or alternative spicing Bourdon et al. 2005; Joruiz and Bourdon 2016). A prominent variant *Δ133p53* in humans is transcribed from an internal promoter in intron 4 and is evolutionarily conserved in zebrafish (termed *Δ113p53*), suggesting an important role in p53 functions (Bourdon et al. 2005; Chen et al. 2005). *Δ113p53* encodes an N-terminally truncated protein with a deletion of the transactivation domains and a partial deletion of the DNA-binding domain (Bourdon et al. 2005; Chen et al. 2005; Chen and Peng 2009). *Δ113p53* is itself a p53 target gene that is strongly upregulated following DNA damage, due to binding of full-length p53 to p53 response elements within intron4 of the *p53* locus (Bourdon et al. 2005; Chen et al. 2005; Marcel et al. 2010; Aoubala et al. 2011). Previous work showed that Δ113p53 functions to protect cells from apoptosis by differentially modulating p53 response genes, promoting the expression of cell cycle arrest genes while antagonizing p53-regulated apoptotic activity (Chen et al. 2009; Aoubala et al. 2011; Ou et al. 2014). Thus, the *Δ113p53/Δ133p53* variants function evolutionarily to promote cell survival when genome integrity is compromised to allow for DNA repair to take place (Gong et al. 2015).

Although p53 has a well-established role in maintaining genome stability to prevent tumor progression, a role for the p53 pathway in regulating development and organogenesis is less well understood. Loss-of-function mutations in p53 at first appeared to result mostly in viable offspring with no obvious developmental defects (Donehower et al. 1992). However, a number of p53 mutant mice die in utero due to exencephaly, a defect in neural tube closure due to overgrowth of neural progenitors in the midbrain, due at least in part to decreased apoptosis of neural progenitors (Sah et al., 1995). On the other hand, unrestrained activity of p53 can inappropriately induce p53 target genes and trigger cell cycle arrest and/or cell death during development (Jacobs et al. 2004; Dugani et al. 2009). For instance, loss of mdm2 or mdm4, negative regulators of the p53 protein, cause early embryonic lethality in a p53-dependent manner (Marine et al. 2006). More modest levels of p53 activation are associated with defects similar to those seen in patients with CHARGE syndrome (Van Nostrand et al. 2014). Recent work in zebrafish identified *digestive organ expansion factor* (*def*) as a pan-endodermal-enriched factor that is essential for expansive growth of digestive organs (Chen et al. 2005). Analysis of *def* null mutants revealed a selective regulation of the p53 isoform Δ113p53, that contributed to cell cycle arrest leading to reduced size and defective digestive organ development (Chen et al. 2005, 2009).

Here, we report a role for NHS in the development of the vertebrate retina. Knockdown of *nhs* in zebrafish and *frog* resulted in a reduced eye size. We show the small retinal size caused by *nhs* deficiency is due a decrease in RPC proliferation. The small retina, caused by nhs deficiency, is due a proliferation restriction mediated by the elevated expression of *Δ113p53*. We hypothesize that *nhs* functions in RPCs to limit p53 pathway induction as neural progenitors undergo rapid cell division.

## RESULTS

### The NHS family of genes

The Nance-Horan Syndrome family of genes is conserved among vertebrates including humans, mice, frogs, and zebrafish. Members of the NHS gene family include the prototypical NHS, as well as NHS-like 1 (NHSL1) and NHS-like 2 (NHSL2). The NHS genes are orthologs of the *Drosophila* gene GUK-holder (gukh)(Katoh and Katoh 2004)(Fig. 1A). Although most vertebrates express three *NHS* paralogs, (*NHS*, *NHSL1* and *NHSL2*), zebrafish have two copies of each paralog (*nhsa*, *nhsb*, *nhsl1a*, *nhsl1b*, *nhsl2a*, and *nhsl2b)* due to a genome duplication event in the teleost lineage (Postlethwait et al. 1998, 2000)(Fig. 1A).

**Fig. 1.**
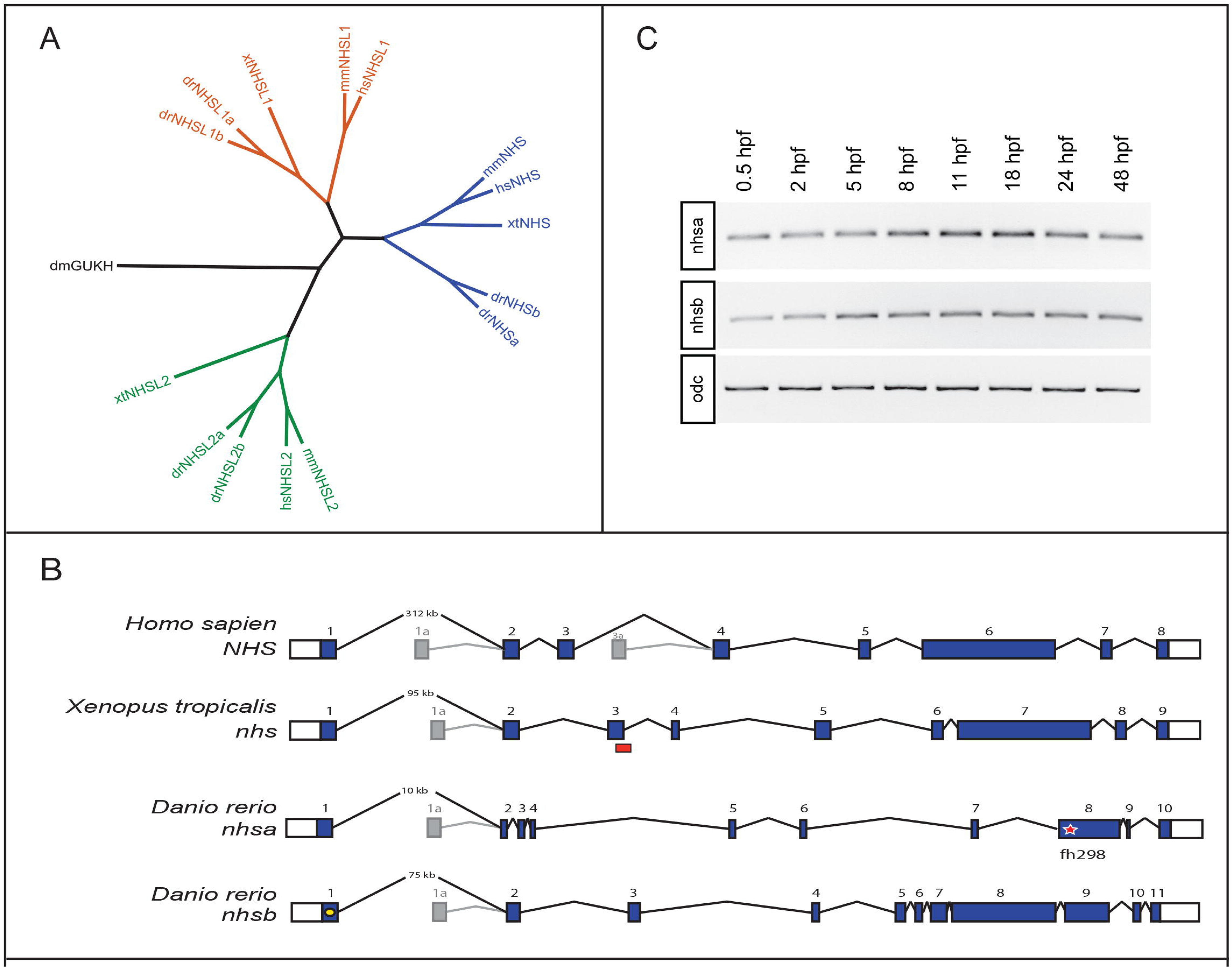
Phlyogeny and genomic architecture of NHS family genes. (A) Phylogenetic tree for the nhs family of genes including nhs, nhs-like 1 (nhsl 1), and nhs-like 2 (nhsl2) and their common homolog in Drosophila, guanylate kinase holder (gukh). Full protein coding sequences were used for each family member.The initial letter hs denotes human, mm mouse, xt Xenopus tropicalis, dm Drosophila melanogaster, and dr Danio rerio. (B) Conserved genomic structure of nhs genes in vertebrates showing relative positions of exons and alternative translation start sites. Dark boxes represent exons, open boxes represent untranslated regions (UTRs), star represents location of nonsense mutation (fh298) in nhsa, red boxes represent predicted morpholino oligonucleotide binding region, yellow dot represents nhsb guide RNA sequence target. (C) Temporal expression patterns of nhsa and nhsb in zebrafish embryos. Polymerase chain reaction from zebrafish cDNA at various timepoints using primers designed to nhsa, nhsb, and the control ornithine decarboxylase (odc).

In this study, we focused only on NHS orthologs. Previously, it was noted that the human NHS gene encompassed an unusually long distance between exon1 and exon2 (Brooks et al. 2004, 2010b). Comparison of other vertebrate genomes showed that the genomic architecture of the NHS gene remained conserved among human, mouse, frog and zebrafish with comparable number of exons. The large distance between exon1 and exon2 of NHS was also detected in other vertebrate genomes (Fig. 1B). The human NHS gene generates multiple protein isoforms by virtue of alternative promoter usage and alternative splicing (Huang et al. 2006; Brooks et al. 2010a). Isoforms arising from the alternative translational start site in exon1a appear to be evolutionarily conserved across vertebrates, as evidenced by sequencing of expressed sequence tags (EST) libraries (Fig. 1B). However, only NHS isoforms encoded from the first translational start site in exon1 are predicted to be essential for development, since nonsense mutations in exon1 in humans leads to NHS (Burdon et al. 2003; Brooks et al. 2004).

We examined the temporal expression pattern of *nhs* paralogs during zebrafish development and find that both *nhsa* and *nhsb* are present in embryos before 2 hours post fertilization (hpf), suggesting that both transcripts are loaded maternally (Fig. 1C). We find that both *nhs* paralogs are expressed through all stages of development (Fig. 1C).

### Zebrafish *nhsb* is expressed in the developing eye and is required for normal retinogenesis

To determine the role of NHS in zebrafish development, we first examined the *nhsa^fh298^* mutants generated by Tilling (Draper et al. 2004). *nhsa^fh298^* carries a truncating nonsense mutation at position Q217* (C4524T) (Fig. 1B). Examination of embryos derived from an incross of *nhsa^fh298/+^* heterozygotes revealed no discernable morphological differences among siblings, suggesting that *nhsa* is dispensable for early development (Fig. S1).

We therefore decided to focus our efforts on zebrafish *nhsb* during embryogenesis. Using *in situ* hybridization, we found expression of *nhsb* in the eye during key stages of retinogenesis. Expression of *nhsb* was seen within proliferating retinal progenitor cells (RPCs) and in the lens at 24 hpf (Fig. 2A). By 48 hpf and 72 hpf, *nhsb* expression was downregulated in proliferating RPCs and not found in the ciliary marginal zone (CMZ). Instead, *nhsb* expression was detected in differentiated cells outside of the CMZ, with staining present in the retinal ganglion cell layer and inner plexiform layer of the retina (Fig. 2B-D).

**Fig. 2.**
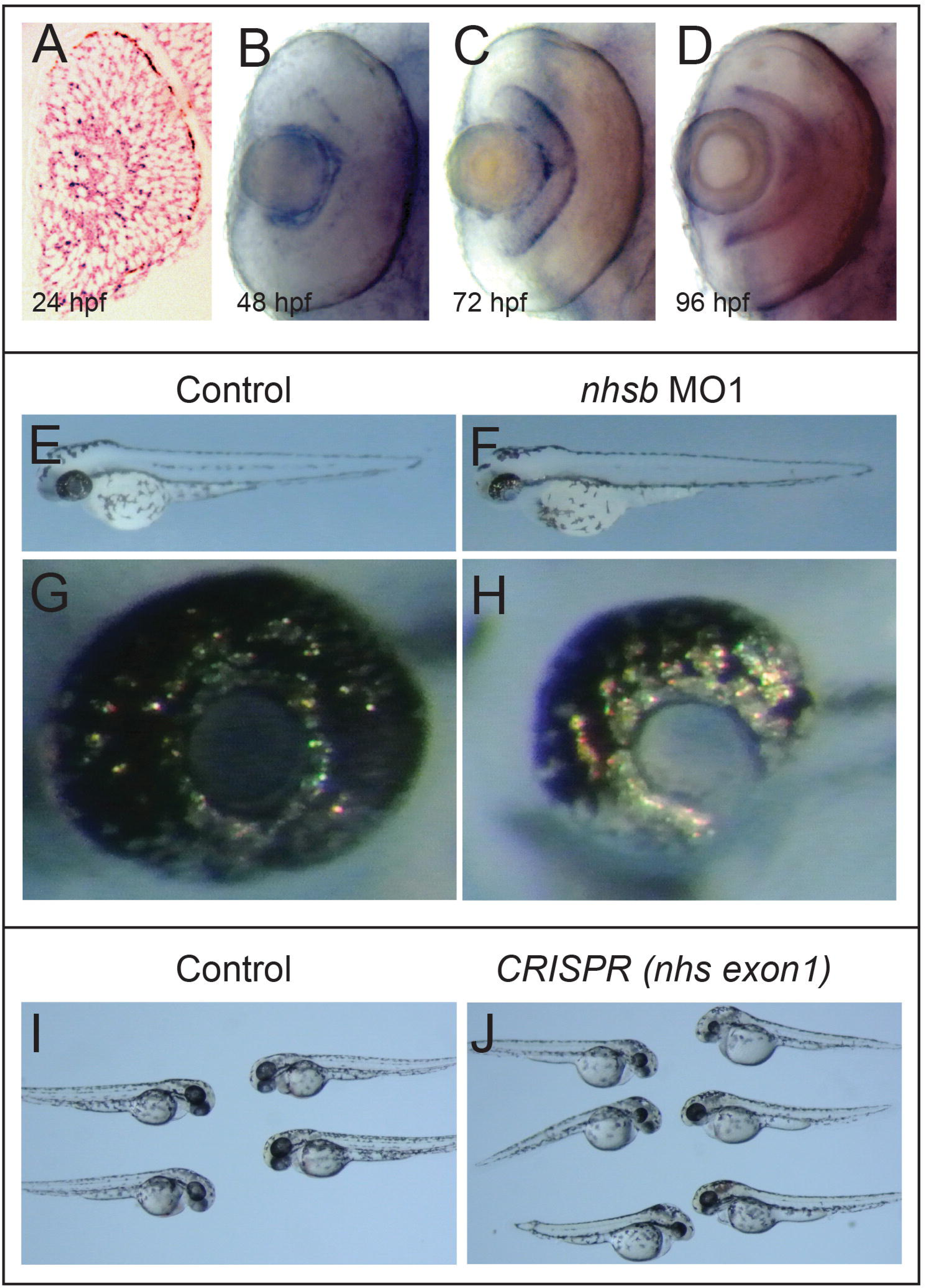
Loss of nhsb function leads to a reduced eye size in zebrafish. Whole-mount in situ hybridization using a nhsb riboprobe to examine the expression patterns nhsb at 24hpf (A), 48 hpf (B), 72 hpf (C), 96 hpf (D). Stereomicroscope photomicrograph of embryos at 72 hpf from control (E) and nhsb morphants (F). In comparison to control embryos (E,G), embryos injected with nhsb MO1 display a smaller eye size. Stereomicroscope photomicrographs of embryos at 48 hpf from control (I) and F0 embryos injected with Cas9/gRNA targeted to exon1 of nhsb (J).

We next examined a role for *nhsb* in zebrafish embryo development. To accomplish this, we designed splice-blocking antisense morpholino oligonucleotides (MO) to the exon2-intron2 boundary of the *nhsb* gene (*nhsb* MO1) (Fig. 1B). We found that *nhsb* MO1-injected embryos initially look indistinguishable from control embryos until 24 hpf, suggesting that knockdown of *nhsb* does not affect early development, eye field specification and/or separation. By 72 hpf, we observed a striking difference in the size and shape of the retina in *nhsb* morphants as compared to controls (Fig. 2E-H). *nhsb* morphants exhibited a small retina located on top of the lens (Fig. 2H). We did not observe any lens defects in *nhsb* MO1-injected embryos that were reminiscent of cataracts as observed in humans with Nance-Horan Syndrome (Burdon et al. 2003; Brooks et al. 2004; Sharma et al. 2008; Coccia et al. 2009). We confirmed that injection of *nhsb* MO1 caused a mis-splicing of the *nhsb* transcript leading to retention of intron2 (Fig. S2). To ensure that the small eye phenotype was specific to knockdown of *nhsb*, we injected a second splice-blocking *nhsb* MO (*nhsb* MO2), targeted to the intron2-exon3 (I2E3) boundary (Fig. 1B). We found that *nhsb* MO2-injected embryos exhibited the same phenotype as that observed with *nhsb* MO1 (hereafter referred to as *nhsb MO)* (Fig. S3).

To further test whether the small eye phenotype was specific to disruptions in the *nhsb* gene, we used CRIPSR/Cas9 to generate insertion/deletion (indel) mutations at the *nhsb* locus. We designed a single-stranded guide RNA (gRNA) to target exon1 of the *nhsb* gene (Fig 1B). It was previously reported that injection of increasing amounts of gRNA can lead to bi-allelic disruptions in the targeted gene (Jao et al. 2013). We found that co-injection of *nhsb^ex1^ gRNA* and *Cas9* mRNA into one-cell stage embryos led to growth defects in the eye of injected founder (F0) embryos (14%, *n* = 245), similar to that seen in *nhsb* morphants (Fig 2I,J). We chose five injected embryos displaying the small eye phenotype and validated indel mutations at the *nhsb* locus using a T7 endonuclease I (T7E1) assay; all assayed embryos showed high mutagenesis rates at the *nhsb* locus (Fig. S4). Taken together, our results demonstrate that the *nhsb* gene is expressed in the retina during key stages of retinogenesis, and that loss-of-function of *nhsb* leads to reduced retinal size in zebrafish.

### Expression and function of NHS is conserved in *Xenopus tropicalis*

To test whether gene expression was conserved in other vertebrates, we examined a role for *nhs* in retina development in *Xenopus tropicalis. We chose X. tropicalis* because they are diploid and possess only one *nhs* gene. First, we made anti-sense *in situ* hybridization probes against the *Xenopus tropicalis nhs* gene (Fig. 1B). Similar to zebrafish *nhsb*, the expression of *nhs* was detected in the retinal progenitor cells and lens of *Xenopus tropicalis* eyes at embryonic stage 31 (Fig. 3A). As retinogenesis proceeded, *nhs* expression in RPCs faded and was found concentrated in the plexiform layers of the retina (Fig. 3A-D).

**Fig. 3.**
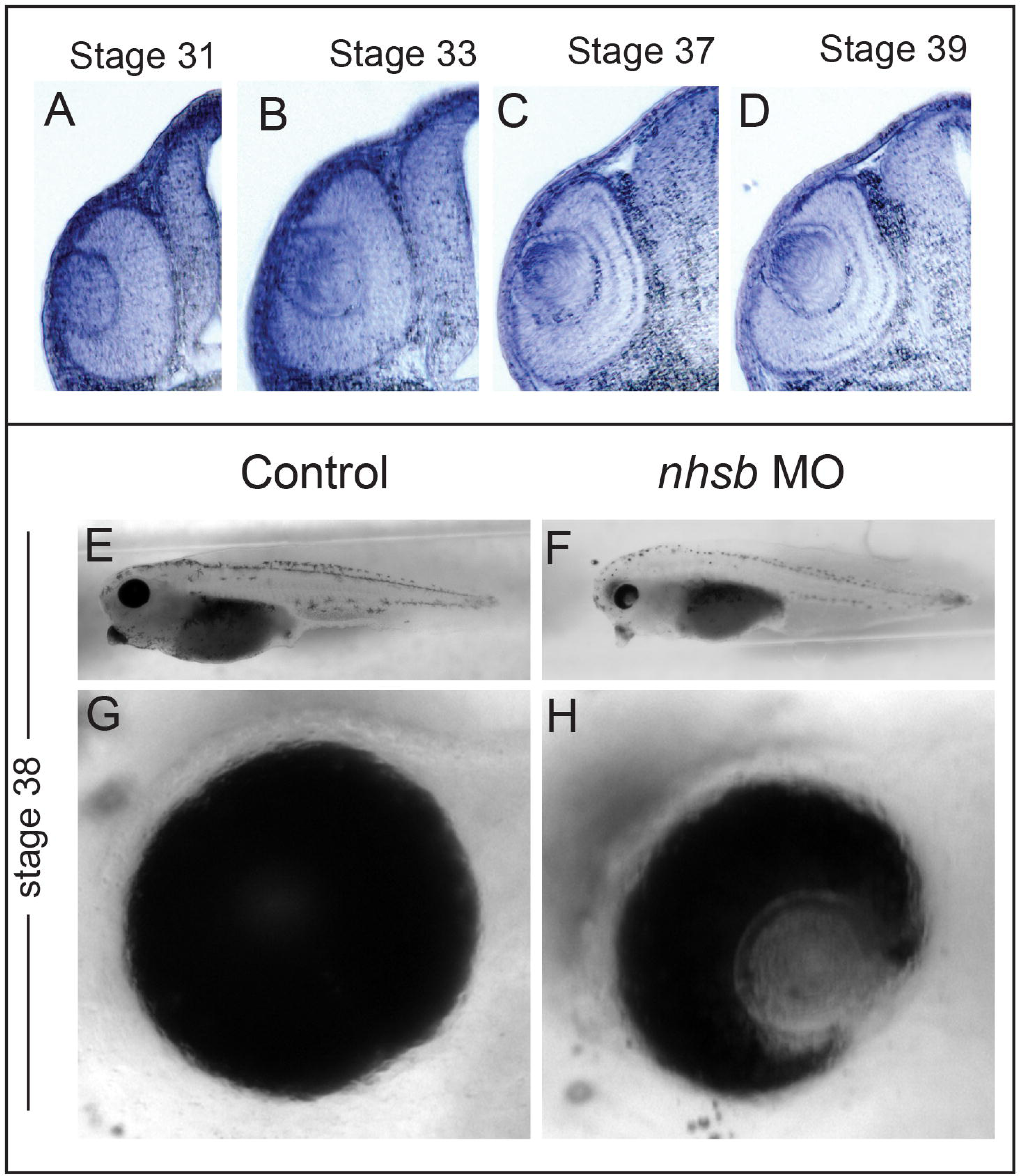
Loss of nhs function causes a small eye in frog. Vibratome sections of Xenopus tropicalis embryos stained by whole-mount in situ hybridization using a probe against nhs at Stage 31 (A), Stage 33 (B), Stage 37 (C), and Stage 39 (D). Stereomicro­scope photomicrographs of Xeno­pus tropicalis embryos at stage 38 of control (E,G) and nhs MO-inject­ed embryos (F,H). Note the small eye phenotype in nhs morphant frogs.

To determine whether the role of Nhs in retina development was evolutionarily conserved among vertebrates, we designed splice-blocking MOs targeting the exon3-intron3 boundary of *Xenopus tropicalis nhs* (*nhs* MO). We co-injected *nhs* MO along with rhodamine-dextran into one cell of a two-cell stage embryo. This experimental approach provided us with an internal control, since only half the embryo received *nhs* MO. We found that 96% of embryos (*n* = 56) injected with *nhs* MO had small retinas as compared to eyes on the contralateral side that did not receive morpholino (Fig. 3E-H). The efficacy of knockdown by the *nhs* MO in *Xenopus* was validated by RT-PCR (Fig. S5). These findings suggest that the role of NHS in retinal development is evolutionarily conserved since knockdown of *nhs* in frog and knockdown of *nhsb* in zebrafish results in a similar small eye phenotype. Taken together, the expression and function of Nhs during key stages of retinogenesis is conserved between *Xenopus tropicalis* and *Danio rerio*.

### Nhs is required for proliferative growth of RPCs in the retina

To further investigate the function of Nhs in retinal development we performed the rest of our analysis on zebrafish *nhsb* morphants. The earliest manifestation of the reduced eye size in *nhsb* morphants occurred between 24hpf - 48 hpf. This corresponds to the period of rapid RPC proliferation followed by neurogenesis and differentiation (He et al. 2012). To determine whether the reduced eye size was due to an increase in RPC cell death, we analyzed wild-type and *nhsb* morphant retinas for apoptotic cells. We evaluated cell death using an anti-cleaved caspase-3 antibody, a marker of apopotic cell death. We found virtually no dying cells in the retinas of either wild-type (0.187% (4/2137 cells, *n* = 6 embryos) or *nhsb*-deficient embryos (0.140%; 2/1427 cells, *n* = 6 embryos) at 36 hpf (Fig. 4A,B). Furthermore, *nhsb* morphant retinas did not display pyknotic morphology characteristic of dying cells (Fig. 4A,B). These results imply that the reduced eye size observed in *nhsb*-depleted embryos was not due to an increase in cell death.

**Fig. 4.**
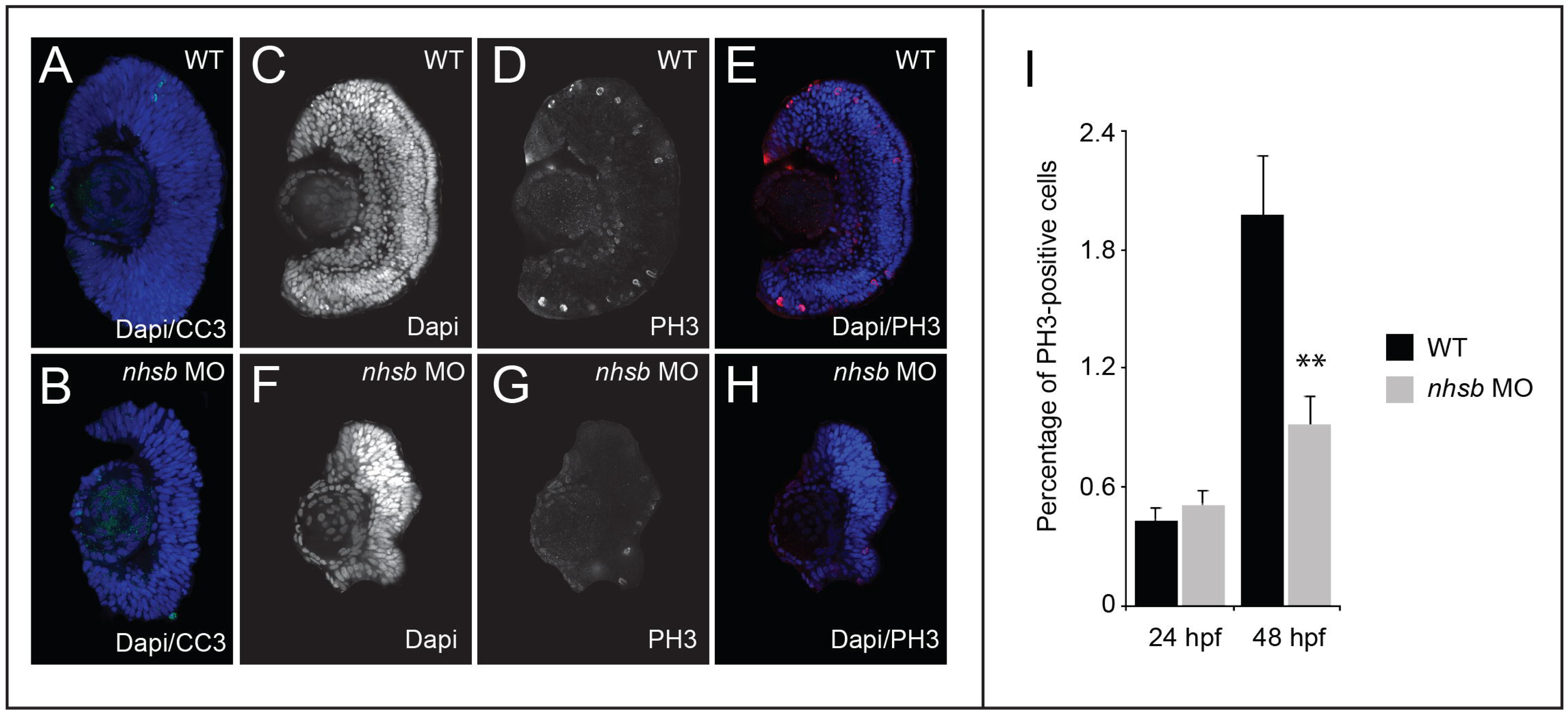
Reduced proliferation in retinal progenitor cells but not cell death following nhsb depletion. Cryosections of zebrafish retina stained for Dapi (blue) and anti-cleaved caspase-3 (CC3) (green) in control wild-type (A), and nhsb morphants (B). Despite having less cells, no increase in CC3 staining is evident in nhsb-depleted embryos compared to control. (C-H) Cryosections of zebrafish retina stained for Dapi (blue) and anti-phosphohistone3 (PH3) (red) at 48 hpf. (I) Quantitation of PH3-positive cells in the retina of control and nhsb morphants.The number of PH3-positive cells are significantly reduced in nhsb-morphant retinas compared to control wild-type retinas at 48 hpf indicating a decrease in proliferating cells when nhsb is depleted. (** p < 0.05, Student’s t test)

To determine whether the small eye phenotype observed in *nhsb* morphants was due to a defect in RPC proliferation, we analyzed cryosections stained with phospho-histone 3 (PH3) antibody, a marker of M-phase of the cell cycle. At 24 hpf, a timepoint in which no morphological difference in eye size was observed, we found no significant difference in the proportion of PH3-positive cells in *nhsb* morphants compared to wild-type embryos (wt, 0.43 ± 0.05%, *n* = 538 cells, 4 embryos; *nhsb* MO, 0.51 ± 0.04%, *n* =745 cells, 5 embryos)(Fig. 4B-I). At 48 hpf, however, the number of cells at M-phase was significantly fewer in the retina of *nhsb* morphants (0.92 ± 0.08%, *n* = 655 cells, 4 embryos) compared to wild-type (1.98 ± 0.18%, *n* = 2218 cells, 4 embryos) embryos (*p* < 0.05)(Fig. 4B-I). This suggests that reduced proliferation of RPCs may contribute to the lack of eye growth in the absence of *nhsb* function.

We next sought to determine whether differentiation of RPCs into neurons was affected by loss of *nhsb*. To accomplish this, we made use of *TgBAC(neurod1:EGFP*) transgenic zebrafish that expresses GFP under control of the promoter for *neuroD* (Thomas et al. 2012). This transgene is not expressed in proliferating RPCs, but is expressed in a subset of retinal neurons undergoing differentiation (Thomas et al. 2012). Some of the first differentiating neurons identified with this transgenic line can be seen around 34 hpf (Fig. 5A-C). Many GFP-positive neurons are present in the retina of wild-type embryos at this timepoint, and this number increased by 48 hpf (Fig. 4G-I). In contrast, by 34 hpf, no GFP-positive neurons are observed in *nhsb* morphants (Fig. 4D-F). By 48 hpf, however, a few GFP-expressing neurons can be visualized in *nhsb* morphant retinas (Fig. 4J-L), suggesting that the differentiation of neurons from RPCs is not blocked but is delayed in *nhsb*-depleted embryos. These findings are consistent with the idea that *nhsb* knockdown mainly affects expansive growth of RPCs but does not inhibit neuronal differentiation.

**Fig.5.**
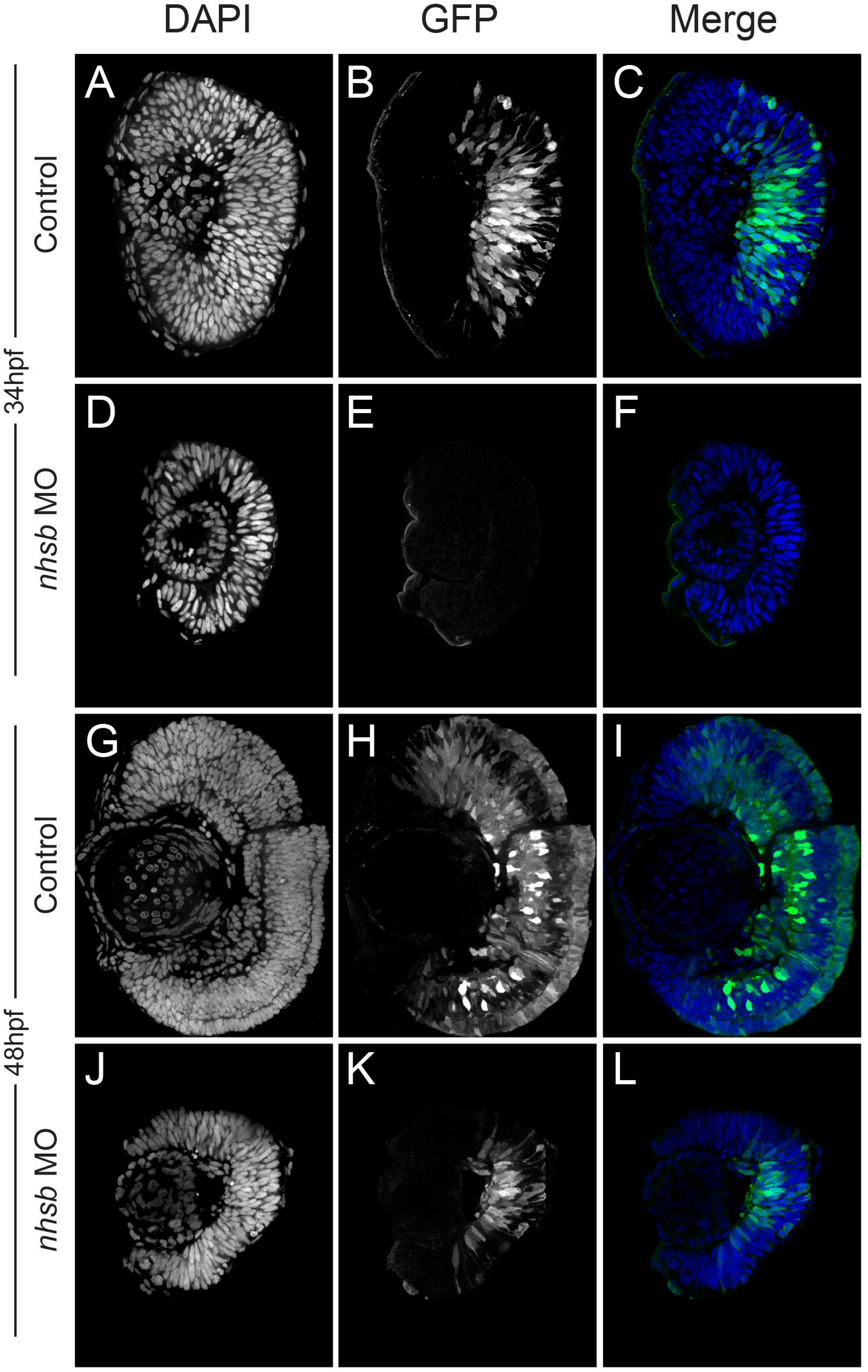
nhsb knockdown leads to a delay in neuronal differentiation. (A) Assessment of GFP-positive cells in cryosections from TgBAC(neurod1:EGFP) transgenic fish injected with nhsb MO at 34 hpf. At this timepoint, neurons are just differentiating in control embryos, whereas no GFP-positive cells were detected in nhsb morphant TgBAC(neurod1:EGFP) fish. (B) At 48 hpf, many neurons have differentiated in control embryos, whereas only a few neurons have terminally differentiated in the nhsb morphants TgBAC(neurod1:EGFP) fish.

### Loss of *nhsb* leads to a marked upregulation of *Δ113p53* and *p21*, but not full-length *p53*

To gain insight into the downstream function of Nhsb, we compared retinal expression profiles between wild-type and *nhsb* morphants. Total RNA was isolated from 40 zebrafish eyes dissected from wild-type and *nhsb* morphant embryos at 48 hpf for microarray analysis using Affymetrix zebrafish GeneChips. We used SAM analysis and found 1,212 significant genes that were differentially expressed in wild-type eyes compared to *nhsb* morphant eyes (Table S2A,B).

A subset of genes with significantly higher expression in *nhsb*-depleted eyes included genes that are markers of RPC identity such as *mz98/col15a1b* (*collagen typeXV, alpha 1b), ccnd1* (*cyclin D1*), *dlc* (*deltaC*), and *pold1* (*DNA polymerase delta 1).* (Fig. 6A). We also found a subset of genes whose expression was reduced in *nhsb* morphants compared to control, including markers of differentiated neurons such as *neuroD*, *opn1sw1* (*opsin1 short-wave-sensitive 1*), *rho* (*rhodopsin*), *calb2a* (*calbindin2a*), *snap25a* (s*ynaptosomal-associated protein, 25a*) (Fig. 6A). These findings are consistent with the idea that RPCs in *nhsb* morphant retinas proliferate slower, maintaining their identity as RPCs with delayed neuronal differentiation.

**Fig. 6.**
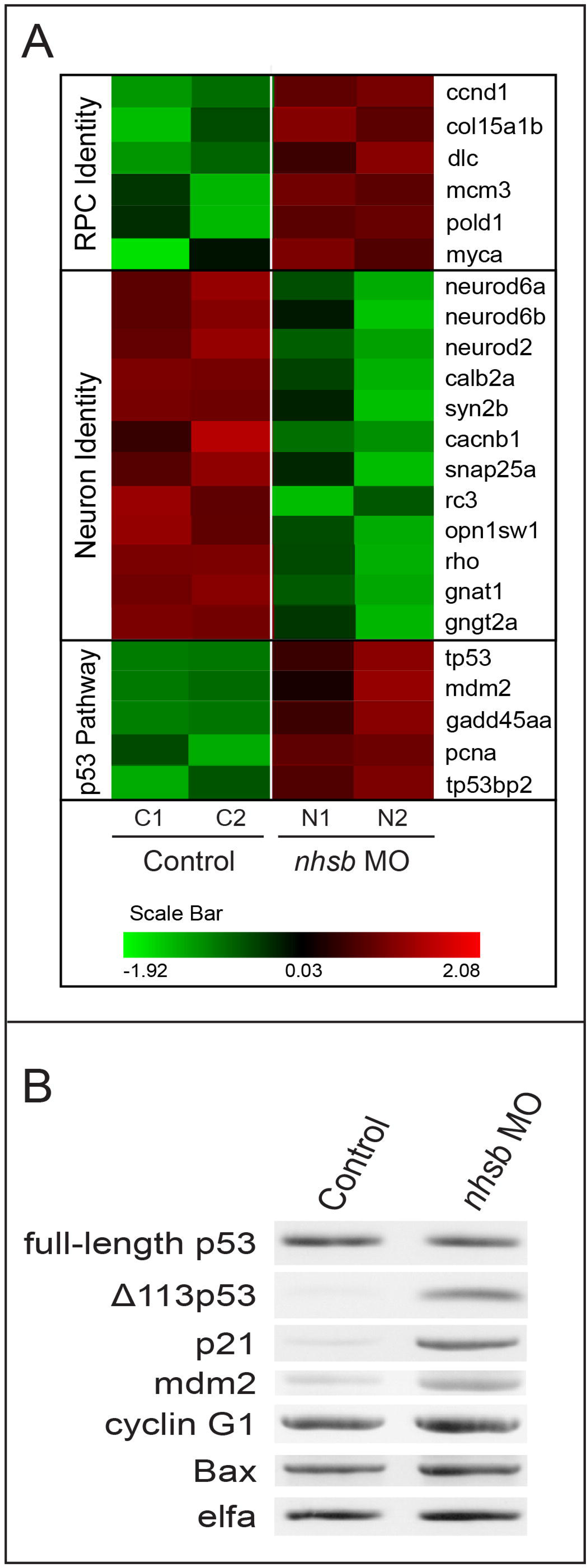
Δ113p53 but not full length p53 is upregulated in nhsb morphants. (A) Clustering and heat map of gene transcripts regulated by nhsb-depletion identified in microarray experiments. Red and green shading, respectively, indicates the highest and lowest expression levels, as indicated in the scale bar at the bottom of the figure. Following knockdown of nhsb, markers of RPC identity were elevated, whereas marker of terminally differentiating neurons were downregulated, and genes associated with the p53 pathway were epregulated compared to wild-type control retinas (B) Semi-quantitative RT-PCR examination of expression of various p53 response genes in nhsb morphant retina and wild-type control retinas. nhsb morphants had elevated levels of Δ113p53, p21, mdm2, and cyclin Gl, but full-length p53 and bax remained unchanged

Surprisingly, functional annotation analysis using the Database for Annotation, Visualization and Integrated Discovery (DAVID) identified a number of genes that were differentially expressed between wild-type and *nhsb*-depleted retinas with known roles in the regulation of cell cycle progression, including the tumor-suppressor gene *tp53* and its target gene *mdm2* (Fig. 6A). This suggests that loss of *nhsb* may regulate cell cycle progression through the p53 pathway.

To validate our microarray results, we performed semi-quantitative RT-PCR, using RNA extracted from dissected retinas of wild-type and *nhsb* morphants. We found that the levels of full-length *p53* transcripts were unchanged in *nhsb* morphants compared to wildtype controls, a result that differed from our microarray expression analysis (Fig. 6B). Whereas the microarray probe binds to the C-terminal *tp53* coding region, our RT-PCR primers amplify N-terminal *tp53* coding region from exon1 to exon4 (Table S1). However, using primers that specifically amplify the truncated *Δ113p53* transcript, we found a marked up-regulation of *Δ113p53* expression in *nhsb* morphant eyes compared to wild-type controls (Fig. 6B). Our results demonstrate that the elevated p53 transcripts detected by the Affymetrix probe, directed against the C-terminal coding region of p53, is due to the upregulation of the N-terminal truncated p53 variant *Δ113p53*, but not full-length *p53*.

We next asked whether other p53 target genes were affected by *nhsb* depletion. Semi-quantitative RT-PCR results showed that additional *p53* target genes were up-regulated in *nhsb* morphant eyes compared to wild-type eyes, including *mdm2, ccng1 (cyclinG1), and cdkn1a*/*p21^WAF1/CIP-1^* (p21), a cyclin dependent kinase inhibitor that blocks the G1-to-S-phase transition (Cazzalini et al. 2010)(Fig. 5B). On the other hand, expression levels of the pro-apoptotic factor *bax* did not show apparent differences between *nhsb* morphants and control eyes (Fig. 5B). Taken together, these data are consistent with our histological data that the reduced eye size in *nhsb* morphants is not caused by increased apoptosis but rather is due to a proliferation restriction imposed on RPCs by elevated expression of the p53-target gene and cell-cycle checkpoint protein, *p21*.

### Knockdown of *p53*, reduces *Δ113p53* and *p21*, and rescues the *nhsb* morphant phenotype

Previous data from *Chen* and colleagues demonstrated that the induction of *Δ113p53* was p53-dependent (Chen et al. 2009). Our data suggest that *nhsb* may be a negative regulator of *Δ113p53* expression. Since Δ113p53 is itself a transcriptional target of full-length *p53* that binds to promoter elements in intron 4 (Chen et al. 2009), we reasoned that the small-eye phenotype seen in *nhsb* morphants may be rescued by knock-down of full-length *p53*. To determine whether full-length p53 is required for the elevated levels of *Δ113p53* caused by *nhsb* depletion, we co-injected *nhsb* MO2 with *p53* MO^ATG^ that specifically targets the translation start site of the full-length p53 protein (Robu et al. 2007). We found that co-injection of *nhsb* MO and *p53* MO^ATG^ rescued the small eye phenotype caused by injection of *nhsb* MO alone. (Fig. 7A). Splice blocking efficiency and knockdown of *nhsb* was confirmed by RT-PCR (Fig. S6) and phenotypes quantified (Fig. S7). Interestingly, co-injection of *nhsb* MO and *p53* MO^ATG^ led to a reduced expression of *Δ113p53* and *p21* transcripts, compared to *nhsb* MO alone (Fig. 6B). Our findings indicate that the upregulation of *Δ113p53* and *p21* in *nhsb-*depleted retina is dependent on full length p53, consistent with the notion that *Δ113p53* and *p21* are p53 target genes (Chen et al. 2009). Overall this result suggests that the proliferation restriction enforced on RPCs was alleviated when levels of *Δ113p53* and *p21* were reduced.

**Fig. 7.**
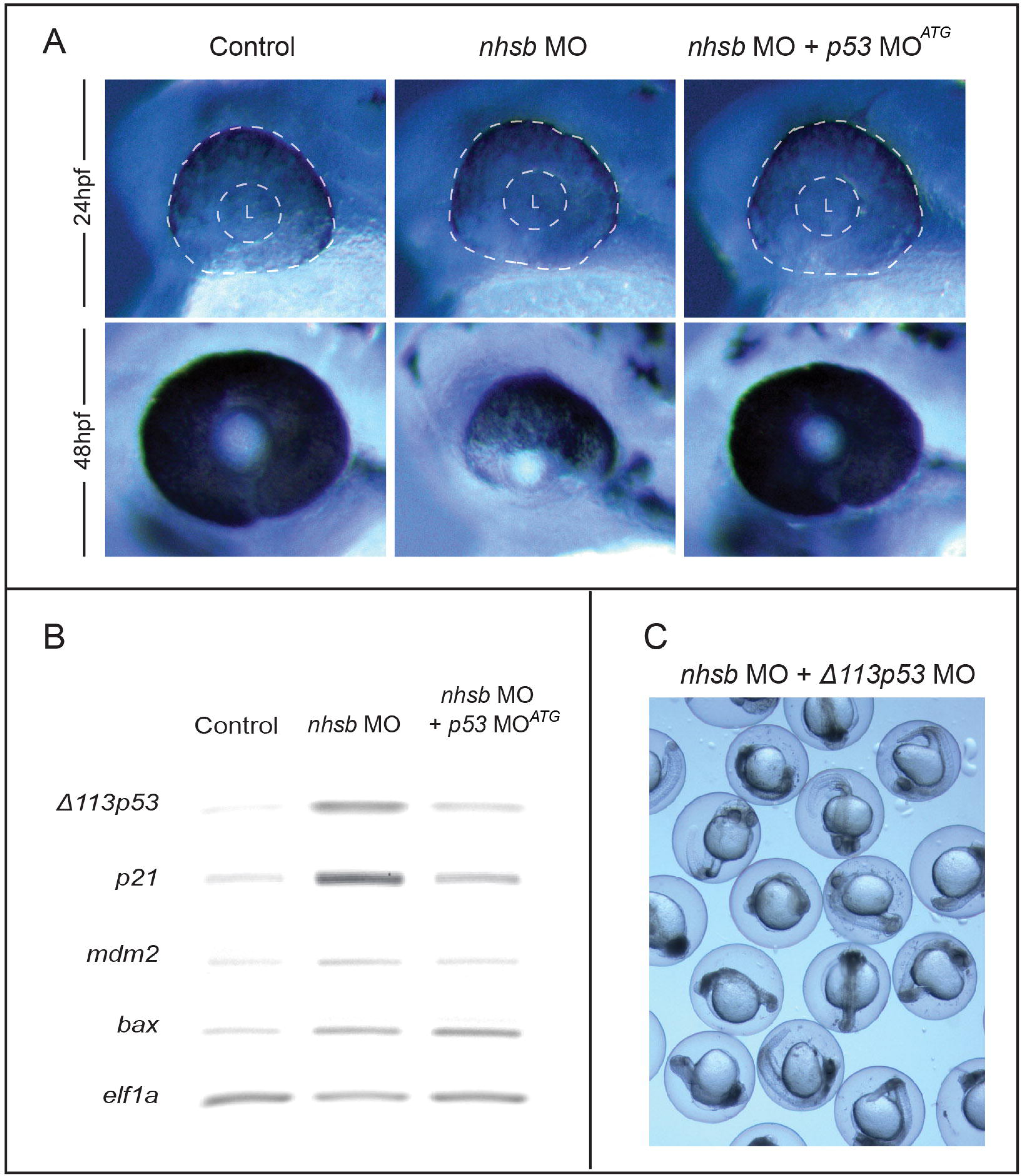
Knockdown of p53 rescues nhsb morphant phenotype and Δ113p53 levels. (A) Stereomicroscope images of zebrafish retina at 24 hpf and 48 hpf injected with nhsb MO alone or in combination with p53 MOATG (L = lens). Retinal size was not dramatically differ­ent in any condition at 24 hpf. However, injection of p53 MOATG targeting full-length p53 transcripts rescued retinal size in nhsb morphants. (B) Semi-quantitative RT-PCR examination of p53 target genes. Note that injection of p53 MOATG inhibited the upregulation of Δ113p53 and p21 transcripts caused by nhsb-depletion. (C) Injection of Δ113p53 MO that specifically targets the Δ113p53 variant and not full-length p53 led to elevated cell death.

It was previously reported that *Δ113p53* antagonizes the apoptotic activities of *p53* (Chen and Peng 2005; Chen et al. 2009), thereby promoting cell survival. To determine whether expression of *Δ113p53* variant is required for cell survival in *nhsb* morphants, we designed a second morpholino targeting the splicing site of the *Δ113p53* transcript at the exon ES - intron 4 splice junction (*Δ113p53* splMO). The activity of *Δ113p53* splMO is predicted to affect splicing of the *Δ113p53* transcript, but not full-length *p53* transcripts. Injection of either *p53* MO^ATG^ or *Δ113p53* splMO alone had no discernable effect on retinal development (data not shown). However, embryos co-injected with *nhsb* MO and *Δ113p53* splMO resulted in embryonic death, indicative of an increase in *p53*-mediated apoptosis when *Δ113p53* transcripts are absent.

Taken together, these results demonstrate that the upregulation of *Δ113p53* protects retinal progenitors from undergoing cell death in *nhsb* knockdown embryos, while imposing a proliferation restriction in a manner that likely involves other p53 target genes, such as the cell cycle inhibitor *p21*. Our data supports a model where *nhsb* is a negative regulator of *Δ113p53* expression and exerts its function in development though regulation of the p53 pathway.

## Discussion

The vertebrate retina develops from a small pool of retinal progenitor cells (RPCs) comprising the optic cup. Just prior to the onset of retinal neurogenesis, RPCs undergo cycles of rapid self-replication to expand the tissue (Agathocleous and Harris 2009; He et al. 2012). The proliferative capacity of RPC’s are diminished when key transcription factors like *Rx1*, *Pax6, Chx10,* and *Sox2* are inhibited, whereas activation of these transcription factors often lads to increased mitosis and a larger retina (Marquardt et al. 2001; Green et al. 2003; Philips et al. 2005; Van Raay et al. 2005; Taranova et al. 2006; Zaghloul and Moody 2007; Oron-Karni et al. 2008). Temporal expression of bHLH proteins like neurogenin 2 (Ngn2), atonal homolog 5 (ATH5) and NeuroD trigger cell cycle exit to initiate specification and differentiation in a highly coordinated manner (Agathocleous and Harris 2009). Extrinsic signals from the retina microenvironment also affects division and differentiation of RPCs. Ligands like Sonic Hedgehog (Shh) promotes rapid proliferation of RPCs by shortening cell cycle length through the activation of G1 and G2 phase cyclins (Wang et al. 2005; Locker et al. 2006; Agathocleous et al. 2007). It has been proposed that Shh stimulates the transition of early RPC from slow cycling to faster cycling in advance of terminal differentiation (Locker et al. 2006; Agathocleous et al. 2007). Our results demonstrate that knockdown of Nhs in zebrafish and *Xenopus* leads to a reduction in proliferation of RPCs before the onset of differentiation. We propose that loss of Nhs function leads to an increase in cell cycle length that may be due to the elevated levels of cyclin-dependent kinase inhibitors, such as p21, as a result of p53 pathway activation. Our results show that Nhs-depletion does not inhibit, but rather delays the differentiation of RPCs. Although differentiation proceeds, it remains unclear whether loss of Nhs function affects the differentiation of RPCs into all possible cell types in the retina. Nevertheless, our findings indicate Nhs regulates overall retinal size by promoting the faster cycling of RPCs associated with the expansion of the RPC population in advance of terminal differentiation.

Although it is clear that the p53 tumor suppressor plays important roles in cancer progression, a role for p53 (or any or the p53 isoforms) in embryonic development or development of the nervous system is less clear. p53 acts primarily as a transcription factor, although it has been shown to have a cytoplasmic roles (Vousden 2006; Endo et al. 2008). In response to DNA damage or other cellular stresses, there are two main outcomes of p53 activation, namely cell cycle arrest or apoptosis. In the nervous system, p53 plays a key role, along with other p53 family members, p63 and p73, in regulating survival and apoptosis in the developing and adult nervous system (Miller et al. 2000; Jacobs et al. 2004). p53 becomes a major pro-apoptotic proteins eliciting naturally occurring cell death in peripheral neurons (Aloyz et al. 1998; Jacobs et al. 2005)and neural progenitor cells (Dugani et al. 2009; Fatt et al. 2014). Consistent with this, mice deficient in DNA ligase 4 (Lig4) or XRCC4, two components of the nonhomologous end-joining (NHEJ) pathway that repairs double-stranded DNA breaks, exhibit neuronal apoptosis that has been shown to be p53-dependent (Frank et al. 1998; Gao et al. 1998). A screen of zebrafish mutants harboring mutations various categories of cell essential genes, such as DNA replication, transcription, telomere maintenance, ribosome biogenesis, splicing, chaperoning, endocytosis, and cellular transport, all share an upregulation of p53 and excessive neural apoptosis (Danilova et al. 2010). Activation of the p53 pathway then is likely responsible for a wide range of defects seen in human congenital disorders (Mendrysa et al. 2011).

Few studies, however, have investigated the role of p53 isoforms, such as Δ113p53/Δ113p53, in embryonic development. The zebrafish protein Δ113p53, and its human counterpart, Δ133p53, are N-terminally truncated forms of p53 that are missing the transactivation domain, and MDM2-interacting motif, with a partial deletion of the DNA-binding domain (Bourdon et al. 2005; Chen et al. 2005, 2009). Transcriptional expression of Δ113p53/Δ133p53 is initiated by an alternative p53 promoter in intron 4 that is completely dependent on full length p53 protein (Bourdon et al. 2005; Chen et al. 2009; Aoubala et al. 2011). Δ113p53/Δ133p53 is strongly induced by DNA double strand breaks, where it antagonizes p53-mediated apoptosis(Gong et al. 2015). Previous work had shown that Δ113p53 does not act in a dominant-negative fashion, but by differentially modulating p53 target gene activation to protect cells from apoptosis (Chen et al. 2009; Ou et al. 2014). In zebrafish, *Δ113p53* was identified in the analysis of a zebrafish mutant *def,* a pan-endodermal-enriched factor that is essential for the expansive growth of digestive organs (Chen et al. 2005). In *def* null mutants, *Δ113p53* transcripts were upregulated and induced the expression of p53-responsive genes to trigger arrest of cell cycle, but not apoptosis, leading to reduced organ growth (Chen et al. 2005, 2009). We identified a similar role for *nhsb* in expansion growth of the retina. In the absence of *nhsb*, *Δ113p53* expression increased, which concomitant increase of the p53 target gene *p21*. Increased levels of *p21* enforced a proliferation restriction on RPCs, while the pro-apoptotic gene *bax* was not increased in hypoplastic retinas. Our findings are in agreement with the previously reported role for *Δ113p53* as regulator of *p21*, but not *bax* (Chen et al. 2005, 2009). These results suggest that i) Nhs is required in retinal progenitor cells to keep levels of *Δ113p53* low, and that ii) loss of *nhs* causes a small retina due to cell cycle arrest, but not apoptosis. Indeed, when *Δ113p53* were depleted, loss of *nhsb* led to an increase in cell death.

How does Nhs regulate the *Δ113p53* transcript levels? Our data suggest that Nhs is a negative regulator of *Δ113p53* expression. Nhs could regulate *Δ113p53* levels directly or indirectly through modulation of the p53 pathway. Our experiments indicate full length p53 is required for the micropthalmia and *Δ113p53* induction caused by *nhsb* deficiency. However, we show that transcript levels of full-length *p53* are unchanged in *nhsb* morphants, suggesting that Nhs does not regulate p53 at the level of transcription. One possibility is that Nhs regulates the steady state levels of p53 protein, and in the absence of Nhs, p53 levels would accumulate and elevate expression of *Δ113p53* and *p21*. Alternatively, Nhs could limit the transcriptional activity of p53 without affecting protein levels. It was recently reported that Def is a nucleolar protein that mediates p53 protein degradation through a proteasome-independent pathway (Tao et al. 2013). Future studies will be aimed at uncovering the mechanism of how Nhs modulates the p53 pathway.

Finally, our study uncovers a developmental role for Nhs that provides an underlying mechanism for pathology associated with Nance-Horan Syndrome (NHS). Affected males have severe bilateral congenital dense nuclear cataracts that lead to profound vision loss and usually require surgery at an early age (Burdon et al. 2003; Huang et al. 2007; Sharma et al. 2008; Coccia et al. 2009). Despite cataract surgery during infancy, many NHS patients continue to exhibit visual impairment that may be caused by an underlying retinopathy (Mathys et al. 2007; Ding et al. 2009). Our findings that Nhs is essential for proper retinal development may clarify why NHS patients continue to exhibit ocular abnormalities and vision loss, even when cataracts have been corrected by surgery. NHS is caused predominantly by truncating (nonsense) mutations in the *NHS* gene that have been found in each of the 8 exons. There is no apparent clinical difference between mutations in C-terminal exons compared to N-terminal exons (Tug et al. 2013). Previous reports have suggested that humans have at least three isoforms of NHS transcripts generated by the use of alternative transcriptional start sites, arising from exon1, exon1a, and exon1b (see Fig 1B). Transcripts encoded from exon1 is known to be important in the pathogenesis of NHS, because patient nonsense mutations identified in exon1 are only predicted to affect this isoform (Burdon et al. 2003; Brooks et al. 2004; Huang et al. 2006; Hong et al. 2014). This exon1 is evolutionarily conserved in vertebrates. Our results presented here using CRISPR genome editing with guide RNAs directed against exon1 of the zebrafish *nhsb* gene showed a similar micropthalmia compared with morpholino oligonucleotide-mediated knockdown. These results are consistent with the human molecular genetics demonstrating that NHS isoforms encoded from exon1 are relevant to the pathogenesis of Nance-Horan Syndrome.

## Materials and Methods

### Zebrafish and *Xenopus tropicalis* Husbandry

Zebrafish (Danio rerio) were maintained according to standard procedures and staged as previously described (Kimmel et al. 1995). The *nhsa^fh298^* mutant line carries a truncating nonsense mutation at position Q217* and was generated by TILLING (Draper et al. 2004). The *neurod:EGFP* transgenic line is registered as *TgBAC(neurod1:EGFP)nl1Tg* at The Zebrafish International Resource Center (ZIRC)(Obholzer et al. 2008). *Xenopus tropicalis* embryos were obtained and cultured using standard methods (Sive et al. 2000).

### Phlyogenetic analysis

Coding DNA sequences were gathered from, Ensembl (http://www.ensembl.org). Splice variants were checked by Mouse Genome Informatics (http://www.informatics.jax.org/), Zfin (http://zfin.org/), Xenbase (http://www.xenbase.org) and FlyBase (http://flybase.org/). *Xenopus* and zebrafish genes were manually curated using UCSC genome browser (https://genome.ucsc.edu/), Xenbase and Zfin. The following genes were used in our analysis:

*<u>human</u>:* NHS (ENSG00000188158), NHSL1 (ENSG00000135540), NHSL2 (ENSG00000204131). *<u>Mus musculus</u>:* NHS (ENSMUSG00000059493), NHSL1 (ENSMUSG00000039835), NHSL2 (ENSMUSG00000079481). *<u>Xenopus tropicalis</u>:* NHS (ENSXETG00000001112), NHSL1 (ENSXETG00000006237), NHSL2 (ENSXETG00000022580). *<u>Danio rerio</u>*: *nhsa* (ENSDARG00000070227), *nhsb* (ENSDARG00000079977), *nhsl1a* (ENSDARG00000054537), *nhsl1b* (ENSDARG00000042627), *nhsl2a* (ENSDARG00000089066), *nhsl2b* (ENSDARG00000090164). *<u>Drosophila melanogaster</u>*: *gukh* (FBgn0026239). Above genes were entered into ClustlW2 Phylogeny (http://www.ebi.ac.uk/) and aligned by multiple sequence alignment. A Newick output was generated using the Neighbour-joining clustering method. Cladogram of Coding DNA was composed using Genious 7.0.6 software.

### Whole-mount in situ hybridization

RNA in situ hybridization was carried out as previously described (Walsh et al., 2011). The gene specific probe fragments to the 3’UTR region of *nhs* genes were obtained by PCR (see Table S1 for primer pairs) from cDNA of 24 hpf zebrafish wild type embryos or early-stage *X. tropicalis* embryos. PCR products were cloned into pCR4-Blunt, sequenced. Sense and antisense riboprobes were transcribed from linearized plasmids and labeled with digoxigenin (DIG RNA-labelling reagents, Roche Biochemicals). Primers used to amplify zebrafish *nhsb* were 5’ tgatctaccttttctcccatgccatt 3’ and 5’ tctcacacaccacagaggctcca 3’. Primers used to amplify *Xenopus tropicalis nhs* were 5’ tgccagcccacattgattatag 3’ and 5’ ccttaaaaagcaggccacagtt 3’.

### Morpholino injections

Antisense MOs were injected at the 1-cell stage in zebrafish embryos and into one cell of a two-cell stage *Xenopus* embryo. Microinjections of antisense MOs were carried out using an Eppendorf microinjector and Zeiss stereoscope, as described (Dickinson and Sive, 2009). Morpholinos were supplied by Gene Tools and were as follows: *zebrafish nhsb*: MO1 E2I2 5’CCAGAAGACCCATCAGTACCTCTGT3’, 4 ng, MO2 I2E3 5’ CTGCTGCTGGTCTGTGAAGCAAACA 3’, 3.5 ng, *Xenopus tropicalis nhs:* MO E3I3 5’ GTGTGTAATATATCTTACCTCTCCG 3’, 8.5 ng. The *p53* MO^ATG^ targets the translation start site of full-length p53 as described (Robu et al. 2007): 5’GCGCCATTGCTTTGCAAGAATTG 3’, 3 ng. A morpholino that targets the splice junction from exon ES to intron 4 of p53 and disrupts splicing of *Δ113p53* isoform while leaving full-length p53 intact: splMO 5’ TGTCTTTTCAAATGTCTTACCCTCC 3’, 3 ng.

### CRISPR Design and injection

We designed guide RNA (gRNA) sequence directed against exon1 of *nhsb* (5’ GGTCCGGGATAGAGCCACAT 3’). For making gRNA, the guide sequence template was generated by annealing and filling-in two oligonucleotides: guide oligo (5’ aattaatacgactcactataGGTCCGGGATAGAGCCACATgttttagagctagaaatagc3’) and scaffold template (5’GATCCGCACCGACTCGGTGCCACTTTTTCAAGTTGATAACGGAC TAGCCTTATTTTAACTTGCTATTTCTAGCTCTAAAAC3’). The resulting gRNA template that contains a T7 promoter, was purified using DNA clean columns (ZymoResearch). *gRNA* was generated by in vitro transcription using T7 RNA polymerase (Megascript T7 Kit, Invitrogen). After transcription, gRNA was column purified (ZymoResearch). For making nls-zCas9-nls RNA, the template DNA (pT3TS-nls-zCas9-nls; kind gift of Wenbiao Chen (Addgene plasmid # 46757) was linearized by XbaI, phenol-chloroform extracted followed by ethanol precipitation. Capped nls-zCas9-nls mRNA was synthesized using mMessage T3 kit (Invitrogen) and purified using Illustra Probe-quant columns (GE Healthcare). The mix of gRNA (100ng/μL) and nls-zCas9-nls mRNA (300 ng/μL) was injected into one-cell stage embryos.

### T7 Endonuclease I Assay

Genomic DNA was prepared from whole embryos at 24 hpf. A short stretch of genomic region (∼735 bp) flanking the target site was PCR amplified from genomic DNA. PCR product was denatured and slowly allowed to re-anneal to encourage heteroduplex formation. The re-annealing procedure consisted of a 5 minute denaturation step, followed by cooling to 85°C at −2°C per second and further to 25°C at a rate of −0.1°C per second. The re-annealed amplicon was then digested with 20 units of T7 endonuclease I (New England Biolabs) at 37°C for 15 minutes. The reaction was stopped by adding 1 μL of 0.5M EDTA. The sample was run through a 2% agarose gel and visualized by ethidium bromide staining.

### Reverse Transcription-Polymerase Chain Reaction (RT-PCR)

Splice-blocking MOs were validated by RT-PCR. Primers used to validate are listed in Supplementary Table 1. Briefly, total RNA was isolated from whole embryos using an RNeasy kit (Qiagen) and genomic contamination was eliminated by on-column DNA digestion (Qiagen). First Strand cDNA was synthesized using the Protoscript M-MuLV First-Strand cDNA Synthesis Kit (New England Biolabs). Equal concentrations of RNA (1ug) from all developmental timepoints were used in the cDNA synthesis reaction. Concentration of cDNA across developmental timepoints was validated using primers specific to the ornithine decarboxylase (ODC) gene or *elf1a*. RNA extracted from retinas were used to validate array data. For semiquantitative RT-PCR, the amount of template and the number of PCR cycles were optimized to ensure that the reactions were in the linear range of amplification. The primer pairs and detailed PCR conditions used to amplify each of these genes are listed in Table S1.

### Affymetrix Array

We dissected 40 eyes each from wildtype or *nhsb* MO2-injected embryos at 48hpf. This experiment was performed in duplicate. Dissections were done in Half Ringers Solution (2.2mM CaCl2, 5.6mM KCl, 154 mM NaCl, 2.4mM NaHCO3, 2 mM Tris-Cl, adjust to pH 7.4) on ice. Total RNA was extracted from pooled tissues using RNeasy kit (Qiagen) and treated with DNAse I. We used 200ng input total RNA for cRNA amplification using the Affymetrix One Cycle kit. Amplicons were fragmented, hybridized to GeneChip Zebrafish Genome Arrays and scanned (Affymetrix). Quality control of total RNA, amplified RNA, and fragmented RNA was checked using Experion (Bio-Rad). GeneChip arrays were scanned on an Affymetrix Scanner 3000 7G scanner. Data was analyzed by Affymetrix Expression Console software. Raw intensity data was exported and filtered with a lower cut off value of 2 to assure that probe intensities data were generated within the dynamic range of fluorescence. Data was normalized by Robust Multi-array Average (RMA) analysis to account for a possible probe effect, and uploaded into TMEV for Significance Analysis of Microarrays (SAM) analysis. The resulting 1,212 significant genes were entered into DAVID for further gene ontology analysis.

### Immunostaining and Imaging

Embryos were fixed in 4% PFA overnight at 4°C, incubated in 30% sucrose and saturated in OCT (Tissue-Tec), frozen on cryostat stages prior to storage at −20°C. Coronal sections of 10uM were cut on a cryostat and mounted to Colorfrost Plus, microscope slides (Fisher). Immunostaining was performed on sections with the following antibodies: anti-phosphohistone 3 (pH3) (1:1000 Cell Signaling); anti-GFP (1:2000, Torrey Pines); cleaved caspase 3 (1:500) (Cell Signaling). Primary antibodies were detected using secondary antibodies: goat anti-mouse Alexa 564 or goat anti-rabbit Alexa 488 (Life Technologies). Nuclei were labeled with Dapi. Sections were mounted in 100% glycerol and coverslipped. Confocal images of labeled cryostat sections or whole-mount embryos were obtained on a Carl Zeiss Spinning Disk Laser Confocal Observer Z1 (VCU) using either 20X or 63X objectives.

## References

Accogli A, Traverso M, Madia F, et al (2017) A novel Xp22.13 microdeletion in Nance-Horan syndrome. Birth Defects Res 109:866–868. doi: 10.1002/bdr2.1032

Agathocleous M, Harris WA (2009) From Progenitors to Differentiated Cells in the Vertebrate Retina. Annu Rev Cell Dev Biol 25:45–69. doi: 10.1146/annurev.cellbio.042308.113259

Agathocleous M, Locker M, Harris WA, Perron M (2007) A General Role of Hedgehog in the Regulation of Proliferation. Cell Cycle 6:156–159. doi: 10.4161/cc.6.2.3745

Aloyz RS, Bamji SX, Pozniak CD, et al (1998) p53 is essential for developmental neuron death as regulated by the TrkA and p75 neurotrophin receptors. J Cell Biol 143:1691–703

Aoubala M, Murray-Zmijewski F, Khoury MP, et al (2011) p53 directly transactivates Δ133p53α, regulating cell fate outcome in response to DNA damage. Cell Death Differ 18:248–258. doi: 10.1038/cdd.2010.91

Boije H, MacDonald RB, Harris WA (2014) Reconciling competence and transcriptional hierarchies with stochasticity in retinal lineages. Curr Opin Neurobiol 27:68–74. doi: 10.1016/j.conb.2014.02.014

Bourdon J-C, Fernandes K, Murray-Zmijewski F, et al (2005) p53 isoforms can regulate p53 transcriptional activity. Genes Dev 19:2122–2137. doi: 10.1101/gad.1339905

Brooks SP, Coccia M, Tang HR, et al (2010a) The Nance-Horan syndrome protein encodes a functional WAVE homology domain (WHD) and is important for co-ordinating actin remodelling and maintaining cell morphology. Hum Mol Genet 19:2421–2432. doi: 10.1093/hmg/ddq125

Brooks SP, Coccia M, Tang HR, et al (2010b) The Nance-Horan syndrome protein encodes a functional WAVE homology domain (WHD) and is important for co-ordinating actin remodelling and maintaining cell morphology. Hum Mol Genet 19:2421–32. doi: 10.1093/hmg/ddq125

Brooks SP, Ebenezer ND, Poopalasundaram S, et al (2004) Identification of the gene for Nance-Horan syndrome (NHS). J Med Genet 41:768–771. doi: 10.1136/jmg.2004.022517

Burdon KP, McKay JD, Sale MM, et al (2003) Mutations in a novel gene, NHS, cause the pleiotropic effects of Nance-Horan syndrome, including severe congenital cataract, dental anomalies, and mental retardation. Am J Hum Genet 73:1120–1130. doi: 10.1086/379381

Cazzalini O, Scovassi AI, Savio M, et al (2010) Multiple roles of the cell cycle inhibitor p21CDKN1A in the DNA damage response. Mutat Res Mutat Res 704:12–20. doi: 10.1016/j.mrrev.2010.01.009

Chen J, Ng SM, Chang C, et al (2009) p53 isoform delta113p53 is a p53 target gene that antagonizes p53 apoptotic activity via BclxL activation in zebrafish. Genes Dev 23:278–90. doi: 10.1101/gad.1761609

Chen J, Peng J (2009) p53 Isoform Δ113p53 in Zebrafish. Zebrafish 6:389–395. doi: 10.1089/zeb.2009.0598

Chen J, Peng J (2005) Δ113p53 / Δ133p53 : survival and integrity. 6:6–7

Chen J, Ruan H, Ng SM, et al (2005) Loss of function of def selectively up-regulates Delta113p53 expression to arrest expansion growth of digestive organs in zebrafish. Genes Dev 19:2900–11. doi: 10.1101/gad.1366405

Chograni M, Rejeb I, Jemaa L Ben, et al (2011) The first missense mutation of NHS gene in a Tunisian family with clinical features of NHS syndrome including cardiac anomaly. Eur J Hum Genet 19:851–6. doi: 10.1038/ejhg.2011.52

Coccia M, Brooks SP, Webb TR, et al (2009) X-linked cataract and Nance-Horan syndrome are allelic disorders. Hum Mol Genet 18:2643–55. doi: 10.1093/hmg/ddp206

Danilova N, Kumagai A, Lin J (2010) p53 Upregulation Is a Frequent Response to Deficiency of Cell-Essential Genes. PLoS One 5:e15938. doi: 10.1371/journal.pone.0015938

Ding X, Patel M, Herzlich AA, et al (2009) Ophthalmic pathology of Nance-Horan syndrome: case report and review of the literature. Ophthalmic Genet 30:127–35. doi: 10.1080/13816810902822021

Donehower LA, Harvey M, Slagle BL, et al (1992) Mice deficient for p53 are developmentally normal but susceptible to spontaneous tumours. Nature 356:215–21. doi: 10.1038/356215a0

Draper BW, McCallum CM, Stout JL, et al (2004) A high-throughput method for identifying N-ethyl-N-nitrosourea (ENU)-induced point mutations in zebrafish. Methods Cell Biol 77:91–112

Dugani CB, Paquin A, Fujitani M, et al (2009) p63 antagonizes p53 to promote the survival of embryonic neural precursor cells. J Neurosci 29:6710–21. doi: 10.1523/JNEUROSCI.5878-08.2009

Endo Y, Sugiyama A, Li S-A, et al (2008) Regulation of clathrin-mediated endocytosis by p53. Genes to Cells 13:375–386. doi: 10.1111/j.1365-2443.2008.01172.x

Fadool JM, Dowling JE (2008) Zebrafish: a model system for the study of eye genetics. Prog Retin Eye Res 27:89–110. doi: 10.1016/j.preteyeres.2007.08.002

Fatt MP, Cancino GI, Miller FD, Kaplan DR (2014) p63 and p73 coordinate p53 function to determine the balance between survival, cell death, and senescence in adult neural precursor cells. Cell Death Differ 21:1546–1559. doi: 10.1038/cdd.2014.61

Florijn RJ, Loves W, Maillette de Buy Wenniger-Prick LJJM, et al (2006) New mutations in the NHS gene in Nance–Horan Syndrome families from the Netherlands. Eur J Hum Genet 14:986–990. doi: 10.1038/sj.ejhg.5201671

Frank KM, Sekiguchi JM, Seidl KJ, et al (1998) Late embryonic lethality and impaired V(D)J recombination in mice lacking DNA ligase IV. Nature 396:173–177. doi: 10.1038/24172

Gao Y, Sun Y, Frank KM, et al (1998) A critical role for DNA end-joining proteins in both lymphogenesis and neurogenesis. Cell 95:891–902

Gómez-Laguna L, Martínez-Herrera A, Reyes-de la Rosa A del P, et al (2017) Nance–Horan syndrome in females due to a balanced X;1 translocation that disrupts the *NHS* gene: Familial case report and review of the literature. Ophthalmic Genet 1–7. doi: 10.1080/13816810.2017.1363245

Gong L, Gong H, Pan X, et al (2015) p53 isoform Δ113p53/Δ133p53 promotes DNA double-strand break repair to protect cell from death and senescence in response to DNA damage. Cell Res 25:351–369. doi: 10.1038/cr.2015.22

Green ES, Stubbs JL, Levine EM (2003) Genetic rescue of cell number in a mouse model of microphthalmia: interactions between Chx10 and G1-phase cell cycle regulators. Development 130:539–52

Harris SL, Levine AJ (2005) The p53 pathway: positive and negative feedback loops. Oncogene 24:2899–2908. doi: 10.1038/sj.onc.1208615

He J, Zhang G, Almeida AD, et al (2012) How variable clones build an invariant retina. Neuron 75:786–98. doi: 10.1016/j.neuron.2012.06.033

Hindley C, Philpott A (2012) Co-ordination of cell cycle and differentiation in the developing nervous system. Biochem J 444:375–382. doi: 10.1042/BJ20112040

Hong N, Chen Y, Xie C, et al (2014) Identification of a novel mutation in a Chinese family with Nance-Horan syndrome by whole exome sequencing. J Zhejiang Univ B 15:727–734. doi: 10.1631/jzus.B1300321

Hu M, Easter SS (1999) Retinal neurogenesis: the formation of the initial central patch of postmitotic cells. Dev Biol 207:309–21. doi: 10.1006/dbio.1998.9031

Huang KM, Wu J, Brooks SP, et al (2007) Identification of three novel NHS mutations in families with Nance-Horan syndrome. Mol Vis 13:470–474. doi: v13/a49 [pii]

Huang KM, Wu J, Duncan MK, et al (2006) Xcat, a novel mouse model for Nance-Horan syndrome inhibits expression of the cytoplasmic-targeted Nhs1 isoform. Hum Mol Genet 15:319–327. doi: 10.1093/hmg/ddi449

Jacobs WB, Govoni G, Ho D, et al (2005) P63 Is an Essential Proapoptotic Protein during Neural Development. Neuron 48:743–756. doi: 10.1016/j.neuron.2005.10.027

Jacobs WB, Walsh GS, Miller FD (2004) Neuronal survival and p73/p63/p53: a family affair. Neuroscientist 10:443–55. doi: 10.1177/1073858404263456

Jao L, Wente SR, Chen W (2013) Efficient multiplex biallelic zebra fish genome editing using a CRISPR nuclease system. 110:1–6. doi: 10.1073/pnas.1308335110

Joruiz SM, Bourdon J-C (2016) p53 Isoforms: Key Regulators of the Cell Fate Decision. Cold Spring Harb Perspect Med 6:a026039. doi: 10.1101/cshperspect.a026039

Joseph B, Hermanson O (2010) Molecular control of brain size: Regulators of neural stem cell life, death and beyond. Exp Cell Res 316:1415–1421. doi: 10.1016/j.yexcr.2010.03.012

Katoh M, Katoh M (2004) Identification and characterization of human GUKH2 gene in silico. Int J Oncol 24:1033–8

Kimmel CB, Ballard WW, Kimmel SR, et al (1995) Stages of embryonic development of the zebrafish. Dev Dyn 203:253–310. doi: 10.1002/aja.1002030302

Li A, Li B, Wu L, et al (2015) Identification of a Novel *NHS* Mutation in a Chinese Family with Nance-Horan Syndrome. Curr Eye Res 40:434–438. doi: 10.3109/02713683.2014.959606

Li Z, Hu M, Ochocinska MJ, et al (2000) Modulation of cell proliferation in the embryonic retina of zebrafish (Danio rerio). Dev Dyn 219:391–401. doi: 10.1002/1097-0177(2000)9999:9999<::AID-DVDY1063>3.0.CO;2-G

Locker M, Agathocleous M, Amato MA, et al (2006) Hedgehog signaling and the retina: insights into the mechanisms controlling the proliferative properties of neural precursors. Genes Dev 20:3036–3048. doi: 10.1101/gad.391106

Marcel V, Vijayakumar V, Fernández-Cuesta L, et al (2010) p53 regulates the transcription of its Δ133p53 isoform through specific response elements contained within the TP53 P2 internal promoter. Oncogene 29:2691–2700. doi: 10.1038/onc.2010.26

Marine J-C, Francoz S, Maetens M, et al (2006) Keeping p53 in check: essential and synergistic functions of Mdm2 and Mdm4. Cell Death Differ 13:927–934. doi: 10.1038/sj.cdd.4401912

Marquardt T, Ashery-Padan R, Andrejewski N, et al (2001) Pax6 is required for the multipotent state of retinal progenitor cells. Cell 105:43–55

Mathys R, Deconinck H, Keymolen K, et al (2007) Severe visual impairment and retinal changes in a boy with a deletion of the gene for Nance-Horan syndrome. Bull Soc Belge Ophtalmol 49–53

McCubrey JA, Lertpiriyapong K, Fitzgerald TL, et al (2017) Roles of TP53 in determining therapeutic sensitivity, growth, cellular senescence, invasion and metastasis. Adv Biol Regul 63:32–48. doi: 10.1016/j.jbior.2016.10.001

Mendrysa SM, Ghassemifar S, Malek R (2011) p53 in the CNS: Perspectives on Development, Stem Cells, and Cancer. Genes Cancer 2:431–42. doi: 10.1177/1947601911409736

Miller FD, Pozniak CD, Walsh GS (2000) Neuronal life and death: an essential role for the p53 family. Cell Death Differ 7:880–8. doi: 10.1038/sj.cdd.4400736

Obholzer N, Wolfson S, Trapani JG, et al (2008) Vesicular Glutamate Transporter 3 Is Required for Synaptic Transmission in Zebrafish Hair Cells. J Neurosci 28:2110–2118. doi: 10.1523/JNEUROSCI.5230-07.2008

Oron-Karni V, Farhy C, Elgart M, et al (2008) Dual requirement for Pax6 in retinal progenitor cells. Development 135:4037–4047. doi: 10.1242/dev.028308

Ou Z, Yin L, Chang C, et al (2014) Protein Interaction Between p53 and Δ113p53 Is Required for the Anti-Apoptotic Function of Δ113p53. J Genet Genomics 41:53–62. doi: 10.1016/j.jgg.2014.01.001

Philips GT, Stair CN, Young Lee H, et al (2005) Precocious retinal neurons: Pax6 controls timing of differentiation and determination of cell type. Dev Biol 279:308–321. doi: 10.1016/j.ydbio.2004.12.018

Postlethwait JH, Woods IG, Ngo-Hazelett P, et al (2000) Zebrafish comparative genomics and the origins of vertebrate chromosomes. Genome Res 10:1890–902. doi: 10.1101/GR.164800

Postlethwait JH, Yan Y-L, Gates MA, et al (1998) Vertebrate genome evolution and the zebrafish gene map. Nat Genet 18:345–349. doi: 10.1038/ng0498-345

Ramprasad VL, Thool A, Murugan S, et al (2005) Truncating Mutation in the *NHS* Gene: Phenotypic Heterogeneity of Nance-Horan Syndrome in an Asian Indian Family. Investig Opthalmology Vis Sci 46:17. doi: 10.1167/iovs.04-0477

Robu ME, Larson JD, Nasevicius A, et al (2007) P53 Activation By Knockdown Technologies. PLoS Genet 3:787–801. doi: 10.1371/journal.pgen.0030078

Sah VP, Attardi LD, Mulligan GJ, et al (1995) A subset of p53-deficient embryos exhibit exencephaly. Nat Genet 10:175–80. doi: 10.1038/ng0695-175

Sharma S, Ang SL, Shaw M, et al (2006) Nance-Horan syndrome protein, NHS, associates with epithelial cell junctions. Hum Mol Genet 15:1972–1983. doi: 10.1093/hmg/ddl120

Sharma S, Burdon KP, Dave A, et al (2008) Novel causative mutations in patients with Nance-Horan syndrome and altered localization of the mutant NHS-A protein isoform. 1856–1864

Shoshany N, Avni I, Morad Y, et al (2017) *NHS* Gene Mutations in Ashkenazi Jewish Families with Nance–Horan Syndrome. Curr Eye Res 42:1240–1244. doi: 10.1080/02713683.2017.1304560

Sive H, Grainger R, Harland R (2000) Early Development of Xenopus laevis: A Laboratory Manual. Cold Spring Harbor Laboratory Press

Tao T, Shi H, Guan Y, et al (2013) Def defines a conserved nucleolar pathway that leads p53 to proteasome-independent degradation. Cell Res 23:620–634. doi: 10.1038/cr.2013.16

Taranova O V., Magness ST, Fagan BM, et al (2006) SOX2 is a dose-dependent regulator of retinal neural progenitor competence. Genes Dev 20:1187–1202. doi: 10.1101/gad.1407906

Thomas JL, Ochocinska MJ, Hitchcock PF, Thummel R (2012) Using the Tg(nrd:egfp)/albino zebrafish line to characterize in vivo expression of neurod. PLoS One 7:e29128. doi: 10.1371/journal.pone.0029128

Tian Q, Li Y, Kousar R, et al (2017) A novel NHS mutation causes Nance-Horan Syndrome in a Chinese family. BMC Med Genet 18:2. doi: 10.1186/s12881-016-0360-9

Tug E, Dilek NF, Javadiyan S, et al (2013) A Turkish family with Nance-Horan Syndrome due to a novel mutation. Gene 525:141–5. doi: 10.1016/j.gene.2013.03.094

Van Nostrand JL, Brady CA, Jung H, et al (2014) Inappropriate p53 activation during development induces features of CHARGE syndrome. Nature 514:228–32. doi: 10.1038/nature13585

Van Raay TJ, Moore KB, Iordanova I, et al (2005) Frizzled 5 Signaling Governs the Neural Potential of Progenitors in the Developing Xenopus Retina. Neuron 46:23–36. doi: 10.1016/j.neuron.2005.02.023

Vogelstein B, Lane D, Levine AJ (2000) Surfing the p53 network. 408:

Vousden KH (2006) Outcomes of p53 activation - spoilt for choice. J Cell Sci 119:5015–5020. doi: 10.1242/jcs.03293

Vousden KH, Lane DP (2007) p53 in health and disease. Nat Rev Mol Cell Biol 8:275–83. doi: 10.1038/nrm2147

Wang Y, Dakubo GD, Thurig S, et al (2005) Retinal ganglion cell-derived sonic hedgehog locally controls proliferation and the timing of RGC development in the embryonic mouse retina. Development 132:5103–5113. doi: 10.1242/dev.02096

Willardsen MI, Link BA (2011) Cell biological regulation of division fate in vertebrate neuroepithelial cells. Dev Dyn 240:1865–1879. doi: 10.1002/dvdy.22684

Zaghloul NA, Moody SA (2007) Alterations of rx1 and pax6 expression levels at neural plate stages differentially affect the production of retinal cell types and maintenance of retinal stem cell qualities. Dev Biol 306:222–240. doi: 10.1016/j.ydbio.2007.03.017

