## Supplemental Figures for "Loss of Nance-Horan Syndrome b (*nhsb*) prevents expansion growth of retinal progenitor cells by selective up-regulation of *Δ113p53*"

**Figure S1**

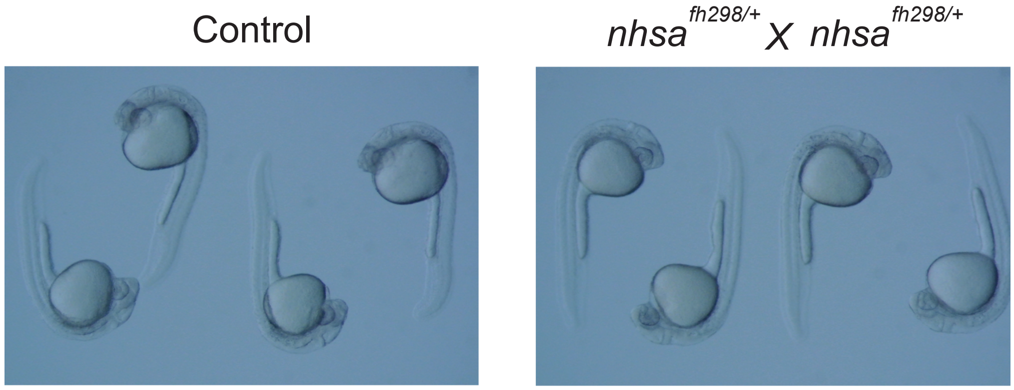


**Fig. S1**

Analysis of progeny generated from a nhsb$^{fh298/+}$ incross. No observable morphological defects were noticed in progeny generated from these crosses.

**Figure S2**

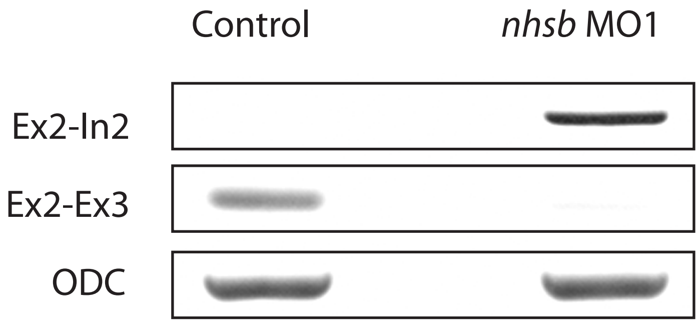


**Fig. S2**

RT-PCR validation of mis-splicing in *nhsb* MO1-injected embryos with retention of intron2 (In2) in the transcript using primers spanning exon2 (Ex2) to In2, and a failure to detect amplicons of transcript with proper splicing of Ex2 to Ex3. RT-PCR of ODC was used as a control.

### Figure S3

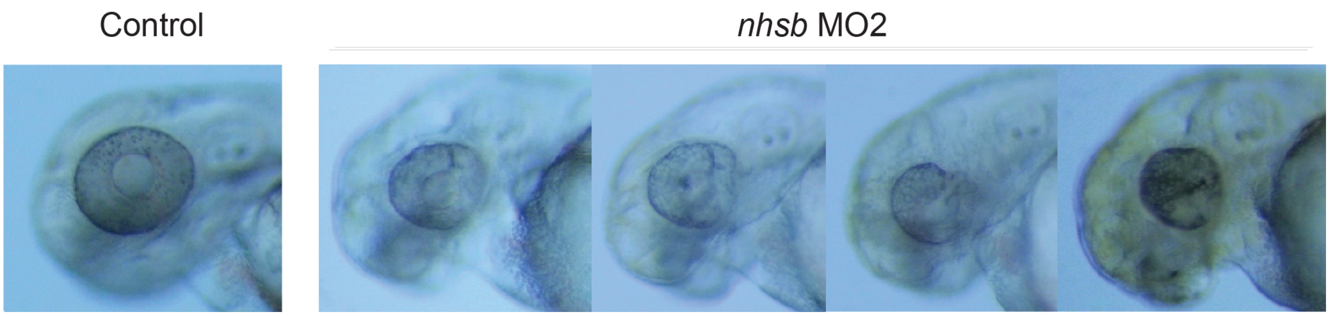


### Fig. S3

*nhsb* depletion leads to a small eye phenotype. conpered to control, embryos injected with nhsb MO2 ( I2E3) led to a small eye phenotype. Several examples of embryos injected with nhsb MO2 are shown. This was similar to that seen with nhsb MO1.

### Figure S4

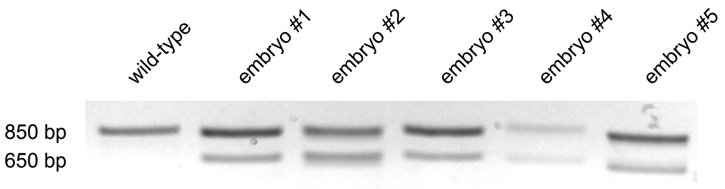


### Fig. S4

Validation of mutagenesis at the *nhsb* exon1 locus using T7 endonuclease I assay on embryos injected with Cas9 mRNA and *nhsb* ex1 gRNA.

**Figure S5**

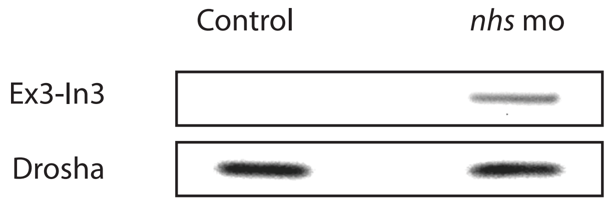


**Fig. S5**

RT-PCR validation of mis-splicing in *nhs* MO-injected *Xenopus tropicalis* embryos with retention of intron3 (In3) in the transcript using primers spanning exon3 (Ex2) to In3. RT-PCR of *drosha* was used as a control.

**Figure S6**

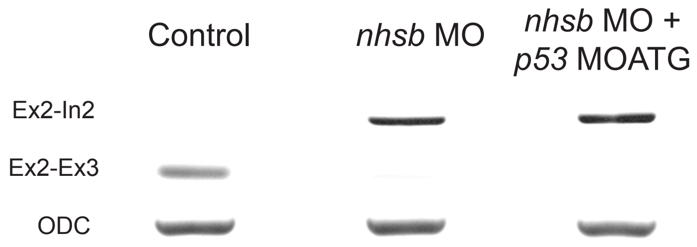


**Fig. S6**

RT-PCR validation of mis-splicing in zebrafish injected with *nhsb* MO combined with *p53* MOATG. Importantly, co-injection of *p53* MOATG had no effect on the splicing defects caused by *nhsb* MO, including retention of intron3 (In3) in the transcript and misplicing from Ex2 to Ex 3. RT-PCR of *odc* was used as a control.

### Figure S7

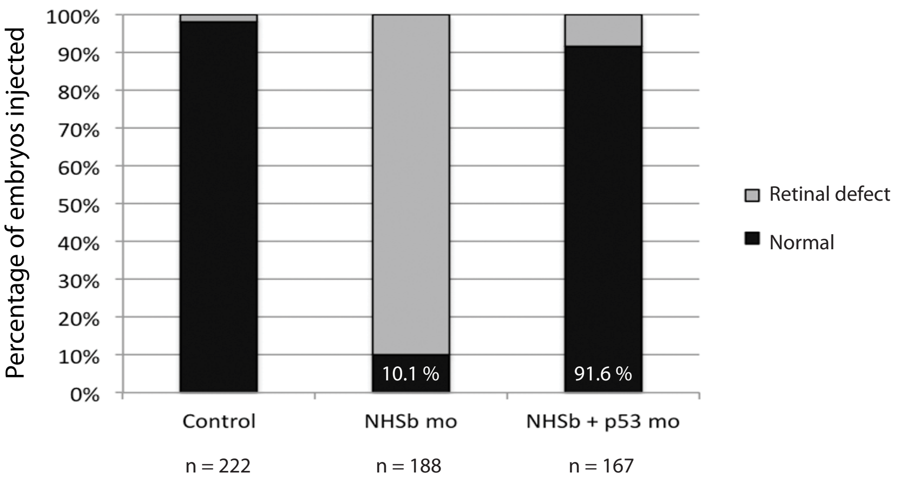


### Fig. S7

Knockdown of p53 rescues *nhsb* morphant phenotype. Embryos injected with nhsb MO alone or in combination with p53 MOATG were scored for retinal size. The majority of embryos coinjected with nhsb and p53 MOATG had normal eyes compared to embryos injected with nhsb MO alone.

Supplementary Table 1: Primers used for RT-PCR

| Primer | Sequence | Cycles |
|---|---|---|
| p53-full length - forward | TGGAGAGGAGGTCGGCAAAATCAA | 25 |
| p53- full length - reverse | GACTGCGGGAACCTGAGCCTAAAT | |
| p21-forward | GAAGCGCAAACAGACCAACAT | 24 |
| p21-reverse | GCAGCTCAATTACGATAAAGA | |
| mdm2-forward | CTCGCAGTGAGGGCAGTGAAG | 25 |
| mdm2-reverse | TCTAGGCACGTAGCGGGAAGG | |
| Bax-forward | GGAGGCGATACGGGCAGTG | 28 |
| Bax-reverse | TTGCGAATCACCAATGCTGTG | |
| cyclin-G1-forward | GCCCTTTACAGTCCAGCCCAAATC | 28 |
| cyclin-G1-reverse | CTGTGCCTCAAGCCTCTCGATGTA | |
| D113p53-forward | ATATCCTGGCGAACATTTGGAGGG | 22 |
| D113p53-reverse | CCTCCTGGTCTTGTAATGTCAC | |
| elfa-forward | CTTCTCAGGCTGACTGTGC | 20 |
| elfa-reverse | CCGCTAGCATTACCCTCC | |
| nhsb exon2 F1 | CACGGCCAGCGTGTCTGGAG | |
| nhsb exon3 R1 | CCTCTCTGCTCTCGCCGTCTGA | |
| nhsb exon2 F2 | GCTGCACCGACATGCCAAGC | |
| nhsb intron2 R2 | TCCACACACACACACGGTGCAA | |
| drosha F | TTACAGACCGCTGTTTGCTG | |
| drosha R | CAATTCGAGAGGGAGTTTCG | |
| ODC forward | ACACTATGACGGCTTGCACCG | |
| ODC reverse | CCCACTGACTGCACGATCTGG | |
| xt_nhs forward | ACCGCCATGCCAAGCAGAGT | |
| xt_nhs intron reverse1 | GGGTTTTCATGAGCTGTATGCCACA | |
| xt_nhs exon reverse2 | GGTGCTCTTTGACTTCTCGTCTCTTCA | |

### SAM positive genes

| Probe ID | Gene Tile | Gene Symbol |
|---|---|---|
| Dr.3758.1.A1_at | apoptosis antagonizing transcription factor | aatf |
| Dr.20276.1.A1_at | angiotensin I converting enzyme (peptidyl-dipeptidase A) 1 | ace |
| Dr.13171.1.A1_at | acidic repeat containing | acrc |
| Dr.26324.1.A1_at | acyl-CoA synthetase short-chain family member 1 | acss1 |
| Dr.21047.1.S1_at | activin receptor IIb | acvr2b |
| Dr.10904.2.S2_at | adducin 3 (gamma) a | add3a |
| Dr.9210.2.S1_a_at | acireductone dioxygenase 1 \| uncharacterized LOC554736 | adi1 \| LOC554736 |
| Dr.4408.1.S1_at | adenosine kinase a | adka |
| Dr.13359.1.S1_at | adenylosuccinate lyase | adsl |
| Dr.18323.1.S1_at | "adenylosuccinate synthase, like" | adssl |
| Dr.19099.1.A1_at | aspartylglucosaminidase | aga |
| Dr.6924.1.S1_at | S-adenosylhomocysteine hydrolase | ahcy |
| Dr.24918.1.A1_at | A kinase (PRKA) anchor protein (gravin) 12b | akap12b |
| Dr.26094.1.A1_at | A kinase (PRKA) anchor protein 17A | akap17a |
| Dr.4274.1.A1_at | "aldo-keto reductase family 7, member A3 (aflatoxin aldehyde reductase)" | akr7a3 |
| Dr.19785.1.A1_at | "alkB, alkylation repair homolog 6 (E. coli)" | alkbh6 |
| Dr.1368.2.A1_at | Aly/REF export factor | alyref |
| Dr.25421.1.S1_at | "anillin, actin binding protein" | anln |
| Dr.25562.1.A1_at | "acidic (leucine-rich) nuclear phosphoprotein 32 family, member B" | anp32b |
| Dr.5499.1.S1_at | "adaptor-related protein complex 1, sigma 2 subunit" | ap1s2 |
| Dr.4447.1.S1_at | anterior pharynx defective 1B | aph1b |
| Dr.5674.2.S1_at | apolipoprotein C-I like | apoc1l |
| Dr.4828.2.A1_at | ADP-ribosylation factor 1 | arf1 |
| Dr.6190.1.A1_at | ADP-ribosylation factor 6a \| ADP-ribosylation factor 6b | arf6a \| arf6b |
| Dr.9086.1.A1_at | rho/rac guanine nucleotide exchange factor (GEF) 18a | arhgef18a |
| Dr.22964.1.S1_at | rho/rac guanine nucleotide exchange factor (GEF) 18b | arhgef18b |
| Dr.24199.1.A1_s_at | cAMP-regulated phosphoprotein 19b | arpp19b |
| Dr.4206.1.A1_at | ASF1 anti-silencing function 1 homolog Bb | asf1bb |
| Dr.4095.1.A1_at | argininosuccinate synthetase 1 | ass1 |
| Dr.25732.1.A1_at | "ATPase family, AAA domain containing 3B" | atad3b |
| Dr.14282.1.S1_at | activating transcription factor 3 | atf3 |
| Dr.9110.1.A1_at | activating transcription factor 5b | atf5b |
| Dr.481.1.S1_at | 5-aminoimidazole-4-carboxamide ribonucleotide formyltransferase/IMP cyclohydrolase | atic |
| Dr.566.1.S1_at | atonal homolog 1a | atoh1a |
| Dr.888.1.S1_at | "ATPase, Na+/K+ transporting, alpha 1 polypeptide" | atp1a1 |
| Dr.11552.1.S1_at | "ATPase, Ca++ transporting, cardiac muscle, fast twitch 1 like" | atp2a1l |
| Dr.2043.1.A1_at | AU RNA binding protein/enoyl-Coenzyme A hydratase | auh |
| Dr.8332.1.A1_at | aurora kinase B | aurkb |
| Dr.23942.1.S1_at | "UDP-GlcNAc:betaGal beta-1,3-N-acetylglucosaminyltransferase 5a" | b3gnt5a |
| Dr.16002.2.S1_at | "bromodomain adjacent to zinc finger domain, 1A" | baz1a |
| Dr.19405.1.A1_at | B-cell receptor-associated protein 31 | bcap31 |
| Dr.5925.1.A1_at | brambleberry | bmb |
| Dr.25506.2.A1_at | "bone morphogenetic protein receptor, type 1aa" | bmpr1aa |
| Dr.8289.1.S1_at | "bone morphogenetic protein receptor, type 1ba" | bmpr1ba |
| Dr.5508.1.S1_at | "BMS1-like, ribosome assembly protein (yeast)" | bms1l |
| Dr.12295.2.A1_at | block of proliferation 1 | bop1 |
| Dr.15442.1.A1_at | BRCA1 interacting protein C-terminal helicase 1 | brip1 |
| Dr.6511.1.S1_at | B-cell translocation gene 2 | btg2 |
| Dr.17403.1.A1_s_at | BUB1 mitotic checkpoint serine/threonine kinase | bub1 |

| Probe ID | Gene Tile | Gene Symbol |
|---|---|---|
| Dr.1506.1.S1_at | NHP2 ribonucleoprotein homolog (yeast) | nhp2 |
| Dr.13635.1.S1_at | NHP2 non-histone chromosome protein 2-like 1b (S. cerevisiae) | nhp2l1b |
| Dr.17882.1.A1_at | nidogen 2a (osteonidogen) | nid2a |
| Dr.16108.1.S1_at | nuclear import 7 homolog (S. cerevisiae) | nip7 |
| Dr.17452.1.S1_at | nipsnap homolog 3 (C. elegans) | nipsnap3 |
| Dr.23851.1.A1_at | NK2 transcription factor related 2b | nkx2.2b |
| Dr.2745.1.A1_at | NMD3 homolog (S. cerevisiae) | nmd3 |
| Dr.7209.1.S1_at | "non-metastatic cells 6, protein expressed in (nucleoside-diphosphate kinase)" | nme6 |
| Dr.4323.1.A1_at | N-myristoyltransferase 1a | nmt1a |
| Dr.9728.1.A1_at | NIN1/RPN12 binding protein 1 homolog (S. cerevisiae) | nob1 |
| Dr.1412.2.A1_at | nucleolar protein 11 | nol11 |
| Dr.24822.1.S1_at | nucleolar protein 12 | nol12 |
| Dr.1726.1.A1_at | nucleolar protein family 6 (RNA-associated) | nol6 |
| Dr.12637.1.A1_at | nucleolar protein 9 | nol9 |
| Dr.5522.1.S1_at | nucleolar and coiled-body phosphoprotein 1 | nolc1 |
| Dr.18213.1.A1_at | nodal modulator | nomo |
| Dr.14960.1.S1_at | NOP2 nucleolar protein homolog (yeast) | nop2 |
| Dr.20916.1.A1_at | NOP58 ribonucleoprotein homolog (yeast) | nop58 |
| Dr.282.1.S1_at | NOP58 ribonucleoprotein homolog (yeast) | nop58 |
| Dr.25314.1.A1_at | notch homolog 3 | notch3 |
| Dr.22.1.S1_at | "nuclear receptor subfamily 2, group F, member 5" | nr2f5 |
| Dr.16181.1.A1_at | "nuclear receptor subfamily 2, group F, member 6b" | nr2f6b |
| Dr.4037.1.A1_at | N-terminal Xaa-Pro-Lys N-methyltransferase 1 | ntmt1 |
| Dr.15726.1.S1_at | "NUF2, NDC80 kinetochore complex component, homolog" | nuf2 |
| Dr.9746.9.A1_at | nucleoporin 107 | nup107 |
| Dr.5851.1.S1_at | nucleoporin 35 | nup35 |
| Dr.16553.1.A1_at | nucleoporin 37 | nup37 |
| Dr.16507.1.A1_at | nucleoporin 43 | nup43 |
| Dr.16507.2.S1_at | nucleoporin 43 | nup43 |
| Dr.20267.1.A1_at | nucleoporin 62 like | nup62l |
| Dr.4121.1.A1_at | nucleoporin 88 | nup88 |
| Dr.4435.1.A1_at | nucleoporin like 1 | nupl1 |
| Dr.14413.1.A1_at | nucleoporin like 2 | nupl2 |
| Dr.2359.1.S1_at | nuclear VCP-like | nvl |
| Dr.17474.1.S1_at | 2-oxoglutarate and iron-dependent oxygenase domain containing 1 | ogfod1 |
| Dr.4155.1.A1_at | oligodendrocyte transcription factor 3 | olig3 |
| Dr.17944.1.S1_at | "origin recognition complex, subunit 5" | orc5 |
| Dr.24945.1.S1_at | "origin recognition complex, subunit 6" | orc6 |
| Dr.7063.1.A1_at | O-sialoglycoprotein endopeptidase | osgep |
| Dr.26322.1.A1_at | "proliferation-associated 2G4, a" | pa2g4a |
| Dr.4846.1.S1_at | "proliferation-associated 2G4, a" | pa2g4a |
| Dr.5506.1.A1_at | "proliferation-associated 2G4, b" | pa2g4b |
| Dr.5701.1.S1_at | PAK1 interacting protein 1 | pak1ip1 |
| Dr.14405.1.S1_at | proliferation associated nuclear element | pane1 |
| Dr.12458.1.A1_at | "poly (ADP-ribose) polymerase family, member 2" | parp2 |
| Dr.17254.1.A1_at | PARP1 binding protein | parpbp |
| Dr.17401.1.A1_at | prolyl-tRNA synthetase (mitochondrial) | pars2 |
| Dr.522.1.A1_at | PAX interacting (with transcription-activation domain) protein 1 | paxip1 |
| Dr.18893.1.S1_at | "cleavage and polyadenylation factor subunit, homolog (S. cerevisiae)" | pcf11 |

| | | |
|---|---|---|
| Dr.6627.1.A1_at | budding uninhibited by benzimidazoles 1 homolog beta a | bub1ba |
| Dr.11708.1.S1_at | basic leucine zipper and W2 domains 2 | bzw2 |
| Dr.17482.1.S1_at | c1orf109 homolog (H. sapiens) | c1orf109 |
| Dr.25459.1.A1_at | carbonic anhydrase VIII | ca8 |
| Dr.21713.1.A1_at | --- \| --- | CABIN1 \| SI:CH211-53F16.1 |
| Dr.15338.1.A1_at | cache domain containing 1 | cachd1 |
| Dr.2668.1.S1_at | "carbamoyl-phosphate synthetase 2, aspartate transcarbamylase, and dihydroorotase" | cad |
| Dr.2668.2.A1_at | "carbamoyl-phosphate synthetase 2, aspartate transcarbamylase, and dihydroorotase" | cad |
| Dr.18018.1.A1_at | cancer susceptibility candidate 5 \| uncharacterized LOC100537099 | casc5 \| LOC100537099 |
| Dr.18018.2.S1_at | cancer susceptibility candidate 5 \| uncharacterized LOC100537099 | casc5 \| LOC100537099 |
| DrAffx.1.14.S1_at | "core-binding factor, beta subunit" | cbfb |
| Dr.20034.1.S1_at | "chromobox homolog 8a (Pc class homolog, Drosophila)" | cbx8a |
| Dr.8713.1.A1_at | coiled-coil domain containing 93 | ccdc93 |
| Dr.25190.1.S1_at | cyclin A2 | ccna2 |
| Dr.24753.1.S1_at | cyclin D1 | ccnd1 |
| Dr.24753.1.S2_at | cyclin D1 | ccnd1 |
| Dr.1212.1.S1_at | cyclin G1 | ccng1 |
| Dr.25673.1.S1_at | cell division cycle 25 homolog | cdc25 |
| Dr.25673.2.S1_at | cell division cycle 25 homolog | cdc25 |
| Dr.8228.1.S1_at | cell division cycle 25 homolog | cdc25 |
| Dr.3850.1.A1_at | cell division cycle 6 homolog (S. cerevisiae) | cdc6 |
| Dr.17883.1.S1_at | cyclin-dependent kinase 2 | cdk2 |
| Dr.8833.1.S1_at | "cysteine dioxygenase, type I" | cdo1 |
| Dr.14422.1.A1_at | cell adhesion molecule-related/down-regulated by oncogenes | cdon |
| Dr.1019.1.A1_at | chromatin licensing and DNA replication factor 1 | cdt1 |
| Dr.6575.1.S1_at | "CCAAT/enhancer binding protein (C/EBP), beta" | cebpb |
| Dr.18034.1.A1_at | "CCAAT/enhancer binding protein (C/EBP), zeta" | cebpz |
| Dr.361.1.S1_at | histone H3-like centromeric protein A \| selenoprotein H | cenpa \| seph |
| Dr.18241.1.A1_at | centromere protein J | cenpj |
| Dr.12523.1.A1_at | centrosomal protein 128 | cep128 |
| Dr.20507.1.A1_at | centrosomal protein 55 like | cep55l |
| Dr.16276.1.A1_at | cingulin-like 1 | cgnl1 |
| Dr.6898.1.A1_at | coiled-coil-helix-coiled-coil-helix domain containing 2 | chchd2 |
| Dr.7190.1.A1_at | coiled-coil-helix-coiled-coil-helix domain containing 3 | chchd3 |
| Dr.24974.1.S1_at | chromodomain helicase DNA binding protein 1-like | chd1l |
| Dr.15882.1.A1_at | CHK1 checkpoint homolog (S. pombe) | chek1 |
| Dr.26409.1.A1_at | "checkpoint with forkhead and ring finger domains, E3 ubiquitin protein ligase" | chfr |
| Dr.10196.1.S1_at | chromatin modifying protein 2Bb | chmp2bb |
| Dr.20903.1.S1_at | "CTF18, chromosome transmission fidelity factor 18 homolog (S. cerevisiae)" | chtf18 |
| Dr.24240.1.S1_at | "CTF8, chromosome transmission fidelity factor 8 homolog (S. cerevisiae)" | chtf8 |
| Dr.7926.1.A1_at | "cirrhosis, autosomal recessive 1A (cirhin)" | cirh1a |
| Dr.3606.1.A1_at | clustered mitochondria (cluA/CLU1) homolog | cluh |
| Dr.24219.1.S1_at | "CCHC-type zinc finger, nucleic acid binding protein a" | cnbpa |
| Dr.24219.1.S2_at | "CCHC-type zinc finger, nucleic acid binding protein a" | cnbpa |

| | | |
|---|---|---|
| Dr.15074.1.A1_at | polycomb group ring finger 6 | pcgf6 |
| Dr.372.1.S1_at | l-isoaspartyl protein carboxyl methyltransferase | pcmt |
| Dr.1348.1.S1_at | proliferating cell nuclear antigen | pcna |
| Dr.9847.2.A1_at | phosducin a | pdca |
| Dr.13356.1.S1_at | programmed cell death 11 | pdcd11 |
| Dr.23128.1.A1_at | PDZ and LIM domain 7 | pdlim7 |
| Dr.9343.1.S1_x_at | "PDS5, regulator of cohesion maintenance, homolog A (S. cerevisiae)" | pds5a |
| Dr.2260.1.S1_at | pelota homolog (Drosophila) | pelo |
| Dr.13701.1.A1_at | progressive external ophthalmoplegia 1 | peo1 |
| Dr.10257.1.A1_at | peptidase D | pepd |
| Dr.191.1.S1_at | pescadillo | pes |
| Dr.1058.1.S1_at | prohibitin | phb |
| Dr.5033.1.A1_at | prohibitin 2 | phb2 |
| Dr.7304.1.S1_at | PHD finger protein 6 | phf6 |
| Dr.11242.1.A1_at | "pleckstrin homology-like domain, family A, member 3" | phlda3 |
| Dr.14395.1.A1_at | PIH1 domain containing 1 | pih1d1 |
| Dr.58.1.A1_at | "phosphoinositide-3-kinase, regulatory subunit 3a (gamma)" | pik3r3a |
| Dr.13623.1.S1_at | PITH (C-terminal proteasome-interacting domain of thioredoxin-like) domain containing 1 | pithd1 |
| Dr.18144.1.A1_a_at | pleiomorphic adenoma gene X | plagx |
| Dr.2929.1.A1_at | pleiomorphic adenoma gene X | plagx |
| Dr.6805.1.A1_at | "phospholipase C, gamma 1" | plcg1 |
| Dr.21825.1.A1_at | "pleckstrin homology domain containing, family A (phosphoinositide binding specific) member 8" | plekha8 |
| Dr.24256.1.S1_at | "pleckstrin homology domain containing, family F (with FYVE domain) member 2" | plekhf2 |
| Dr.20131.4.S1_at | polo-like kinase 1 (Drosophila) | plk1 |
| Dr.2155.1.S1_at | polo-like kinase 3 (Drosophila) | plk3 |
| Dr.5434.1.S1_at | proteolipid protein 1a | plp1a |
| Dr.5434.1.S2_at | proteolipid protein 1a | plp1a |
| DrAffx.2.16.S1_at | phorbol-12-myristate-13-acetate-induced protein 1 | pmaip1 |
| Dr.21058.1.S1_at | POC1 centriolar protein homolog A (Chlamydomonas) | poc1a |
| Dr.15162.2.S1_at | protein O-fucosyltransferase 1 | pofut1 |
| Dr.519.1.A1_at | "polymerase (DNA directed), alpha 1" | pola1 |
| Dr.4879.1.S1_at | "polymerase (DNA directed), alpha 2" | pola2 |
| Dr.9175.1.A1_at | "polymerase (DNA directed), delta 1, catalytic subunit" | pold1 |
| Dr.25641.1.A1_at | "polymerase (DNA-directed), delta 3, accessory subunit" | pold3 |
| Dr.2435.1.S1_at | "polymerase (DNA directed), epsilon 2" | pole2 |
| Dr.6031.1.A1_at | "polymerase (DNA directed), lambda" | poll |
| Dr.770.1.A1_at | polymerase (RNA) I polypeptide A | polr1a |
| Dr.3838.1.A1_at | polymerase (RNA) I polypeptide C | polr1c |
| Dr.10110.1.S1_a_at | "polymerase (RNA) II (DNA directed) polypeptide E, b" | polr2eb |
| Dr.5495.1.S1_at | polymerase (RNA) III (DNA directed) polypeptide A | polr3a |
| Dr.4746.1.S1_at | polymerase (RNA) III (DNA directed) polypeptide E | polr3e |
| Dr.6372.1.A1_at | POM121 membrane glycoprotein (rat) | pom121 |
| Dr.26358.1.A1_at | phosphoribosyl pyrophosphate amidotransferase | ppat |
| Dr.6397.1.S1_at | peptidylprolyl isomerase (cyclophilin)-like 4 | ppil4 |

| | | |
|---|---|---|
| Dr.116.1.S1_at | cyclin Pas1/PHO80 domain containing 1 | cnppd1 |
| Dr.4832.1.A1_at | "collagen, type XV, alpha 1b" | col15a1b |
| Dr.17635.1.A1_at | "collagen, type XXVIII, alpha 1" | col28a1 |
| Dr.11526.1.S1_at | COP9 constitutive photomorphogenic homolog subunit 3 | cops3 |
| Dr.21.1.A1_at | "COX10 homolog, cytochrome c oxidase assembly protein, heme A: farnesyltransferase (yeast)" | cox10 |
| Dr.3731.1.A1_at | cleavage and polyadenylation specific factor 3 | cpsf3 |
| Dr.21961.1.A1_at | "cellular retinoic acid binding protein 2, b" | crabp2b |
| Dr.25494.1.A1_at | v-crk sarcoma virus CT10 oncogene homolog (avian) | crk |
| Dr.18310.1.S1_at | cryptochrome DASH | cry-dash |
| Dr.10332.1.S1_at | cryptochrome 5 | cry5 |
| Dr.13038.1.S1_at | "crystallin, lambda 1" | cryl1 |
| Dr.16251.1.A1_at | cysteine-serine-rich nuclear protein 1a | csrnp1a |
| Dr.1279.1.S1_at | "cleavage stimulation factor, 3' pre-RNA, subunit 3" | cstf3 |
| Dr.3749.1.A1_at | "CTD (carboxy-terminal domain, RNA polymerase II, polypeptide A) small phosphatase like 2b" | ctdspl2b |
| Dr.8148.1.S1_at | "catenin (cadherin-associated protein), alpha 1" | ctnna1 |
| Dr.3048.1.A1_at | "catenin, beta like 1" | ctnnbl1 |
| Dr.25770.1.S1_at | CTP synthase 1a | ctps1a |
| Dr.3485.1.S1_at | cathepsin A | ctsa |
| Dr.3374.1.S1_at | "cathepsin B, a" | ctsba |
| Dr.4782.1.S1_at | cathepsin C | ctsc |
| Dr.19238.1.S1_at | cathepsin D | ctsd |
| Dr.15507.2.A1_at | cathepsin L.1 | ctsl.1 |
| Dr.21960.1.A1_at | "cathepsin L, 1 b" | ctsl1b |
| Dr.25683.1.S1_at | "cathepsin L, 1 b \| cathepsin L1-like \| cathepsin L1-like \| Cathepsin L1-like \| Cathepsin L1-like \| cathepsin L1-like \| si:dkey-269i1.3 \| si:dkey-26g8.4 \| wu:fa26c03 \| wu:fb37b09 \| zgc:174153 \| zgc:174855" | ctsl1b \| LOC100001785 \| LOC100536033 \| MGC174152 \| MGC174155 \| MGC174857 \| si:dkey-269i1.3 \| si:dkey-26g8.4 \| wu:fa26c03 \| wu:fb37b09 \| zgc:174153 \| zgc:174855 |
| Dr.24219.5.S1_at | "cathepsin S, b.1" | ctssb.1 |
| Dr.11380.1.A1_at | "cytochrome P450, family 27, subfamily C, polypeptide 1 \| excision repair cross-complementing rodent repair deficiency, complementation group 3" | cyp27c1 \| ercc3 |
| Dr.11729.2.A1_at | "cytochrome P450, family 2, subfamily AD, polypeptide 2" | cyp2ad2 |
| Dr.3552.1.A1_at | "cytochrome P450, family 2, subfamily X, polypeptide 10.2 \| cytochrome P450 2F2-like" | cyp2x10.2 \| LOC100148115 |
| Dr.17457.1.A1_at | "cytochrome P450, family 46, subfamily A, polypeptide 2" | cyp46a2 |
| Dr.24486.1.A1_at | cysteine and tyrosine-rich protein 1 | cyyr1 |
| Dr.15604.1.A1_at | dynein assembly factor with WDR repeat domains 1 | daw1 |
| Dr.8661.1.A1_at | death-associated protein 6 | daxx |
| Dr.8070.1.S1_at | developing brain homeobox 1a | dbx1a |
| Dr.245.1.A1_at | DEAD/H (Asp-Glu-Ala-Asp/His) box helicase 11 | ddx11 |
| Dr.24235.1.A1_at | DEAD (Asp-Glu-Ala-Asp) box polypeptide 18 | ddx18 |
| Dr.3281.1.A1_at | DEAD (Asp-Glu-Ala-Asp) box polypeptide 31 | ddx31 |
| Dr.13799.1.S1_x_at | DEAD (Asp-Glu-Ala-Asp) box polypeptide 52 | ddx52 |
| Dr.16883.1.S1_at | DEAD (Asp-Glu-Ala-Asp) box polypeptide 55 | ddx55 |

| | | |
|---|---|---|
| Dr.17119.1.A1_at | "protein phosphatase 2, regulatory subunit B'', beta" | ppp2r3b |
| Dr.25412.1.A1_a_at | "protein phosphatase 4, regulatory subunit 1" | PPP4R1 (2 of 2) |
| Dr.2790.1.A1_at | "protein phosphatase 5, catalytic subunit" | ppp5c |
| Dr.165.1.S1_at | "protein phosphatase 6, regulatory subunit 3" | ppp6r3 |
| Dr.2988.1.A1_at | "peroxisome proliferator-activated receptor gamma, coactivator-related 1" | pprc1 |
| Dr.14630.1.A1_at | peptidylprolyl isomerase domain and WD repeat containing 1 | ppwd1 |
| Dr.377.1.S1_at | primase polypeptide 1 | prim1 |
| Dr.16922.1.A1_at | "protein kinase, cAMP-dependent, regulatory, type I, alpha (tissue specific extinguisher 1) b" | prkar1ab |
| Dr.5651.1.S1_at | protein arginine methyltransferase 5 | prmt5 |
| Dr.3855.1.S1_at | protein arginine N-methyltransferase 7 | prmt7 |
| Dr.3564.1.S1_at | PRP19/PSO4 homolog (S. cerevisiae) | prp19 |
| Dr.2143.1.A1_at | PRP38 pre-mRNA processing factor 38 (yeast) domain containing B | prpf38b |
| Dr.23930.1.S1_at | PRP40 pre-mRNA processing factor 40 homolog A (yeast) | prpf40a |
| Dr.2896.1.S1_at | phosphoribosyl pyrophosphate synthetase 1A | prps1a |
| Dr.13586.1.A1_at | proline rich 11 | prr11 |
| Dr.20872.1.S1_at | "proteasome (prosome, macropain) subunit, alpha type, 6a" | psma6a |
| Dr.15385.1.S1_at | "proteasome (prosome, macropain) subunit, beta type, 4" | psmb4 |
| Dr.25360.1.A1_s_at | "proteasome (prosome, macropain) 26S subunit, ATPase, 1b" | psmc1b |
| Dr.8135.1.S1_at | proteasome activator subunit 1 | psme1 |
| Dr.20803.1.S1_at | polypyrimidine tract binding protein 1a | ptbp1a |
| Dr.5460.1.A1_at | polypyrimidine tract binding protein 1a | ptbp1a |
| Dr.20073.1.A1_at | polypyrimidine tract binding protein 1b | ptbp1b |
| Dr.16120.1.S1_at | prostaglandin E synthase 2-like | ptgesl |
| Dr.10653.1.A1_s_at | "protein tyrosine phosphatase, non-receptor type 12" | ptpn12 |
| Dr.14692.1.S1_at | "protein tyrosine phosphatase, non-receptor type 12" | ptpn12 |
| Dr.19015.1.A1_at | "protein tyrosine phosphatase, non-receptor type 13" | ptpn13 |
| Dr.12168.1.A1_at | peptidyl-tRNA hydrolase 2 | ptrh2 |
| Dr.14734.1.A1_a_at | pseudouridylate synthase 7 homolog (S. cerevisiae) | pus7 |
| Dr.20928.1.S1_at | parvalbumin 1 | pvalb1 |
| Dr.18470.1.A1_at | poliovirus receptor-related 3 like | pvrl3l |
| Dr.16123.1.S1_at | PWP1 homolog (S. cerevisiae) | pwp1 |
| Dr.7795.2.A1_at | quinoid dihydropteridine reductase b1 | qdprb1 |
| Dr.6000.1.A1_at | glutamine-rich 1 | qrich1 |
| Dr.9747.1.A1_at | queuine tRNA-ribosyltransferase 1 | qtrt1 |
| Dr.15011.1.A1_at | "RAB12, member RAS oncogene family" | rab12 |
| Dr.2730.1.S1_at | RAB interacting factor | rabif |
| Dr.15214.1.S1_at | RAD18 homolog (S. cerevisiae) | rad18 |
| Dr.1514.1.A1_at | "RAD51 homolog (RecA homolog, E. coli) (S. cerevisiae)" | rad51 |

| | | |
|---|---|---|
| Dr.2066.1.A1_at | DEK oncogene | dek |
| Dr.4024.1.S1_at | DEAH (Asp-Glu-Ala-His) box polypeptide 15 | dhx15 |
| Dr.5257.1.S1_at | digestive organ expansion factor homolog | diexf |
| Dr.10492.1.S1_at | deltaA | dla |
| Dr.574.1.S1_at | deltaB | dlb |
| Dr.574.2.S1_at | deltaB | dlb |
| Dr.16183.1.S1_at | deltaC | dlc |
| Dr.8086.1.S1_s_at | deltaC | dlc |
| Dr.20958.1.S1_at | deltaD | dld |
| Dr.20429.2.S1_a_at | "discs, large (Drosophila) homolog-associated protein 5" | dlgap5 |
| Dr.7204.2.A1_a_at | dynamin 2b | dnm2b |
| Dr.5403.1.A1_at | "deoxynucleotidyltransferase, terminal, interacting protein 2" | dnttip2 |
| Dr.3526.1.S1_at | denticleless homolog (Drosophila) | dtl |
| Dr.5295.1.A1_at | "dystrobrevin, beta a \| dystrobrevin, gamma-like" | dtnba \| dtng |
| Dr.19959.1.S1_at | "dihydrouridine synthase 2-like, SMM1 homolog (S. cerevisiae)" | dus2l |
| Dr.26415.1.A1_x_at | "dihydrouridine synthase 2-like, SMM1 homolog (S. cerevisiae)" | dus2l |
| Dr.17122.1.A1_at | dihydrouridine synthase 3-like (S. cerevisiae) | dus3l |
| Dr.10567.1.A1_at | dual specificity phosphatase 16 | dusp16 |
| Dr.24920.1.S1_at | "dishevelled, dsh homolog 2 (Drosophila)" | dvl2 |
| Dr.4922.1.A1_at | E2F transcription factor 4 | e2f4 |
| Dr.21523.1.A1_at | E2F transcription factor 7 | e2f7 |
| Dr.21415.1.A1_at | E2F transcription factor 8 | e2f8 |
| Dr.14998.2.S1_at | "enoyl CoA hydratase 1, peroxisomal" | ech1 |
| Dr.12679.1.S1_at | epithelial cell transforming sequence 2 oncogene | ect2 |
| Dr.14507.1.A1_at | embryonic ectoderm development | eed |
| Dr.10127.1.A1_at | ephrin A5a | efna5a |
| Dr.12617.1.A1_at | ephrin B3b | efnb3b |
| Dr.17549.1.A1_at | early growth response 2a | egr2a |
| Dr.22232.1.A1_at | "enoyl-Coenzyme A, hydratase/3-hydroxyacyl Coenzyme A dehydrogenase" | ehhadh |
| Dr.15002.1.S1_a_at | "eukaryotic translation initiation factor 1A, X-linked, a" | eif1axa |
| Dr.15002.1.S1_at | "eukaryotic translation initiation factor 1A, X-linked, a" | eif1axa |
| Dr.17703.1.A1_at | eukaryotic translation initiation factor 2D | eif2d |
| Dr.17196.1.S1_at | "eukaryotic translation initiation factor 3, subunit Ja" | eif3ja |
| Dr.1756.1.S1_at | "eukaryotic translation initiation factor 4A, isoform 3" | eif4a3 |
| Dr.8.1.S1_at | eukaryotic translation initiation factor 5A2 | eif5a2 |
| Dr.8.1.S2_at | eukaryotic translation initiation factor 5A2 | eif5a2 |
| Dr.5598.1.S1_at | eukaryotic translation initiation factor 6 | eif6 |
| Dr.24196.1.S1_at | "ELAV (embryonic lethal, abnormal vision, Drosophila)-like 1 (Hu antigen R)" | elavl1 |
| Dr.4460.1.A1_at | essential meiotic endonuclease 1 homolog 1 (S. pombe) | eme1 |
| Dr.4460.2.S1_at | essential meiotic endonuclease 1 homolog 1 (S. pombe) | eme1 |
| Dr.4460.3.S1_at | essential meiotic endonuclease 1 homolog 1 (S. pombe) | eme1 |
| Dr.14980.1.S1_at | "enkurin, TRPC channel interacting protein" | enkur |
| Dr.3945.1.A1_at | erythrocyte membrane protein band 4.1 like 5 | epb41l5 |
| Dr.5765.1.S1_at | eph receptor A2 | epha2 |
| Dr.25694.1.S1_at | eph receptor A4a | epha4a |
| Dr.4703.1.S2_at | eph receptor B2b \| targeting protein for Xklp2-B-like | ephb2b \| LOC100536090 |
| Dr.9388.1.A1_at | "excision repair cross-complementing rodent repair deficiency, complementation group 6-like" | ercc6l |
| Dr.13342.1.A1_at | Ets2 repressor factor | erf |
| Dr.3337.1.S1_at | establishment of cohesion 1 homolog 2 | esco2 |
| Dr.4113.1.S1_at | eukaryotic translation termination factor 1 | etf1 |

| | | |
|---|---|---|
| Dr.1514.2.S1_at | "RAD51 homolog (RecA homolog, E. coli) (S. cerevisiae)" | rad51 |
| Dr.12483.1.A1_at | RAD51 associated protein 1 | rad51ap1 |
| Dr.321.1.S1_at | ras-related nuclear protein | ran |
| Dr.12411.1.S1_at | RAN binding protein 1 | ranbp1 |
| Dr.25678.2.A1_at | Ran GTPase activating protein 1b | rangap1b |
| Dr.13285.1.A1_at | "RasGEF domain family, member 1Bb" | rasgef1bb |
| Dr.11735.1.A1_at | "RAS-like, family 11, member A" | rasl11a |
| Dr.21306.1.A1_at | Ras association (RalGDS/AF-6) domain family 8b | rassf8b |
| Dr.1119.1.S1_at | "retinoblastoma binding protein 4, like" | rbb4l |
| Dr.3888.1.A1_at | RNA binding motif protein 24a | rbm24a |
| Dr.26430.1.A1_at | regulator of chromosome condensation 1 | rcc1 |
| Dr.12392.1.S1_at | ring finger and CHY zinc finger domain containing 1 | rchy1 |
| Dr.6949.1.A1_at | RNA terminal phosphate cyclase-like 1 | rcl1 |
| Dr.15761.1.A1_at | RecQ protein-like (DNA helicase Q1-like) | recql |
| Dr.25449.1.A1_at | RecQ protein-like 5 | recql5 |
| Dr.9525.1.A1_at | RALBP1 associated Eps domain containing 1 | reps1 |
| Dr.17100.1.A1_at | replication factor C (activator 1) 1 | rfc1 |
| Dr.4387.3.S1_at | replication factor C (activator 1) 5 | rfc5 |
| Dr.23832.1.S1_at | "regulatory factor X, 1b (influences HLA class II expression)" | rfx1b |
| Dr.20543.1.S1_at | "regulatory factor X, 4" | rfx4 |
| Dr.17903.1.A1_at | "rhomboid, veinlet-like 3 (Drosophila)" | rhbdl3 |
| Dr.25770.2.A1_at | "rhomboid, veinlet-like 3 (Drosophila)" | rhbdl3 |
| Dr.1709.1.S1_at | "ras homolog gene family, member Gc" | rhogc |
| Dr.2150.1.A1_at | RAP1 interacting factor homolog (yeast) | rif1 |
| Dr.2150.1.A1_x_at | RAP1 interacting factor homolog (yeast) | rif1 |
| Dr.14704.1.S1_at | "ribonuclease H2, subunit A" | rnaseh2a |
| Dr.4468.1.S1_at | "ribonuclease H2, subunit B" | rnaseh2b |
| Dr.1706.1.S1_at | "rho-associated, coiled-coil containing protein kinase 2a" | rock2a |
| Dr.7625.1.S1_at | replication protein A1 | rpa1 |
| Dr.4556.1.S1_at | replication protein A2 | rpa2 |
| Dr.16936.1.S1_at | RNA polymerase II associated protein 3 | rpap3 |
| Dr.17277.1.S1_at | Ras-related GTP binding Ca | rragca |
| Dr.732.1.S1_at | ribosomal RNA processing 15 homolog (S. cerevisiae) | rrp15 |
| Dr.24135.1.A1_at | ribosomal RNA processing 7 homolog A (S. cerevisiae) | rrp7a |
| Dr.7888.1.S1_at | RRS1 ribosome biogenesis regulator homolog (S. cerevisiae) | rrs1 |
| Dr.11481.1.A1_at | R-spondin homolog (Xenopus laevis) | rspo1 |
| Dr.18135.1.A1_at | rhotekin 2 | rtkn2 |
| Dr.103.1.S1_at | sal-like 4 (Drosophila) | sall4 |
| Dr.17653.2.S1_a_at | "spermidine/spermine N1-acetyltransferase 1a, duplicate 2" | sat1a.2 |
| Dr.1204.1.S1_at | SDA1 domain containing 1 | sdad1 |
| Dr.8010.1.S1_at | succinate dehydrogenase complex assembly factor 2 | sdhaf2 |
| Dr.20191.1.S1_at | SEH1-like (S. cerevisiae) | seh1l |
| Dr.18416.1.A1_at | selenium binding protein 1 | selenbp1 |
| Dr.1330.1.S1_at | "selenoprotein P, plasma, 1a" | sepp1a |
| Dr.16590.1.S1_at | Sep (O-phosphoserine) tRNA:Sec (selenocysteine) tRNA synthase | sepsecs |
| Dr.20973.1.S1_at | SET domain containing 6 | setd6 |
| Dr.24150.1.A1_at | serine hydroxymethyltransferase 2 (mitochondrial) | shmt2 |
| Dr.1632.1.A1_at | si:ch211-156j22.4 | si:ch211-156j22.4 |
| Dr.15996.1.S1_at | si:ch211-217k17.7 | si:ch211-217k17.7 |
| Dr.4479.1.A1_at | si:ch211-244o22.2 | si:ch211-244o22.2 |

| | | |
|---|---|---|
| Dr.7714.1.S1_at | ets variant 5b | etv5b |
| Dr.14893.1.A1_at | "nuclease EXOG, mitochondrial-like" | exog |
| Dr.9715.1.S1_a_at | exosome component 4 | exosc4 |
| Dr.17708.1.S1_at | exosome component 5 | exosc5 |
| Dr.3788.1.A1_at | enhancer of zeste homolog 2 (Drosophila) | ezh2 |
| Dr.17772.1.A1_at | coagulation factor II (thrombin) receptor-like 1.2 | f2rl1.2 |
| Dr.3676.1.A1_at | "family with sequence similarity 210, member A" | fam210a |
| Dr.11988.1.A1_at | "family with sequence similarity 53, member B" | fam53b |
| Dr.12682.1.A1_at | "Fanconi anemia, complementation group G" | fancg |
| Dr.12462.1.A1_at | FAST kinase domains 2 | fastkd2 |
| Dr.21842.1.S1_at | FAT tumor suppressor homolog 1 | fat1 |
| Dr.5345.1.S1_at | F-box protein 5 | fbxo5 |
| Dr.18882.1.A1_at | ferredoxin 1-like | fdx1l |
| Dr.1294.1.S1_at | flap structure-specific endonuclease 1 | fen1 |
| Dr.409.1.S1_at | fibroblast growth factor receptor 4 | fgfr4 |
| Dr.16271.1.A1_at | focadhesin | focad |
| Dr.12986.1.A1_a_at | v-fos FBJ murine osteosarcoma viral oncogene homolog | fos |
| Dr.12986.1.A1_at | v-fos FBJ murine osteosarcoma viral oncogene homolog | fos |
| Dr.17623.1.S1_at | forkhead box M1 | foxm1 |
| Dr.13699.1.S1_at | forkhead box N4 | foxn4 |
| Dr.18562.1.A1_a_at | folylpolyglutamate synthase | fpgs |
| Dr.18562.1.A1_at | folylpolyglutamate synthase | fpgs |
| Dr.18562.2.A1_x_at | folylpolyglutamate synthase | fpgs |
| Dr.16630.1.S1_at | "GA repeat binding protein, beta 2a" | gabpb2a |
| Dr.23587.1.A1_at | "growth arrest and DNA-damage-inducible, alpha, a" | gadd45aa |
| Dr.23694.1.A1_at | galactokinase 2 | galk2 |
| Dr.14883.1.S1_at | GAR1 ribonucleoprotein homolog (yeast) | gar1 |
| Dr.15740.1.A1_at | GC-rich sequence DNA-binding factor 2 | gcfc2 |
| Dr.1491.1.S1_at | "glycine cleavage system protein H (aminomethyl carrier), a" | gcsha |
| Dr.17767.1.S1_at | gem (nuclear organelle) associated protein 5 | gemin5 |
| Dr.17767.2.A1_at | gem (nuclear organelle) associated protein 5 | gemin5 |
| Dr.4685.1.S1_at | gem (nuclear organelle) associated protein 8 | gemin8 |
| Dr.15045.1.S1_at | GINS complex subunit 2 | gins2 |
| Dr.13962.1.S1_at | GINS complex subunit 3 | gins3 |
| Dr.8091.1.S1_at | GLI-Kruppel family member GLI2a | gli2a |
| Dr.2113.2.S1_a_at | GM2 ganglioside activator | gm2a |
| Dr.18605.1.A1_at | "glia maturation factor, gamma" | gmfg |
| Dr.14739.1.S1_at | "geminin, DNA replication inhibitor" | gmnn |
| Dr.8251.1.A1_at | guanine monphosphate synthetase | gmps |
| Dr.16124.1.S1_at | guanine nucleotide binding protein-like 2 (nucleolar) \| nucleolar GTP-binding protein 2-like | gnl2 \| LOC100536859 |
| Dr.7311.1.S1_at | guanine nucleotide binding protein-like 3 (nucleolar) | gnl3 |
| Dr.3178.1.A1_at | guanine nucleotide binding protein-like 3 (nucleolar)-like | gnl3l |
| Dr.10191.1.S1_at | golgi associated PDZ and coiled-coil motif containing | gopc |
| Dr.2793.1.A1_at | glypican 4 | gpc4 |
| Dr.17912.1.S1_at | glycoprotein M6Bb | gpm6bb |
| Dr.22965.1.S1_at | G-rich RNA sequence binding factor 1 | grsf1 |

| | | |
|---|---|---|
| Dr.133.1.A1_at | si:ch211-255a21.1 | si:ch211-255a21.1 |
| Dr.12375.1.S1_at | si:ch211-285f17.1 | si:ch211-285f17.1 |
| Dr.16494.1.A1_at | si:dkey-11e23.4 | si:dkey-11e23.4 |
| Dr.22416.1.A1_at | si:dkey-120m5.1 | si:dkey-120m5.1 |
| Dr.6105.1.A1_at | si:dkey-147f3.4 | si:dkey-147f3.4 |
| Dr.685.1.S1_at | si:dkey-170l10.1 | si:dkey-170l10.1 |
| Dr.22184.1.A1_at | si:dkey-269d20.3 | si:dkey-269d20.3 |
| Dr.4307.1.A1_s_at | --- | SI:DKEY-269O24.6 |
| Dr.12485.1.A1_at | si:dkey-27p23.3 | si:dkey-27p23.3 |
| Dr.16696.1.S1_at | si:dkey-286j15.3 | si:dkey-286j15.3 |
| Dr.15473.1.A1_at | si:dkey-97o5.1 | si:dkey-97o5.1 |
| Dr.3249.1.A1_at | si:dkeyp-113d7.1 | si:dkeyp-113d7.1 |
| Dr.12668.1.A1_at | si:dkeyp-26a9.2 | si:dkeyp-26a9.2 |
| Dr.6786.1.A1_at | si:dkeyp-35b8.5 | si:dkeyp-35b8.5 |
| Dr.12509.1.S1_at | si:dkeyp-84a8.8 | si:dkeyp-84a8.8 |
| Dr.18733.1.S1_at | si:dkeyp-89c11.2 | si:dkeyp-89c11.2 |
| Dr.25375.1.A1_at | spindle and kinetochore associated complex subunit 3 | ska3 |
| Dr.25571.1.A1_at | S-phase kinase-associated protein 2 (p45) | skp2 |
| Dr.24852.1.S1_at | stem-loop binding protein | slbp |
| Dr.4660.1.S1_at | "solute carrier family 16 (monocarboxylic acid transporters), member 8" | slc16a8 |
| Dr.24146.1.S1_at | "solute carrier family 16 (monocarboxylic acid transporters), member 9b" | slc16a9b |
| Dr.25679.1.S1_at | "solute carrier family 1 (glial high affinity glutamate transporter), member 3a" | slc1a3a |
| Dr.5307.1.S1_at | "solute carrier family 20, member 1b" | slc20a1b |
| Dr.676.1.S1_at | "solute carrier family 25 (mitochondrial carrier; adenine nucleotide translocator), member 4" | slc25a4 |
| Dr.3740.1.A1_at | "solute carrier family 7 (cationic amino acid transporter, y+ system), member 3a" | slc7a3a |
| Dr.10559.1.A1_at | "solute carrier organic anion transporter family, member 2B1" | slco2b1 |
| Dr.20402.1.A1_at | structural maintenance of chromosomes 2 | smc2 |
| Dr.18590.1.A1_at | structural maintenance of chromosomes flexible hinge domain containing 1 | smchd1 |
| Dr.24441.1.S1_at | "SMEK homolog 2, suppressor of mek1 (Dictyostelium)" | smek2 |
| Dr.13752.1.S1_at | small fragment nuclease | smfn |
| Dr.24766.1.S1_at | smoothened homolog (Drosophila) | smo |
| Dr.4416.2.A1_at | smoothened homolog (Drosophila) | smo |
| Dr.14033.1.A1_at | SPARC related modular calcium binding 1 | smoc1 |
| Dr.3569.1.S1_at | sphingomyelin phosphodiesterase 4 | smpd4 |
| Dr.12494.1.A1_at | SET and MYND domain containing 4 | smyd4 |
| Dr.7936.1.A1_at | small nuclear ribonucleoprotein polypeptides B and B1 | snrpb |
| Dr.3953.1.S1_at | "small nuclear ribonucleoprotein D3 polypeptide, like" | snrpd3l |
| Dr.5299.1.S1_at | sorting nexin 12 | snx12 |
| Dr.25917.1.S1_at | sorting nexin 7 | snx7 |
| Dr.6431.1.S1_at | suppressor of cytokine signaling 3a | socs3a |
| Dr.6314.1.S1_at | "superoxide dismutase 2, mitochondrial" | sod2 |
| Dr.5112.1.S3_at | SRY-box containing gene 11b | sox11b |
| Dr.20910.1.S1_at | SRY-box containing gene 19a | sox19a |
| Dr.8215.1.A1_at | SRY-box containing gene 21a | sox21a |
| Dr.10460.1.S1_at | SRY-box containing gene 9a | sox9a |
| Dr.11850.1.S2_at | SRY-box containing gene 9b | sox9b |

| | | |
|---|---|---|
| Dr.5624.1.S1_at | gsk3b interacting protein | gskip |
| Dr.1388.1.S1_at | "G1 to S phase transition 1, like" | gspt1l |
| Dr.1754.1.S1_at | glutathione S-transferase M | gstm |
| Dr.10042.1.A1_at | histone acetyltransferase 1 | hat1 |
| Dr.15779.1.S1_at | "HAUS augmin-like complex, subunit 4" | haus4 |
| Dr.15779.2.A1_at | "HAUS augmin-like complex, subunit 4" | haus4 |
| Dr.834.1.S1_at | "HAUS augmin-like complex, subunit 6" | haus6 |
| DrAffx.2.19.S1_at | hemoglobin beta embryonic-3 | hbbe3 |
| Dr.1758.1.S1_at | histone deacetylase 10 | hdac10 |
| Dr.5019.1.A1_at | HD domain containing 2 | hddc2 |
| Dr.15003.1.A1_at | haloacid dehalogenase-like hydrolase domain containing 2 | hdhd2 |
| Dr.1691.9.S1_at | hatching enzyme 1b | he1b |
| Dr.4330.1.A1_at | HEAT repeat containing 3 | heatr3 |
| Dr.1899.1.S1_at | hairy and enhancer of split-related 15.1 \| hairy and enhancer of split-related 15.2 \| transcription factor HES-5-like | her15.1 \| her15.2 \| LOC100534909 |
| Dr.1899.3.A1_at | hairy and enhancer of split-related 15.1 \| hairy and enhancer of split-related 15.2 \| transcription factor HES-5-like | her15.1 \| her15.2 \| LOC100534909 |
| Dr.1460.1.S1_at | hairy-related 2 | her2 |
| Dr.19467.1.A1_at | hairy and enhancer of split 6 (Drosophila) | hes6 |
| Dr.3681.1.S1_at | histone H1 like | histh1l |
| Dr.15391.1.S1_at | heterogeneous nuclear ribonucleoprotein D | hnrnpd |
| Dr.22828.1.A1_at | hydroxysteroid (17-beta) dehydrogenase 10 | hsd17b10 |
| Dr.10742.2.S1_a_at | "HSPA (heat shock 70kDa) binding protein, cytoplasmic cochaperone 1" | hspbp1 |
| Dr.18265.1.S1_at | id:ibd1172 | id:ibd1172 |
| Dr.7103.1.S1_at | inhibitor of DNA binding 3 | id3 |
| Dr.1636.1.A1_at | interferon-related developmental regulator 2 | ifrd2 |
| Dr.15377.1.S1_at | intraflagellar transport 122 homolog (Chlamydomonas) | ift122 |
| Dr.8145.1.S1_at | insulin-like growth factor 2a | igf2a |
| Dr.8280.1.S1_at | insulin-like growth factor 2 mRNA binding protein 3 | igf2bp3 |
| Dr.8280.1.S2_at | insulin-like growth factor 2 mRNA binding protein 3 | igf2bp3 |
| Dr.8587.1.A1_at | insulin-like growth factor binding protein 1a | igfbp1a |
| Dr.8587.1.A2_at | insulin-like growth factor binding protein 1a | igfbp1a |
| Dr.3939.1.A1_at | im:6903007 | im:6903007 |
| Dr.10170.1.A1_at | im:6908224 | im:6908224 |
| Dr.25631.1.S1_at | "im:6909388 \| radial spoke head 1 homolog \| Rtf1, Paf1/RNA polymerase II complex component, homolog (S. cerevisiae)" | im:6909388 \| LOC100334583 \| rtf1 |
| Dr.19345.1.A1_at | "IMP4, U3 small nucleolar ribonucleoprotein, homolog (yeast)" | imp4 |
| Dr.19345.2.S1_at | "IMP4, U3 small nucleolar ribonucleoprotein, homolog (yeast)" | imp4 |
| Dr.7971.1.A1_at | importin 9 | ipo9 |
| Dr.12308.1.S1_at | interferon regulatory factor 9 | irf9 |
| Dr.1482.1.A1_at | iron-sulfur cluster scaffold homolog (E. coli) b | iscub |
| Dr.7338.1.S1_at | influenza virus NS1A binding protein b | ivns1abpb |
| Dr.12589.1.S1_at | jagged 1b | jag1b |
| Dr.26325.1.A1_at | junctional adhesion molecule 3b | jam3b |
| Dr.10032.1.S1_at | Jun dimerization protein 2 | jdp2 |
| Dr.20639.1.S1_at | KDEL (Lys-Asp-Glu-Leu) containing 1 | kdelc1 |
| Dr.1557.1.S1_at | kinesin family member 11 | kif11 |
| Dr.3076.1.A1_a_at | kinesin family member 15 | kif15 |
| Dr.6662.1.S1_at | kinesin family member 18A | kif18a |
| Dr.18075.1.S1_at | kinesin family member 20Bb | kif20bb |
| Dr.8295.1.S1_at | kinesin family member 23 | kif23 |
| Dr.25399.1.A1_at | kinesin family member 2C | kif2c |
| Dr.18327.1.S1_at | kinesin family member 7 | kif7 |
| Dr.6237.1.S1_at | kinesin family member C1 \| kinesin family member C1-like | kifc1 \| LOC561143 |
| Dr.4616.1.A1_at | lysine-rich nucleolar protein 1 | knop1 |
| Dr.8763.1.A1_at | karyopherin (importin) beta 3 | kpnb3 |

| | | |
|---|---|---|
| Dr.16885.1.A1_at | "serine palmitoyltransferase, long chain base subunit 2a" | sptlc2a |
| Dr.923.1.A1_at | sequestosome 1 | sqstm1 |
| Dr.9914.1.S1_at | "signal recognition particle receptor, B subunit" | srprb |
| Dr.9070.2.A1_at | serine/arginine-rich splicing factor 6a | srsf6a |
| Dr.9070.3.A1_at | serine/arginine-rich splicing factor 6a | srsf6a |
| Dr.3463.1.S1_at | Sjogren syndrome antigen B (autoantigen La) | ssb |
| Dr.395.1.A1_at | "ST8 alpha-N-acetyl-neuraminide alpha-2,8-sialyltransferase 6" | st8sia6 |
| Dr.2778.1.S1_at | syntaxin 4 | stx4 |
| Dr.6399.1.S1_at | suppressor of fused homolog (Drosophila) | sufu |
| Dr.9711.1.A1_at | sulfatase 1 | sulf1 |
| Dr.6641.1.A1_at | "suppressor of var1, 3-like 1 (S. cerevisiae)" | supv3l1 |
| Dr.17081.1.S1_at | surfeit 6 | surf6 |
| Dr.1811.1.S1_at | suppressor of variegation 3-9 homolog 1b | suv39h1b |
| Dr.10125.1.S1_at | TGF-beta activated kinase 1/MAP3K7 binding protein 2 | tab2 |
| Dr.3179.1.A1_at | "transforming, acidic coiled-coil containing protein 3" | tacc3 |
| Dr.3018.1.A1_at | transcription elongation regulator 1a (CA150) | tcerg1a |
| Dr.25530.1.A1_s_at | telomeric repeat binding factor (NIMA-interacting) 1 | terf1 |
| Dr.20000.1.S1_at | "transcription factor A, mitochondrial" | tfam |
| Dr.7899.1.S1_at | THO complex 2 | thoc2 |
| Dr.3077.2.A1_at | THO complex 6 homolog (Drosophila) | thoc6 |
| Dr.13661.1.S1_at | thymocyte nuclear protein 1 | thyn1 |
| Dr.16862.1.A1_at | "TopBP1-interacting, checkpoint, and replication regulator" | ticrr |
| Dr.8747.1.A1_at | tubulointerstitial nephritis antigen-like 1 | tinagl1 |
| Dr.12950.2.S1_a_at | timeless interacting protein | tipin |
| Dr.5123.1.A1_at | tight junction protein 2b (zona occludens 2) | tjp2b |
| Dr.21038.1.S1_at | tight junction protein 3 | tjp3 |
| Dr.7616.1.A1_s_at | transketolase | tkt |
| Dr.3811.1.A1_at | tousled-like kinase 1a | tlk1a |
| Dr.6670.1.A1_at | transmembrane protein 107 | tmem107 |
| Dr.25489.1.S1_at | transmembrane protein 134 | tmem134 |
| Dr.3402.1.A1_at | transmembrane protein 2 | tmem2 |
| Dr.10531.1.S1_a_at | thymopoietin a | tmpoa |
| Dr.26411.2.S1_s_at | "troponin I, skeletal, fast 2a.2 \| troponin I, skeletal, fast 2a.4" | tnni2a.2 \| tnni2a.4 |
| Dr.26411.1.S1_at | "troponin I, skeletal, fast 2a.4" | tnni2a.4 |
| Dr.2195.1.S1_at | transportin 3 | tnpo3 |
| Dr.2710.1.S1_at | target of myb1-like | tom1 |
| Dr.12521.1.S1_at | translocase of outer mitochondrial membrane 20 homolog a (yeast) | tomm20a |
| Dr.5418.1.S1_at | topoisomerase (DNA) II alpha | top2a |
| Dr.17989.1.A1_at | topoisomerase (DNA) II beta | top2b |
| Dr.2052.1.S1_at | tumor protein p53 | tp53 |
| Dr.316.1.A1_at | "tumor protein p53 binding protein, 2" | tp53bp2 |
| Dr.25206.1.S1_at | trophoblast glycoprotein-like | tpbgl |
| Dr.20815.1.S1_at | alpha-tropomyosin | tpma |
| Dr.23293.1.A1_at | tubulin polymerization-promoting protein family member 3 | tppp3 |
| Dr.7278.1.A1_at | transmembrane phosphatase with tensin homology | tpte |
| Dr.10519.1.A1_at | tripartite motif-containing 71 | trim71 |
| Dr.14411.1.S1_at | tRNA selenocysteine 1 associated protein 1a | trnau1apa |
| Dr.16529.1.S1_at | "transient receptor potential cation channel, subfamily V, member 1" | trpv1 |
| Dr.21722.1.A1_at | tetraspanin 17 | tspan17 |
| Dr.5371.1.A1_at | "TSR1, 20S rRNA accumulation, homolog (yeast)" | tsr1 |
| Dr.18300.1.S1_at | "TSR2, 20S rRNA accumulation, homolog (S. cerevisiae)" | tsr2 |
| Dr.18499.1.A1_at | "TSR3, 20S rRNA accumulation, homolog (S. cerevisiae)" | tsr3 |
| Dr.20010.2.A1_at | "tubulin, alpha 8 like" | tuba8l |

| | | |
|---|---|---|
| Dr.12091.1.A1_at | "laminin, alpha 5" | lama5 |
| Dr.4129.1.S1_at | "laminin, beta 1a" | lamb1a |
| Dr.17237.1.S1_at | leucine aminopeptidase 3 | lap3 |
| Dr.7573.1.A1_at | "La ribonucleoprotein domain family, member 1B" | larp1b |
| Dr.1928.1.A1_at | leucyl-tRNA synthetase b | larsb |
| Dr.9238.1.A1_at | "LATS, large tumor suppressor, homolog 1 (Drosophila)" | lats1 |
| Dr.8109.1.S1_at | lymphocyte cytosolic plastin 1 | lcp1 |
| Dr.1831.1.S1_at | lunatic fringe homolog | lfng |
| Dr.16539.1.S1_at | lin-52 homolog (C. elegans) | lin52 |
| Dr.25051.1.S2_at | lamin B1 | lmnb1 |
| Dr.15991.1.S1_at | e3 ubiquitin-protein ligase RNF182-like | LOC100000332 |
| Dr.15595.1.S1_at | schwannomin-interacting protein 1-like | LOC100003436 |
| Dr.18597.1.A1_at | coiled-coil domain containing 83 | LOC100004068 |
| Dr.4413.1.A1_at | uncharacterized LOC100004835 \| si:dkeyp-1a11.3 | LOC100004835 \| si:dkeyp-1a11.3 |
| Dr.18357.1.A1_at | uncharacterized LOC100005596 | LOC100005596 |
| Dr.3804.2.A1_a_at | v-type proton ATPase 116 kDa subunit a isoform 3-like \| zgc:55891 | LOC100147923 \| zgc:55891 |
| Dr.5372.7.A1_at | hairy-related 4.2-like | LOC100148329 |
| Dr.5372.7.A1_x_at | hairy-related 4.2-like | LOC100148329 |
| Dr.25373.1.A1_at | zinc finger protein 84-like \| zinc finger protein 2 homolog \| zinc finger protein 2 homolog \| zgc:173705 \| zgc:173714 \| zgc:174314 | LOC100148574 \| LOC100329629 \| LOC100329790 \| zgc:173705 \| zgc:173714 \| zgc:174314 |
| Dr.850.1.A1_at | Zinc finger protein 774-like | LOC100149164 |
| Dr.22797.1.A1_at | lysosomal-associated protein transmembrane 5 | LOC100151049 |
| Dr.16938.1.A1_at | uncharacterized LOC100151491 | LOC100151491 |
| Dr.8442.1.A1_at | uncharacterized LOC100189617 \| zgc:123008 | LOC100189617 \| zgc:123008 |
| Dr.15071.1.A1_at | uncharacterized LOC100329277 | LOC100329277 |
| Dr.6117.1.S1_at | zinc finger protein 658-like \| zinc finger protein 510-like \| zinc finger protein 569-like \| si:dkey-30f3.2 \| zgc:173702 \| zgc:173706 \| zgc:173709 \| zgc:173716 \| zgc:174690 \| zgc:174702 \| zgc:174703 \| zgc:174704 | LOC100329723 \| LOC100538254 \| LOC557877 \| si:dkey-30f3.2 \| zgc:173702 \| zgc:173706 \| zgc:173709 \| zgc:173716 \| zgc:174690 \| zgc:174702 \| zgc:174703 \| zgc:174704 |
| Dr.1691.1.S1_at | ribonucleoside-diphosphate reductase subunit M2-like \| ribonucleotide reductase M2 polypeptide | LOC100330864 \| rrm2 |
| Dr.6673.1.A1_at | protein Shroom4-like \| shroom family member 4 | LOC100331658 \| shroom4 |
| Dr.12962.1.A1_at | bromo adjacent homology domain-containing 1 protein-like \| monoacylglycerol O-acyltransferase 3a | LOC100331694 \| mogat3a |
| Dr.12145.1.A1_at | PCNA-associated factor-like | LOC100333229 |
| Dr.19013.1.S1_at | proteasome assembly chaperone 4-like | LOC100333944 |

| | | |
|---|---|---|
| Dr.664.1.S1_at | "tubulin, alpha 8 like 4" | tuba8l4 |
| Dr.14735.1.A1_at | taxilin gamma | txlng |
| Dr.18396.1.S1_at | thioredoxin interacting protein a | txnipa |
| Dr.4986.1.S1_at | thioredoxin reductase 1 | txnrd1 |
| Dr.1047.1.S1_at | thymidylate synthase | tyms |
| Dr.3305.1.S1_at | "ubiquitin-conjugating enzyme E2G 1b (UBC7 homolog, yeast)" | ube2g1b |
| Dr.18169.1.S1_at | "ubiquitin-like, containing PHD and RING finger domains, 1 \| ubiquitin-like with PHD and ring finger domains 1" | uhrf1 \| uhrf1 |
| Dr.9809.2.S1_at | "ubiquitin-like, containing PHD and RING finger domains, 1 \| ubiquitin-like with PHD and ring finger domains 1" | uhrf1 \| uhrf1 |
| Dr.12611.1.S1_at | unc-119 homolog 1 | unc119.1 |
| Dr.13796.1.S1_at | UPF3 regulator of nonsense transcripts homolog B (yeast) | upf3b |
| Dr.7852.1.S1_at | "UTP11-like, U3 small nucleolar ribonucleoprotein (yeast)" | utp11l |
| Dr.3262.1.S1_at | "utp15, U3 small nucleolar ribonucleoprotein, homolog" | utp15 |
| Dr.14584.1.S1_at | "UTP23, small subunit (SSU) processome component, homolog (yeast)" | utp23 |
| Dr.15435.1.S1_at | "UTP6, small subunit (SSU) processome component, homolog (yeast)" | utp6 |
| Dr.11692.1.S1_at | --- | VASP |
| Dr.597.1.S2_at | vascular endothelial growth factor Aa | vegfaa |
| Dr.15781.1.S1_at | vaccinia related kinase 2 | vrk2 |
| Dr.9655.1.A1_at | vessel-specific 1 | vsg1 |
| Dr.6157.1.S1_at | V-set and immunoglobulin domain containing 10 | vsig10 |
| Dr.546.1.S1_at | visual system homeobox 2 | vsx2 |
| Dr.2528.2.S1_at | WD repeat domain 3 | wdr3 |
| Dr.3743.1.S1_at | WD repeat domain 43 | wdr43 |
| Dr.4658.1.S1_at | WD repeat domain 46 | wdr46 |
| Dr.16864.1.A1_at | WD repeat domain 74 | wdr74 |
| Dr.16984.1.A1_s_at | WD repeat domain 92 | wdr92 |
| Dr.2130.1.A1_at | Wolf-Hirschhorn syndrome candidate 1 | whsc1 |
| Dr.25142.1.S1_at | "WD repeat containing, antisense to TP73" | wrap73 |
| Dr.842.1.A1_at | wu:fa55b05 | wu:fa55b05 |
| Dr.5708.1.A1_at | wu:fb10b07 | wu:fb10b07 |
| Dr.23570.1.A1_at | wu:fb13b10 | wu:fb13b10 |

| | | |
|---|---|---|
| Dr.13772.1.A1_at | "zinc finger protein 36, C3H1 type-like 1-like" | LOC100334443 |
| Dr.15136.1.A1_at | "type II inositol-1,4,5-trisphosphate 5-phosphatase-like \| type II inositol-1,4,5-trisphosphate 5-phosphatase-like" | LOC100534683 \| LOC563543 |
| Dr.236.1.S1_at | apoptosis-enhancing nuclease-like | LOC100534720 |
| Dr.6127.1.A1_at | "tyrosyl-tRNA synthetase, mitochondrial-like \| tyrosyl-tRNA synthetase 2, mitochondrial" | LOC100534736 \| yars2 |
| Dr.23733.1.S1_at | UBX domain-containing protein 1-like \| UBX domain protein 1 | LOC100534742 \| ubxn1 |
| Dr.1511.1.A1_at | zinc finger protein 135-like | LOC100534888 |
| Dr.16383.1.A1_at | BRCA1-A complex subunit RAP80-like | LOC100535018 |
| Dr.11967.1.A1_at | uncharacterized LOC100535034 \| wu:fe23c11 | LOC100535034 \| wu:fe23c11 |
| Dr.16086.1.S1_at | uncharacterized LOC100535593 \| transcriptional repressor NF-X1-like | LOC100535593 \| LOC562071 |
| Dr.11847.1.S1_s_at | neurogenic locus notch homolog protein 1-like \| notch homolog 1a | LOC100535971 \| notch1a |
| Dr.11189.1.S1_at | uncharacterized LOC100536039 | LOC100536039 |
| Dr.4703.1.S1_at | targeting protein for Xklp2-B-like | LOC100536090 |
| Dr.18257.2.S1_at | zinc finger protein 135-like | LOC100536203 |
| Dr.2076.2.A1_x_at | zinc finger protein 551-like \| gastrula zinc finger protein XlCGF57.1-like \| zgc:173703 \| zgc:173713 | LOC100536334 \| LOC100537105 \| zgc:173703 \| zgc:173713 |
| Dr.23191.1.A1_at | uncharacterized LOC100536508 | LOC100536508 |
| Dr.15040.1.S1_at | tRNA (guanine-N(7)-)-methyltransferase subunit WDR4-like \| WD repeat domain 4 | LOC100536637 \| wdr4 |
| Dr.18321.1.S1_at | "lysosome membrane protein 2-like \| scavenger receptor class B, member 2" | LOC100536792 \| scarb2 |
| Dr.15020.1.A1_at | protein Spindly-like \| zgc:171223 | LOC100537471 \| zgc:171223 |
| Dr.13669.1.A1_at | uncharacterized LOC100537802 \| uncharacterized LOC799918 | LOC100537802 \| LOC799918 |
| Dr.12672.1.A1_at | mitochondrial import inner membrane translocase subunit TIM44-like \| translocase of inner mitochondrial membrane 44 homolog (yeast) | LOC100537910 \| timm44 |
| Dr.15869.1.A1_at | uncharacterized LOC100538063 | LOC100538063 |
| Dr.8554.1.A1_at | v-set domain-containing T-cell activation inhibitor 1-like | LOC100538282 |
| Dr.18233.1.A1_at | kinesin family member 20A | LOC325449 |
| Dr.15915.1.S1_at | G patch domain containing 4-like | LOC555692 |
| Dr.11818.1.A1_at | e3 ubiquitin-protein ligase RNF19A-like | LOC557995 |
| Dr.26197.1.S1_at | e3 ubiquitin-protein ligase RNF19A-like | LOC557995 |
| Dr.26197.2.A1_at | e3 ubiquitin-protein ligase RNF19A-like | LOC557995 |
| Dr.19575.2.A1_at | uncharacterized LOC561947 | LOC561947 |
| Dr.9892.1.A1_at | metallo-beta-lactamase domain-containing protein 1-like | LOC564200 |
| Dr.9892.2.S1_at | metallo-beta-lactamase domain-containing protein 1-like | LOC564200 |
| Dr.17663.1.S1_at | uncharacterized LOC565172 \| si:ch211-207m11.1 | LOC565172 \| si:ch211-207m11.1 |
| Dr.17663.2.A1_at | uncharacterized LOC565172 \| si:ch211-207m11.1 | LOC565172 \| si:ch211-207m11.1 |
| Dr.19839.1.A1_at | jun dimerization protein 2-like | LOC570258 |
| Dr.2062.1.S1_at | serine/threonine-protein kinase VRK1-like \| vaccinia related kinase 1 | LOC794952 \| vrk1 |
| Dr.17373.2.S1_at | uncharacterized LOC796453 | LOC796453 |
| Dr.18151.1.S1_at | uncharacterized LOC798783 | LOC798783 |
| Dr.9300.1.A1_at | protein FAM118B-like | LOC799102 |
| Dr.20533.1.A1_at | leucine rich repeat (in FLII) interacting protein 1a | lrrfip1a |

| | | |
|---|---|---|
| Dr.26403.2.S1_at | wu:fb25b09 | wu:fb25b09 |
| Dr.5095.1.A1_at | wu:fb25b09 | wu:fb25b09 |
| Dr.11316.2.A1_at | wu:fb33e04 | wu:fb33e04 |
| Dr.23688.1.A1_at | wu:fb39e08 | wu:fb39e08 |
| Dr.12478.1.S1_at | wu:fb50h02 | wu:fb50h02 |
| Dr.25060.3.A1_at | wu:fb53f04 | wu:fb53f04 |
| Dr.23700.1.A1_at | wu:fb55h10 | wu:fb55h10 |
| Dr.4268.1.S1_at | wu:fb66f03 | wu:fb66f03 |
| Dr.3390.1.S1_at | wu:fb75e06 | wu:fb75e06 |
| Dr.2076.1.A1_x_at | wu:fb94b09 | wu:fb94b09 |
| Dr.5079.1.S1_at | wu:fc02d02 | wu:fc02d02 |
| Dr.21323.1.A1_at | wu:fc10d12 | wu:fc10d12 |
| Dr.3757.1.A1_at | wu:fc13c09 | wu:fc13c09 |
| Dr.23876.1.A1_at | wu:fc44g11 | wu:fc44g11 |
| Dr.21777.1.A1_at | wu:fc55b02 | wu:fc55b02 |
| Dr.21783.1.S1_at | wu:fc55g01 | wu:fc55g01 |
| Dr.1837.1.A1_at | wu:fc83f05 | wu:fc83f05 |
| Dr.25580.1.S1_at | wu:fd23c12 | wu:fd23c12 |
| DrAffx.2.64.S1_at | wu:fd23d07 | wu:fd23d07 |
| Dr.6885.1.A1_at | wu:fe14d06 | wu:fe14d06 |
| Dr.22459.1.A1_at | wu:fe37d09 | wu:fe37d09 |
| Dr.12338.2.A1_a_at | wu:fi28f11 | wu:fi28f11 |
| Dr.7226.1.S1_at | wu:fj65h10 | wu:fj65h10 |
| Dr.7590.1.A1_at | wu:fj68b05 | wu:fj68b05 |
| Dr.10282.1.A1_at | wu:fk30a05 | wu:fk30a05 |
| Dr.10012.1.A1_at | wu:fk83g11 | wu:fk83g11 |
| Dr.5569.2.A1_at | wu:fl08f01 | wu:fl08f01 |
| Dr.10281.1.S1_at | X-ray repair complementing defective repair in Chinese hamster cells 5 | xrcc5 |
| Dr.2355.1.S1_at | Yes-associated protein 1 | yap1 |
| Dr.4644.1.S1_at | YY1 transcription factor b | yy1b |
| Dr.15097.1.A1_at | zinc finger and BTB domain containing 22b | zbtb22b |
| Dr.24762.1.S1_at | zinc finger CCCH-type containing 15 | zc3h15 |
| Dr.6559.1.A1_at | zinc finger CCCH-type containing 7B | zc3h7b |
| Dr.13742.1.S1_at | "zinc finger, CCHC domain containing 9" | zcchc9 |
| Dr.17192.1.S1_at | zgc:100870 | zgc:100870 |
| Dr.26532.1.A1_at | zgc:101819 | zgc:101819 |
| Dr.20417.1.S1_at | zgc:101858 | zgc:101858 |
| Dr.7250.1.A1_at | zgc:110239 | zgc:110239 |

| | | | | | |
|---|---|---|---|---|---|
| Dr.24292.3.A1_a_at | leucine rich repeat neuronal 1 | lrrn1 | Dr.24858.1.S1_at | zgc:110540 | zgc:110540 |
| Dr.3707.1.A1_at | large subunit GTPase 1 homolog (S. cerevisiae) | lsg1 | Dr.5959.1.A1_at | zgc:112020 | zgc:112020 |
| Dr.15890.1.A1_at | LSM12 homolog a (S. cerevisiae) | lsm12a | Dr.5167.1.A1_at | zgc:114181 | zgc:114181 |
| DrAffx.1.53.S1_at | Ly1 antibody reactive homolog (mouse) | lyar | Dr.7787.1.S1_at | zgc:136826 | zgc:136826 |
| Dr.15000.1.S1_at | lysophospholipase I | lypla1 | Dr.11561.1.A1_at | zgc:152651 | zgc:152651 |
| Dr.4833.2.S1_at | MAD2 mitotic arrest deficient-like 1 (yeast) | mad2l1 | Dr.3512.1.A1_at | zgc:152830 | zgc:152830 |
| Dr.4000.1.S1_at | mitogen-activated protein kinase kinase kinase kinase 5 | map4k5 | Dr.12649.1.S1_at | zgc:153041 | zgc:153041 |
| Dr.7930.1.S1_at | mitogen-activated protein kinase 14a | mapk14a | Dr.20487.1.S1_at | zgc:153115 | zgc:153115 |
| Dr.20395.1.A1_at | mitogen-activated protein kinase-activated protein kinase 2a | mapkapk2a | Dr.13693.1.S1_at | zgc:153377 | zgc:153377 |
| Dr.10840.1.S1_at | "microtubule-associated protein, RP/EB family, member 1a" | mapre1a | Dr.15964.1.A1_at | zgc:158292 | zgc:158292 |
| Dr.26268.1.A1_at | --- | MARCO | Dr.23447.1.A1_at | zgc:158345 | zgc:158345 |
| Dr.3498.1.S1_at | "methionine adenosyltransferase I, alpha" | mat1a | Dr.5407.1.A1_at | zgc:158363 | zgc:158363 |
| Dr.20010.6.A1_at | methylcrotonoyl-Coenzyme A carboxylase 2 (beta) | mccc2 | Dr.19956.1.S1_at | zgc:158366 | zgc:158366 |
| Dr.784.1.S1_at | MCM3 minichromosome maintenance deficient 3 (S. cerevisiae) | mcm3 | Dr.1992.1.S1_at | zgc:158399 | zgc:158399 |
| Dr.5091.1.S1_at | "MCM4 minichromosome maintenance deficient 4, mitotin (S. cerevisiae)" | mcm4 | Dr.23944.1.S1_at | zgc:162082 | zgc:162082 |
| Dr.1055.1.S1_at | MCM7 minichromosome maintenance deficient 7 (S. cerevisiae) | mcm7 | Dr.8643.1.A1_x_at | zgc:162193 | zgc:162193 |
| Dr.4989.1.S1_at | minichromosome maintenance complex binding protein | mcmbp | Dr.4334.1.A1_at | zgc:162967 | zgc:162967 |
| Dr.542.1.S1_at | transformed 3T3 cell double minute 2 homolog (mouse) | mdm2 | Dr.2039.1.A1_at | zgc:165515 | zgc:165515 |
| Dr.18106.1.S1_at | maternal embryonic leucine zipper kinase | melk | Dr.4868.1.A1_at | zgc:174888 | zgc:174888 |
| Dr.18296.1.S1_at | mind bomb | mib | Dr.6620.1.A1_at | zgc:174890 | zgc:174890 |
| Dr.8283.1.S1_at | muscle-specific beta 1 integrin binding protein | mibp | Dr.2985.1.A1_at | zgc:175096 | zgc:175096 |
| Dr.19450.1.S1_at | mical-like 2b | micall2b | Dr.11429.1.A1_at | zgc:175133 | zgc:175133 |
| Dr.6498.1.S1_at | mesoderm induction early response 1 homolog a (Xenopus laevis) | mier1a | Dr.15851.1.A1_at | zgc:193690 | zgc:193690 |
| Dr.26464.1.S1_at | MYC induced nuclear antigen-like | minal | Dr.26453.1.S1_at | zgc:64116 | zgc:64116 |
| Dr.13195.1.A1_at | "mutL homolog 1, colon cancer, nonpolyposis type 2 (E. coli)" | mlh1 | Dr.1863.1.A1_at | zgc:77086 | zgc:77086 |
| Dr.1996.1.A1_at | MAX-like protein X | mlx | Dr.12453.1.A1_at | zgc:86764 | zgc:86764 |
| Dr.2408.1.A1_at | matrix metalloproteinase 2 | mmp2 | Dr.9464.1.A1_at | zgc:92107 | zgc:92107 |
| Dr.5285.1.A1_at | "MMS22-like, DNA repair protein" | mms22l | Dr.14880.1.S1_at | zinc finger protein 330 | znf330 |
| Dr.13950.1.S1_at | mortality factor 4 like 1 | morf4l1 | Dr.11710.1.A1_at | zinc finger protein 532 | znf532 |
| Dr.7116.1.S1_at | M-phase phosphoprotein 10 (U3 small nucleolar ribonucleoprotein) | mphosph10 | Dr.3201.1.S1_at | zinc finger-like gene 2a | znfl2a |
| Dr.13254.1.A1_at | "membrane protein, palmitoylated 1" | mpp1 | Dr.4773.1.A1_at | "Zwilch, kinetochore associated, homolog (Drosophila)" | zwilch |
| Dr.1842.1.A1_at | "membrane protein, palmitoylated 1" | mpp1 | Dr.3891.1.A1_at | septin 6 | 5-Sep |
| Dr.16031.1.S1_at | mitochondrial ribosomal protein L1 | mrpl1 | Dr.10904.3.A1_at | --- | --- |
| Dr.9805.1.S1_at | mitochondrial ribosomal protein L12 | mrpl12 | Dr.11610.1.S1_at | --- | --- |
| Dr.18076.1.A1_at | mitochondrial ribosomal protein L43 | mrpl43 | Dr.13336.1.S1_at | --- | --- |
| Dr.7183.1.S1_at | mitochondrial ribosomal protein L45 | mrpl45 | Dr.13336.2.S1_a_at | --- | --- |
| Dr.18075.3.A1_s_at | mitochondrial ribosomal protein S18B | mrps18b | Dr.13336.2.S1_at | --- | --- |
| Dr.15402.1.A1_a_at | mitochondrial ribosomal protein S5 | mrps5 | Dr.14236.1.A1_at | --- | --- |
| Dr.15068.1.S1_at | mitochondrial ribosome recycling factor | mrrf | Dr.15231.1.A1_at | --- | --- |
| Dr.4274.2.A1_at | mRNA turnover 4 homolog (S. cerevisiae) | mrto4 | Dr.15497.1.A1_at | --- | --- |
| Dr.4108.1.S1_at | mutS homolog 6 (E. coli) | msh6 | Dr.16254.1.A1_at | --- | --- |
| Dr.20552.1.S1_at | moesin a | msna | Dr.17413.1.A1_at | --- | --- |
| Dr.24940.1.S1_at | moesin a | msna | Dr.17716.1.S1_at | --- | --- |
| Dr.7124.1.S1_at | methylenetetrahydrofolate dehydrogenase (NADP+ dependent) 1b | mthfd1b | Dr.17717.1.A1_at | --- | --- |
| Dr.14691.1.A1_at | metaxin 1b | mtx1b | Dr.18221.1.A1_at | --- | --- |
| Dr.977.2.A1_at | MUS81 endonuclease homolog (yeast) | mus81 | Dr.19431.1.A1_at | --- | --- |
| Dr.19951.1.S1_a_at | MYB binding protein (P160) 1a | mybbp1a | Dr.19532.1.A1_at | --- | --- |
| Dr.25251.1.A1_at | MYB binding protein (P160) 1a | mybbp1a | Dr.19743.1.A1_at | --- | --- |
| Dr.17421.1.A1_at | myeloblastosis oncogene-like 2 | mybl2 | Dr.20010.14.S1_at | --- | --- |
| Dr.21800.1.A1_at | "myosin binding protein C, fast type b" | mybpc2b | Dr.20521.1.A1_at | --- | --- |
| Dr.1.1.S1_at | myelocytomatosis oncogene a | myca | Dr.21280.1.S1_at | --- | --- |
| Dr.9142.1.S1_at | "myosin, heavy chain 10, non-muscle" | myh10 | Dr.21848.1.A1_at | --- | --- |
| Dr.2601.1.S1_at | "N(alpha)-acetyltransferase 15, NatA auxiliary subunit a" | naa15a | Dr.22145.1.A1_at | --- | --- |
| Dr.21461.1.A1_at | nuclear assembly factor 1 homolog (S. cerevisiae) | naf1 | Dr.23212.1.A1_at | --- | --- |
| Dr.23775.1.A1_at | nucleosome assembly protein 1-like 4a | nap1l4a | Dr.23223.1.A1_at | --- | --- |
| Dr.20947.2.A1_a_at | nuclear autoantigenic sperm protein (histone-binding) | nasp | Dr.25403.1.A1_at | --- | --- |
| Dr.17557.1.S1_at | neurocalcin delta a | ncalda | Dr.25672.1.A1_at | --- | --- |
| Dr.25645.1.S1_at | neurocalcin delta a | ncalda | Dr.25683.10.S1_at | --- | --- |
| Dr.9196.1.S1_at | "non-SMC condensin II complex, subunit D3" | ncapd3 | Dr.25703.1.A1_at | --- | --- |
| Dr.15653.1.S1_at | "non-SMC condensin I complex, subunit G" | ncapg | Dr.26056.1.A1_s_at | --- | --- |
| Dr.17297.1.A1_at | "non-SMC condensin II complex, subunit G2" | ncapg2 | Dr.26490.2.A1_at | --- | --- |

| | | | | | | |
|---|---|---|---|---|---|---|
| Dr.3958.1.A1_at | "non-SMC condensin I complex, subunit H" | ncaph | | Dr.3021.1.S1_at | --- | --- |
| Dr.25587.1.A1_at | NDC1 transmembrane nucleoporin | ndc1 | | Dr.337.1.A1_at | --- | --- |
| Dr.1175.1.A1_at | "NADH dehydrogenase (ubiquinone) 1 alpha subcomplex, 10" | ndufa10 | | Dr.3862.1.A1_at | --- | --- |
| Dr.20071.1.S1_at | nei endonuclease VIII-like 1 (E. coli) | neil1 | | Dr.3862.2.S1_at | --- | --- |
| Dr.9529.1.A1_at | nuclear factor (erythroid-derived 2)-like 3 | nfe2l3 | | Dr.42.1.A1_a_at | --- | --- |
| Dr.26514.1.A1_at | NFS1 nitrogen fixation 1 (S. cerevisiae) | nfs1 | | Dr.42.3.A1_x_at | --- | --- |
| Dr.20137.1.S1_at | "neuroguidin, EIF4E binding protein" | ngdn | | Dr.7594.1.A1_at | --- | --- |
| Dr.20137.2.A1_at | "neuroguidin, EIF4E binding protein" | ngdn | | DrAffx.2.105.S1_at | --- | --- |

**Supplemental Figure 4: List of SAM positive genes**

Knockdown of nhsb causes changes in gene expression in the retina. SAM analysis identified significanly upregulated genes in nhsb MO injected embryos compared to control

### SAM negative genes

| Probe ID | Gene Tile | Gene Symbol |
|---|---|---|
| Dr.8142.1.S1_at | arylalkylamine N-acetyltransferase | aanat2 |
| Dr.24913.1.A1_a_at | "ATP-binding cassette, sub-family C (CFTR/MRP), member 2" | abcc2 |
| Dr.25387.1.A1_at | acid phosphatase-like 2 | acpl2 |
| Dr.185.1.A1_at | acyl-CoA synthetase long-chain family member 6 | acsl6 |
| Dr.6517.1.S1_at | adenylosuccinate synthase like 1 | adssl1 |
| Dr.17696.1.A1_at | A kinase (PRKA) anchor protein 11 | akap11 |
| Dr.12901.1.A1_at | A kinase (PRKA) anchor protein 6 | akap6 |
| Dr.8180.1.S1_at | "aminolevulinate, delta-, synthetase 2" | alas2 |
| Dr.4277.1.S1_at | "ankylosis, progressive homolog b" | ankhb |
| Dr.7290.1.A1_at | ankyrin repeat domain 12 | ankrd12 |
| Dr.1190.1.S1_at | annexin A1b | anxa1b |
| Dr.1442.2.S2_a_at | amyloid beta (A4) precursor protein a | appa |
| Dr.2615.1.S1_at | amyloid beta (A4) precursor protein b | appb |
| Dr.5865.1.A1_at | ADP-ribosylation factor interacting protein 2a | arfip2a |
| Dr.12695.1.A1_at | Rho GTPase activating protein 4b | arhgap4b |
| Dr.9845.2.A1_at | "ADP-ribosylation factor-like 3, like 2" | arl3l2 |
| Dr.18638.1.S1_at | ADP-ribosylation factor-like 6 | arl6 |
| Dr.7450.1.A1_at | ADP-ribosylation factor-like 8A | arl8a |
| Dr.17322.1.A1_at | --- | ASH1L |
| Dr.14729.1.S1_at | astrotactin 1 | astn1 |
| Dr.14729.3.S1_a_at | astrotactin 1 | astn1 |
| Dr.15962.1.A1_at | ATG12 autophagy related 12 homolog (S. cerevisiae) | atg12 |
| Dr.18987.1.S1_at | ATG9 autophagy related 9 homolog A (S. cerevisiae) | atg9a |
| Dr.4686.1.S1_at | "ATPase, Na+/K+ transporting, beta 3b polypeptide" | atp1b3b |
| Dr.15256.1.A1_at | "ATP synthase, H+ transporting, mitochondrial F0 complex, subunit b, isoform 1" | atp5f1 |
| Dr.10576.2.A1_at | "ATPase, H+ transporting, lysosomal, V0 subunit c, b" | atp6v0cb |
| Dr.2032.1.A1_at | "ATPase, H+ transporting, V0 subunit D isoform 1" | atp6v0d1 |
| Dr.5981.1.S1_at | "ATPase, H+ transporting, lysosomal V1 subunit B2" | atp6v1b2 |
| Dr.7966.1.S1_at | "ATPase, H+ transporting, lysosomal, V1 subunit E isoform 1b" | atp6v1e1b |
| Dr.370.1.S1_at | "ATPase, H+ transporting, V1 subunit G isoform 1" | atp6v1g1 |
| Dr.7421.1.S1_at | "ATPase, H+ transporting, lysosomal, V1 subunit H" | atp6v1h |
| Dr.11059.1.S1_at | "ATPase, H+ transporting V0 subunit e2" | atpv0e2 |
| Dr.20512.1.A1_at | ataxin 7-like 3 | atxn7l3 |
| Dr.7804.1.S1_at | antizyme inhibitor 1b | azin1b |
| Dr.25155.1.S1_s_at | "ba1 globin \| ba1 globin, like \| ba2 globin" | ba1 \| ba1l \| ba2 |
| Dr.17142.1.A1_at | beta-site APP-cleaving enzyme 1 | bace1 |
| Dr.6278.1.A1_at | BCL2-associated athanogene 4 | bag4 |
| Dr.20665.1.S1_at | Bardet-Biedl syndrome 5 | bbs5 |
| Dr.14345.1.A1_at | Bardet-Biedl syndrome 7 | bbs7 |
| Dr.7959.1.A1_at | B-cell receptor-associated protein 31 | bcap31 |
| Dr.6069.1.S1_at | "branched chain aminotransferase 1, cytosolic" | bcat1 |
| Dr.13843.1.S1_at | "basic helix-loop-helix family, member e22" | bhlhe22 |
| Dr.723.1.A1_at | "biogenesis of lysosomal organelles complex-1, subunit 4, cappuccino" | bloc1s4 |
| Dr.11793.1.A1_at | BCL2/adenovirus E1B interacting protein 3 | bnip3 |
| Dr.12862.1.A1_at | brain specific kinase 146 \| brain specific kinase 146-like | bsk146 \| LOC791840 |

| Probe ID | Gene Tile | Gene Symbol |
|---|---|---|
| Dr.6811.1.A1_at | "ras-related C3 botulinum toxin substrate 3a (rho family, small GTP binding protein Rac3)" | rac3a |
| Dr.26323.1.A1_at | RAP1 GTPase activating protein | rap1gap |
| Dr.4032.1.A1_at | Rap guanine nucleotide exchange factor (GEF) 2 | rapgef2 |
| Dr.12216.1.A1_at | RAS guanyl releasing protein 4 | rasgrp4 |
| Dr.15195.1.A1_at | "RNA binding protein, fox-1 homolog (C. elegans) 1" | rbfox1 |
| Dr.16723.1.A1_at | "RNA binding protein, fox-1 homolog (C. elegans) 1" | rbfox1 |
| Dr.5878.1.A1_at | rabconnectin 3 | rc3 |
| Dr.11063.1.A1_at | --- | RCAN1B |
| Dr.1817.1.A1_at | regulator of calcineurin family member 3 | rcan3 |
| Dr.15089.1.S1_at | rad and gem related GTP binding protein 1 | rem1 |
| Dr.12444.1.A1_at | regulatory factor X-associated protein | rfxap |
| Dr.17500.1.A1_at | RGP1 retrograde golgi transport homolog (S. cerevisiae) | rgp1 |
| Dr.14615.1.A1_at | regulator of G-protein signaling 17 | rgs17 |
| Dr.13030.1.A1_at | regulator of G-protein signaling 5a | rgs5a |
| Dr.354.1.S1_at | rhodopsin | rho |
| Dr.11062.1.A1_at | resistance to inhibitors of cholinesterase 8 homolog A | ric8a |
| Dr.13795.1.A1_at | relaxin 3a | rln3a |
| Dr.12666.1.S1_at | ring finger protein 146 | rnf146 |
| Dr.19063.1.S1_at | ring finger protein 175 | rnf175 |
| Dr.11667.1.S1_at | ring finger protein 41 | rnf41 |
| Dr.18132.1.A1_at | ring finger protein 44 | rnf44 |
| Dr.9906.1.S1_at | retinal outer segment membrane protein 1b | rom1b |
| Dr.17137.1.S1_at | "RAR-related orphan receptor A, paralog b" | rorab |
| Dr.26511.1.S1_at | ribosomal protein L22-like 1 | rpl22l1 |
| Dr.25441.1.S1_at | "ribosomal protein, large P2" | rplp2 |
| Dr.24633.1.S1_at | "ribosomal protein S27, isoform 1" | rps27.1 |
| Dr.1208.1.S1_at | ribosomal L24 domain containing 1 | rsl24d1 |
| Dr.11076.1.A1_at | "runt-related transcription factor 1; translocated to, 1 (cyclin D-related)" | runx1t1 |
| Dr.540.1.S1_at | retinal homeobox gene 3 | rx3 |
| Dr.15019.1.S1_at | RING1 and YY1 binding protein a | rybpa |
| Dr.919.1.A1_at | "SAM domain, SH3 domain and nuclear localisation signals, 1a" | samsn1a |
| Dr.6709.1.S1_at | sb:cb252 | sb:cb252 |
| Dr.15351.1.S1_at | "sema domain, immunoglobulin domain (Ig), transmembrane domain (TM) and short cytoplasmic domain, (semaphorin) 4Ba" | sema4ba |
| Dr.23228.1.A1_at | septin 8b | sept8b |
| Dr.7467.1.S1_at | serine incorporator 5 | serinc5 |
| Dr.2720.1.A1_at | "sarcoglycan, epsilon" | sgce |
| Dr.17453.1.A1_at | SH2 domain containing 3Cb | sh2d3cb |
| Dr.7417.1.S1_at | SH3-domain binding protein 5b (BTK-associated) | sh3bp5b |
| Dr.12559.1.S1_at | si:ch211-11c15.3 | si:ch211-11c15.3 |
| Dr.7912.1.A1_at | si:ch211-147a11.3 | si:ch211-147a11.3 |
| Dr.17180.1.A1_at | si:ch211-181b18.1 | si:ch211-181b18.1 |
| Dr.3636.1.A1_at | si:ch211-194k22.8 | si:ch211-194k22.8 |
| Dr.16426.1.A1_at | si:ch211-197g15.12 | si:ch211-197g15.12 |
| Dr.15919.1.A1_at | si:ch211-239e6.4 | si:ch211-239e6.4 |
| Dr.23244.1.A1_at | si:ch211-248g20.5 | si:ch211-248g20.5 |

| | | | | | |
|---|---|---|---|---|---|
| Dr.25528.1.A1_at | BTB (POZ) domain containing 6b | btbd6b | Dr.9844.1.A1_at | si:ch211-263m18.3 | si:ch211-263m18.3 |
| Dr.7931.1.A1_at | "core 1 synthase, glycoprotein-N-acetylgalactosamine 3-beta-galactosyltransferase, 1a" | c1galt1a | Dr.2877.1.A1_at | si:ch211-76l23.4 | si:ch211-76l23.4 |
| Dr.14041.1.S1_at | "complement component 1, q subcomponent, A chain \| complement component 1, q subcomponent, C chain" | c1qa \| c1qc | Dr.14865.1.A1_at | si:ch73-52e5.2 | si:ch73-52e5.2 |
| Dr.16848.1.A1_at | C1q and tumor necrosis factor related protein 4 | c1qtnf4 | Dr.15375.1.A1_at | --- | SI:DKEY-114C15.7 |
| Dr.13585.1.A1_at | carbonic anhydrase Xa | ca10a | Dr.22835.1.A1_at | si:dkey-12h9.11 | si:dkey-12h9.11 |
| Dr.13585.2.S1_at | carbonic anhydrase Xa | ca10a | Dr.21395.1.A1_at | --- | SI:DKEY-172J4.3 |
| Dr.21636.1.A1_at | calcium channel flower domain containing 1 | cacfd1 | Dr.24069.1.A1_at | si:dkey-174e3.3 | si:dkey-174e3.3 |
| Dr.15379.1.A1_at | "calcium channel, voltage-dependent, beta 1 subunit" | cacnb1 | Dr.19960.1.A1_at | si:dkey-229p15.1 | si:dkey-229p15.1 |
| Dr.13791.1.A1_at | cell adhesion molecule 2a | cadm2a | Dr.24055.1.S1_at | si:dkey-25o1.6 | si:dkey-25o1.6 |
| Dr.450.1.S1_at | carbonic anhydrase | cahz | Dr.24143.1.A1_at | si:dkey-6e12.4 | si:dkey-6e12.4 |
| Dr.22781.1.A1_at | calbindin 1 | calb1 | Dr.5973.1.A1_at | si:dkeyp-110c7.1 | si:dkeyp-110c7.1 |
| Dr.7048.1.A1_at | calbindin 2a | calb2a | Dr.2124.1.A1_at | si:dkeyp-84f11.5 | si:dkeyp-84f11.5 |
| Dr.11068.1.A1_at | calbindin 2b | calb2b | Dr.616.1.S1_at | sine oculis homeobox homolog 3a | six3a |
| Dr.7908.1.S1_at | "calmodulin 1a \| calmodulin 1b \| calmodulin 2a (phosphorylase kinase, delta) \| calmodulin 2b, (phosphorylase kinase, delta) \| calmodulin 3a (phosphorylase kinase, delta) \| calmodulin 3b (phosphorylase kinase, delta)" | calm1a \| calm1b \| calm2a \| calm2b \| calm3a \| calm3b | Dr.6052.1.A1_at | "SLAIN motif family, member 1a" | slain1a |
| Dr.7908.1.S2_at | "calmodulin 1a \| calmodulin 1b \| calmodulin 2a (phosphorylase kinase, delta) \| calmodulin 2b, (phosphorylase kinase, delta) \| calmodulin 3a (phosphorylase kinase, delta) \| calmodulin 3b (phosphorylase kinase, delta)" | calm1a \| calm1b \| calm2a \| calm2b \| calm3a \| calm3b | Dr.26139.1.A1_at | "solute carrier family 25 (mitochondrial carrier, brain), member 14" | slc25a14 |
| Dr.7638.1.S2_at | "calmodulin 3a (phosphorylase kinase, delta)" | calm3a | Dr.26139.1.A1_x_at | "solute carrier family 25 (mitochondrial carrier, brain), member 14" | slc25a14 |
| Dr.14581.1.A1_at | calcium/calmodulin-dependent protein kinase (CaM kinase) II alpha | camk2a | Dr.21187.1.A1_at | "solute carrier family 2 (facilitated glucose transporter), member 3a" | slc2a3a |
| Dr.23041.1.A1_at | calcium/calmodulin-dependent protein kinase (CaM kinase) II beta 1 | camk2b1 | Dr.16728.1.A1_at | "solute carrier family 32 (GABA vesicular transporter), member 1" | slc32a1 |
| Dr.12561.1.A1_at | calcium/calmodulin-dependent protein kinase (CaM kinase) II delta 2 | camk2d2 | Dr.22804.1.S1_at | "solute carrier family 6, member 17" | slc6a17 |
| Dr.25517.1.S1_at | calcium/calmodulin-dependent protein kinase (CaM kinase) II delta 2 | camk2d2 | Dr.22709.1.S1_at | "solute carrier family 6 (neurotransmitter transporter, GABA), member 1a" | slc6a1a |
| Dr.5419.1.S1_at | "capping protein (actin filament) muscle Z-line, alpha 1a" | capza1a | Dr.25448.1.A1_at | "solute carrier family 6 (neurotransmitter transporter, GABA), member 1b" | slc6a1b |
| Dr.14824.1.S1_at | CAS1 domain containing 1 | casd1 | Dr.9900.1.A1_at | "solute carrier family 9 (sodium/hydrogen exchanger), member 7" | slc9a7 |
| Dr.22832.1.A1_at | cerebellin 1 precursor | cbln1 | Dr.9918.1.S1_at | sarcolemma associated protein b | slmapb |
| Dr.13908.2.S1_a_at | chromobox homolog 1b (HP1 beta homolog Drosophila) | cbx1b | Dr.1867.1.S1_at | spermine synthase | sms |
| Dr.12921.1.A1_at | coiled-coil domain containing 82 | ccdc82 | Dr.4117.1.A1_at | SMAD specific E3 ubiquitin protein ligase 1 | smurf1 |
| Dr.10051.1.A1_at | cyclin G2 | ccng2 | Dr.7815.1.S1_at | synaptosome-associated protein 25a | snap25a |
| Dr.20083.1.A1_at | cyclin G2 | ccng2 | Dr.7598.1.S1_at | synaptosome-associated protein 25b | snap25b |
| Dr.10301.1.A1_at | "CD82 antigen, b" | cd82b | Dr.14576.1.S1_at | "synuclein, gamma b (breast cancer-specific protein 1)" | sncgb |
| Dr.26344.3.S1_a_at | cell division cycle 42 | cdc42 | Dr.25550.1.A1_at | "synuclein, gamma b (breast cancer-specific protein 1)" | sncgb |
| Dr.26344.2.S1_a_at | cell division cycle 42 | cdc42 | Dr.14576.2.S1_at | "synuclein, gamma b (breast cancer-specific protein 1)" | sncgb |
| Dr.1816.1.A1_at | CDC42 effector protein (Rho GTPase binding) 3 | cdc42ep3 | Dr.16935.1.A1_a_at | heme-binding protein soul4 | soul4 |
| Dr.11209.1.A1_at | "cadherin 18, type 2" | cdh18 | Dr.4007.1.S1_at | Sp5 transcription factor-like | sp5l |
| Dr.12605.1.S1_at | cyclin-dependent kinase inhibitor 1Bb \| cyclin-dependent kinase inhibitor 1B-like | cdkn1bb \| LOC100534692 | Dr.10224.1.S1_at | signal peptide peptidase-like 2 | sppl2 |
| Dr.13284.1.A1_at | choline kinase alpha | chka | Dr.14335.1.A1_at | shadow of prion protein | sprn |
| Dr.26446.1.S1_at | "cholinergic receptor, nicotinic, beta polypeptide 3a" | chrnb3a | Dr.26334.1.A1_at | serine/arginine-rich protein specific kinase 1a | srpk1a |
| Dr.771.1.S1_at | "creatine kinase, mitochondrial 1" | ckmt1 | Dr.13698.1.S1_at | sperm specific antigen 2 | ssfa2 |
| Dr.26433.1.S1_at | CDC-like kinase 4a | clk4a | Dr.17292.1.A1_at | slingshot homolog 2b (Drosophila) | ssh2b |
| Dr.11272.1.A1_at | connector enhancer of kinase suppressor of Ras 2b | cnksr2b | Dr.7560.1.S1_at | "somatostatin 1, tandem duplicate 1" | sst1.1 |
| Dr.16729.1.A1_at | contactin 1b | cntn1b | Dr.12804.1.S1_at | somatostatin 3 | sst3 |

| | | |
|---|---|---|
| Dr.2038.1.S1_s_at | COMM domain containing 3 | commd3 |
| Dr.12760.1.A1_at | cytochrome c oxidase subunit IV isoform 2 | cox4i2 |
| Dr.956.1.S1_at | cytochrome c oxidase subunit VIb polypeptide 1 | cox6b1 |
| Dr.5965.1.A1_at | copine II | cpne2 |
| Dr.12552.1.S1_at | cellular retinoic acid binding protein 1a | crabp1a |
| DrAffx.1.10.S1_at | cysteine rich transmembrane BMP regulator 1 (chordin like) | crim1 |
| Dr.7400.1.A1_at | "casein kinase 1, epsilon" | csnk1e |
| Dr.7605.1.A1_at | cysteine-serine-rich nuclear protein 2 | csrnp2 |
| Dr.7214.1.A1_at | cortexin 2 \| cortexin-2-like | ctxn2 \| LOC100536175 |
| Dr.4925.1.S1_x_at | cytoglobin 1 | cygb1 |
| Dr.4925.1.S1_at | cytoglobin 1 | cygb1 |
| Dr.3413.1.S1_at | dachshund c | dachc |
| Dr.8900.1.S1_at | death associated protein | dap |
| Dr.1542.1.A1_at | "diazepam binding inhibitor (GABA receptor modulator, acyl-CoA binding protein)" | dbi |
| Dr.12983.1.A1_at | dynactin 5 | dctn5 |
| Dr.16505.1.A1_at | "DCN1, defective in cullin neddylation 1, domain containing 4 (S. cerevisiae)" | dcun1d4 |
| Dr.7643.1.S1_at | DENN/MADD domain containing 6B | dennd6b |
| Dr.11107.1.A1_at | desumoylating isopeptidase 1a | desi1a |
| Dr.16422.1.S1_at | "DIRAS family, GTP-binding RAS-like 1a" | diras1a |
| Dr.24163.1.A1_at | dickkopf homolog 2 (Xenopus laevis) | dkk2 |
| Dr.21953.1.A1_at | "DnaJ (Hsp40) homolog, subfamily C, member 27" | dnajc27 |
| Dr.11133.1.A1_at | dihydropyrimidinase-like 5b | dpysl5b |
| Dr.12889.1.S1_at | "dynein, cytoplasmic 1, light intermediate chain 1" | dync1li1 |
| Dr.16716.1.S1_at | "dynein, light chain, LC8-type 2a" | dynll2a |
| Dr.15116.1.S1_at | "dynein, light chain, LC8-type 2a" | dynll2a |
| Dr.25567.1.S1_at | "eukaryotic translation initiation factor 3, subunit M" | eif3m |
| Dr.11824.1.S1_at | "eukaryotic translation initiation factor 4A, isoform 1B" | eif4a1b |
| Dr.1909.1.S1_at | "E74-like factor 3 (ets domain transcription factor, epithelial-specific )" | elf3 |
| Dr.17507.1.S1_at | ER membrane protein complex subunit 3 | emc3 |
| Dr.3027.1.S1_at | "erythrocyte membrane protein band 4.1b (elliptocytosis 1, RH-linked)" | epb41b |
| Dr.14860.1.A1_at | erythrocyte membrane protein band 4.1-like 3a | epb41l3a |
| Dr.26376.1.A1_at | Enah/Vasp-like b | evlb |
| Dr.5648.1.A1_at | Ewing sarcoma breakpoint region 1a | ewsr1a |
| Dr.6814.1.S1_at | "fatty acid binding protein 3, muscle and heart" | fabp3 |
| Dr.9450.1.A1_at | "family with sequence similarity 84, member A" | fam84a |
| Dr.15754.1.A1_at | F-box and leucine-rich repeat protein 2 | fbxl2 |
| Dr.12852.1.S1_at | FK506 binding protein 1b | fkbp1b |
| Dr.587.1.S1_at | forkhead box B1.2 \| forkhead box protein B1-like | foxb1.2 \| LOC100535158 |
| Dr.10341.1.S1_at | frizzled homolog 9b | fzd9b |
| Dr.12474.1.A1_at | "gamma-aminobutyric acid (GABA) A receptor, delta" | gabrd |
| Dr.9917.1.S1_at | glutamate decarboxylase 1b | gad1b |
| Dr.4315.1.A1_at | glycerophosphodiester phosphodiesterase domain containing 1 | gdpd1 |
| Dr.1118.1.A1_at | glucose-fructose oxidoreductase domain containing 2 | gfod2 |
| Dr.6125.1.A1_at | gamma-glutamyltransferase 7 | GGT7 |
| Dr.9899.1.S1_at | "guanine nucleotide binding protein (G protein), alpha transducing activity polypeptide 1" | gnat1 |
| Dr.9903.1.A1_at | "guanine nucleotide binding protein (G protein), beta 5b" | gnb5b |
| Dr.14190.1.A1_at | "guanine nucleotide binding protein (G protein), gamma 13b" | gng13b |

| | | |
|---|---|---|
| Dr.7026.1.A1_at | SSU72 RNA polymerase II CTD phosphatase homolog (S. cerevisiae) | ssu72 |
| Dr.25704.1.A1_at | "ST6 beta-galactosamide alpha-2,6-sialyltranferase 1" | st6gal1 |
| Dr.15262.1.S1_at | serine/threonine kinase 38 like | stk38l |
| Dr.1841.1.A1_at | stathmin-like 2a | stmn2a |
| Dr.613.1.S1_at | stomatin | stom |
| Dr.25232.1.A1_at | spermatid perinuclear RNA binding protein | strbp |
| Dr.12806.1.A1_at | synapsin IIb | syn2b |
| Dr.7859.1.A1_at | "spectrin repeat containing, nuclear envelope 1b" | syne1b |
| Dr.11716.1.S1_at | synaptophysin b | sypb |
| Dr.25455.1.A1_at | synaptophysin b | sypb |
| Dr.9125.1.S1_at | synaptotagmin XIa | syt11a |
| Dr.13868.1.S1_at | synaptotagmin IV | syt4 |
| Dr.22094.1.A1_at | T-box 3a | tbx3a |
| Dr.12206.2.S1_at | "transcription elongation factor B (SIII), polypeptide 2 (18kD, elongin B)" | tceb2 |
| Dr.25330.1.A1_at | tectonin beta-propeller repeat containing 2 | tecpr2 |
| Dr.13696.1.S1_at | testis expressed 2 | tex2 |
| Dr.11698.1.S1_at | transcription factor AP-2 gamma (activating enhancer binding protein 2 gamma) | tfap2c |
| Dr.16570.2.S1_at | THAP domain containing 4 | thap4 |
| Dr.18844.1.S1_at | "T-cell leukemia, homeobox 1" | tlx1 |
| Dr.14234.2.A1_at | transmembrane emp24 protein transport domain containing 4 | tmed4 |
| Dr.17164.1.A1_at | transmembrane protein 178 | tmem178 |
| Dr.9292.1.A1_at | transmembrane protein 237b | tmem237b |
| Dr.6796.1.A1_at | transmembrane protein 59-like | tmem59l |
| Dr.17560.1.S1_at | transmembrane protein 9 | tmem9 |
| Dr.17246.1.A1_at | tenomodulin | tnmd |
| DrAffx.1.4.S1_at | "tenascin R (restrictin, janusin)" | tnr |
| Dr.8875.1.S1_at | toll interacting protein | tollip |
| Dr.11206.1.S1_at | triosephosphate isomerase 1a | tpi1a |
| Dr.4157.1.S1_at | triosephosphate isomerase 1b | tpi1b |
| Dr.9221.1.S1_at | transformer 2 beta homolog (Drosophila) | tra2b |
| Dr.5637.1.S1_at | tripartite motif-containing 8 | trim8 |
| Dr.12050.1.A1_at | tripartite motif-containing 9 | trim9 |
| Dr.18524.1.A1_at | tetraspanin 13b | tspan13b |
| Dr.13408.1.S1_at | tweety homolog 3b (Drosophila) | ttyh3b |
| Dr.22554.1.A1_at | tubby homolog (mouse) | tub |
| Dr.12544.1.A1_at | tumor suppressor candidate 3 | tusc3 |
| Dr.24972.1.S1_at | ubiquitin-conjugating enzyme E2Q family member 2 | ube2q2 |
| Dr.17859.1.S1_at | ubiquitin-like domain containing CTD phosphatase 1 | ublcp1 |
| Dr.8724.1.S1_at | ubiquitin carboxyl-terminal esterase L1 (ubiquitin thiolesterase) | uchl1 |
| Dr.15856.1.A1_at | uridine-cytidine kinase 2a | uck2a |
| Dr.15745.1.S1_at | urotensin II-related peptide | urp2 |
| Dr.16861.1.A1_at | upstream transcription factor 1 | usf1 |
| Dr.7095.1.A1_at | vesicle-associated membrane protein 1 | vamp1 |
| Dr.4561.1.S1_at | very low density lipoprotein receptor | vldlr |
| Dr.15874.1.S1_at | vacuolar protein sorting 45 homolog (S. cerevisiae) | vps45 |
| Dr.8802.1.A1_at | WD repeat domain 11 | wdr11 |
| Dr.16768.2.S1_at | tryptophan rich basic protein | wrb |

| | | |
|---|---|---|
| Dr.9876.1.S1_at | "guanine nucleotide binding protein (G protein), gamma transducing activity polypeptide 2a" | gngt2a |
| Dr.6487.1.A1_at | gephyrin b | gphnb |
| Dr.7873.1.S1_at | G protein-coupled receptor 85 | gpr85 |
| Dr.3715.1.A1_at | "gene rich cluster, C10 gene" | grcc10 |
| Dr.18279.1.S1_at | "glutamate receptor, ionotropic, AMPA 2a" | gria2a |
| Dr.14327.1.A1_at | G protein-coupled receptor kinase 1 a | grk1a |
| Dr.22673.1.A1_at | "glutamate receptor, metabotropic 1a" | grm1a |
| Dr.8166.1.S1_at | glycogen synthase kinase 3 beta | gsk3b |
| Dr.4727.1.A1_at | gelsolin b | gsnb |
| Dr.17943.1.A1_at | gelsolin b | gsnb |
| Dr.11305.1.A1_at | guanylate kinase 1b | guk1b |
| Dr.25598.1.A1_at | "H2A histone family, member Y2 \| core histone macro-H2A.2-like" | h2afy2 \| LOC100536875 |
| Dr.8617.1.A1_at | hyaluronan and proteoglycan link protein 1b | hapln1b |
| Dr.1450.1.S1_at | hemoglobin alpha embryonic-3 | hbae3 |
| Dr.1450.1.S1_s_at | hemoglobin alpha embryonic-3 | hbae3 |
| DrAffx.2.15.S1_at | hemoglobin beta embryonic-2 | hbbe2 |
| Dr.14159.1.A1_at | histone deacetylase 9b | hdac9b |
| Dr.13839.1.A1_at | "hepatoma-derived growth factor, related protein 3" | hdgfrp3 |
| Dr.9626.1.A1_at | "homocysteine-inducible, endoplasmic reticulum stress-inducible, ubiquitin-like domain member 1" | herpud1 |
| Dr.26441.1.S1_at | hairy/enhancer-of-split related with YRPW motif-like | heyl |
| Dr.10615.1.A1_at | homeodomain interacting protein kinase 3b | hipk3b |
| Dr.24225.1.S1_at | "heterogeneous nuclear ribonucleoprotein K, like" | hnrpkl |
| Dr.11184.1.A1_at | hippocalcin | hpca |
| Dr.610.2.S1_a_at | "heat shock protein 90, alpha (cytosolic), class A member 1, tandem duplicate 1 \| heat shock protein 90, alpha (cytosolic), class A member 1, tandem duplicate 2" | hsp90aa1.1 \| hsp90aa1.2 |
| Dr.19642.1.A1_at | islet cell autoantigen 1 | ica1 |
| Dr.12836.1.S1_at | "inhibitor of DNA binding 2, dominant negative helix-loop-helix protein, b" | id2b |
| Dr.12836.2.A1_at | "inhibitor of DNA binding 2, dominant negative helix-loop-helix protein, b" | id2b |
| Dr.16287.1.A1_at | "immunoglobin superfamily, member 21b" | igsf21b |
| Dr.17303.1.S1_at | im:7142942 | im:7142942 |
| Dr.5816.1.A1_at | "inhibitor of growth family, member 2" | ing2 |
| Dr.13598.1.A1_at | integrator complex subunit 2 | ints2 |
| Dr.5738.1.S1_at | interphotoreceptor retinoid-binding protein | irbp |
| Dr.25683.6.A1_at | interferon regulatory factor 2 binding protein-like | irf2bpl |
| Dr.12603.1.S1_at | iroquois homeobox protein 7 | irx7 |
| Dr.6200.1.A1_at | integral membrane protein 2Ca | itm2ca |
| Dr.11491.1.S1_at | Josephin domain containing 2 | josd2 |
| Dr.10326.1.S1_at | jun B proto-oncogene a | junba |
| Dr.8131.1.S1_at | Kallmann syndrome 1a sequence | kal1a |
| Dr.17145.1.S1_at | potassium channel tetramerisation domain containing 12.1 | kctd12.1 |
| Dr.11022.1.A1_at | lysine (K)-specific demethylase 5Bb | kdm5bb |
| Dr.9880.1.A1_at | keratocan | kera |
| Dr.18282.7.S1_at | "KH domain containing, RNA binding, signal transduction associated 1a" | khdrbs1a |
| Dr.15083.1.A1_at | kinesin family member 1A | kif1a |
| Dr.12174.1.A1_at | kinesin-associated protein 3a | kifap3a |
| Dr.7499.1.A1_at | kinesin family member C3 | kifc3 |
| Dr.9293.1.A1_at | v-Ki-ras2 Kirsten rat sarcoma viral oncogene homolog | kras |
| Dr.15888.1.A1_at | KxDL motif containing 1 | kxd1 |
| Dr.9200.1.A1_at | lysosomal associated protein transmembrane 4 beta | laptm4b |
| Dr.6975.1.A1_at | "La ribonucleoprotein domain family, member 6" | larp6 |
| Dr.17515.1.A1_at | lactate dehydrogenase Bb | ldhbb |
| Dr.5853.1.A1_at | "lectin, galactoside-binding, soluble, 3 binding protein b" | lgals3bpb |
| Dr.277.1.S1_at | LIM homeobox 1b | lhx1b |
| Dr.1225.1.A1_at | LIM domain kinase 2 | limk2 |
| Dr.11345.1.A1_at | lin-7 homolog A (C. elegans) | lin7a |
| Dr.10254.1.A1_at | lin-7 homolog B (C. elegans) | lin7b |

| | | |
|---|---|---|
| Dr.16768.1.A1_at | tryptophan rich basic protein | wrb |
| Dr.4078.1.A1_at | wu:fb06h03 | wu:fb06h03 |
| Dr.24899.1.A1_at | wu:fb15e04 | wu:fb15e04 |
| Dr.3497.1.A1_at | wu:fb26f10 | wu:fb26f10 |
| Dr.2580.1.A1_at | wu:fc07b10 | wu:fc07b10 |
| Dr.24775.2.A1_at | wu:fc21b07 | wu:fc21b07 |
| Dr.3233.1.A1_at | wu:fc35d04 | wu:fc35d04 |
| Dr.2701.1.A1_at | wu:fc35e07 | wu:fc35e07 |
| Dr.21508.1.A1_at | wu:fc38h03 | wu:fc38h03 |
| Dr.9916.1.A1_at | wu:fc47e09 | wu:fc47e09 |
| Dr.2243.1.A1_at | wu:fc50e08 | wu:fc50e08 |
| Dr.1782.1.A1_at | wu:fc52a02 | wu:fc52a02 |
| Dr.2266.1.A1_at | wu:fc63c11 | wu:fc63c11 |
| Dr.21904.1.A1_at | wu:fc79g11 | wu:fc79g11 |
| Dr.6528.1.A1_at | wu:fd60d11 | wu:fd60d11 |
| Dr.13571.1.A1_at | wu:fi03b02 | wu:fi03b02 |
| Dr.6735.1.A1_at | wu:fj37c02 | wu:fj37c02 |
| Dr.22669.1.A1_at | wu:fj38g04 | wu:fj38g04 |
| Dr.22676.1.A1_at | wu:fj39f02 | wu:fj39f02 |
| Dr.14051.1.A1_at | disintegrin and metalloproteinase domain-containing protein 23 | wu:fj40f01 |
| Dr.6457.1.S1_at | wu:fj40g07 | wu:fj40g07 |
| Dr.22764.1.A1_at | wu:fj51e11 | wu:fj51e11 |
| Dr.22777.1.S1_at | wu:fj53a09 | wu:fj53a09 |
| Dr.5985.1.A1_at | wu:fj54e12 | wu:fj54e12 |
| Dr.13462.1.A1_at | wu:fj55b04 | wu:fj55b04 |
| Dr.11830.1.A1_at | wu:fj55b04 | wu:fj55b04 |
| Dr.23743.1.S1_at | wu:fj58g06 | wu:fj58g06 |
| Dr.12924.1.A1_at | wu:fj83e02 | wu:fj83e02 |
| Dr.6450.1.A1_at | wu:fj94a09 | wu:fj94a09 |
| Dr.23250.1.A1_at | wu:fk54e10 | wu:fk54e10 |
| Dr.24492.1.A1_at | wu:fl05f10 | wu:fl05f10 |
| Dr.14080.1.A1_at | wu:fq26c12 | wu:fq26c12 |
| Dr.11266.1.S1_at | "3-monooxygenase/tryptophan 5-monooxygenase activation protein, gamma polypeptide 1" | ywhag1 |
| Dr.2009.1.A1_at | "3-monooxygenase/tryptophan 5-monooxygenase activation protein, gamma polypeptide 2" | ywhag2 |
| Dr.19352.1.A1_at | zinc finger CCCH-type containing 10 | zc3h10 |
| Dr.18465.1.A1_at | "zinc finger, DHHC-type containing 3" | zdhhc3 |
| Dr.175.1.A1_at | zgc:103657 | zgc:103657 |
| Dr.21291.1.A1_at | zgc:109889 | zgc:109889 |
| Dr.15241.1.A1_at | zgc:110063 | zgc:110063 |
| Dr.26355.1.A1_at | zgc:111986 | zgc:111986 |
| Dr.14654.1.S1_at | zgc:114060 | zgc:114060 |
| Dr.26124.1.A1_at | zgc:114199 | zgc:114199 |
| Dr.12242.1.S1_at | zgc:123177 | zgc:123177 |
| Dr.14261.1.A1_at | zgc:123178 | zgc:123178 |
| Dr.2204.1.A1_at | zgc:136474 | zgc:136474 |
| Dr.24487.1.A1_at | zgc:136930 | zgc:136930 |
| Dr.11452.1.A1_at | zgc:153958 | zgc:153958 |
| Dr.22840.1.S1_at | zgc:158423 | zgc:158423 |
| Dr.14493.1.A1_at | zgc:158624 | zgc:158624 |
| Dr.4654.1.A1_at | zgc:162126 | zgc:162126 |
| Dr.9069.1.A1_at | zgc:162161 | zgc:162161 |
| Dr.11110.1.A1_at | zgc:162707 | zgc:162707 |
| Dr.7022.1.A1_at | zgc:165666 | zgc:165666 |
| Dr.15311.1.A1_at | zgc:175128 | zgc:175128 |
| Dr.11252.1.A1_at | zgc:56085 | zgc:56085 |

| | | | | | |
|---|---|---|---|---|---|
| Dr.12903.1.A1_at | leucine rich repeat and Ig domain containing 1b | lingo1b | Dr.26468.1.S1_at | zgc:63972 | zgc:63972 |
| Dr.6295.1.S1_at | LIM domain only 4a | lmo4a | Dr.17194.1.A1_at | zgc:65851 | zgc:65851 |
| Dr.19236.1.S1_at | LIM domain only 4b | lmo4b | Dr.7313.1.A1_at | zgc:73340 | zgc:73340 |
| Dr.14788.1.A1_at | transmembrane protein C9orf5-like | LOC100000119 | Dr.2980.1.A1_at | zgc:77439 | zgc:77439 |
| Dr.7559.1.S1_at | nuclear protein 1-like | LOC100000294 | Dr.15314.1.A1_at | zgc:77816 | zgc:77816 |
| Dr.14134.1.S1_at | synaptotagmin-C-like | LOC100000615 | Dr.18506.1.A1_at | zgc:77880 | zgc:77880 |
| Dr.9425.1.A1_at | novel protein similar to vertebrate adenylate cyclase family | LOC100003595 | Dr.22798.1.A1_at | zgc:85722 | zgc:85722 |
| Dr.20380.1.A1_at | RING finger protein 122-like | LOC100005267 | DrAffx.1.46.S1_at | zgc:86586 | zgc:86586 |
| Dr.17167.1.A1_at | "cadherin 11, type 2 preproprotein-like" | LOC100007192 | Dr.16545.1.A1_at | zgc:91811 | zgc:91811 |
| Dr.9619.1.A1_at | WSC domain-containing protein 1-like | LOC100329926 | Dr.5008.1.S1_at | zgc:92045 | zgc:92045 |
| Dr.17823.1.S1_at | 1-phosphatidylinositol phosphodiesterase-like \| zgc:174938 | LOC100330677 \| zgc:174938 | Dr.15620.1.S1_at | zgc:92046 | zgc:92046 |
| Dr.12487.1.A1_at | neuroendocrine protein 7B2-like \| secretogranin V | LOC100330746 \| scg5 | Dr.3620.1.A1_at | --- | ZNF770 |
| Dr.7100.1.A1_at | uncharacterized LOC100331484 | LOC100331484 | Dr.18421.1.A1_at | zinc and ring finger 1 | znrf1 |
| Dr.16574.1.A1_at | epoxide hydrolase 4-like | LOC100331939 | Dr.14548.1.A1_at | "zinc finger, RAN-binding domain containing 1b" | zranb1b |
| Dr.23961.1.S1_at | uncharacterized LOC100333176 \| uncharacterized LOC100534963 | LOC100333176 \| LOC100534963 | Dr.949.1.S1_at | "zinc finger, RAN-binding domain containing 2" | zranb2 |
| Dr.16437.1.A1_at | "leucyl-tRNA synthetase, cytoplasmic-like \| vascular endothelial zinc finger 1-like" | LOC100334939 \| LOC100535691 | Dr.5111.1.A1_at | septin 3 | 2-Sep |
| Dr.16969.1.S1_at | uncharacterized LOC100534666 | LOC100534666 | Dr.26136.1.A1_at | septin 6 | 5-Sep |
| Dr.7099.1.S1_at | "ribonuclease kappa-B-like \| ribonuclease, RNase K b" | LOC100534769 \| rnasekb | Dr.16787.1.A1_at | --- | --- |
| Dr.21458.1.S1_at | uncharacterized LOC100535266 | LOC100535266 | Dr.16930.1.A1_at | --- | --- |
| Dr.11344.2.S1_at | cytoplasmic polyadenylation element-binding protein 4-like | LOC100535549 | Dr.16584.1.S1_at | --- | --- |
| Dr.11344.1.A1_at | cytoplasmic polyadenylation element-binding protein 4-like | LOC100535549 | Dr.14100.1.S1_at | --- | --- |
| Dr.23415.1.A1_at | uncharacterized LOC100535624 | LOC100535624 | Dr.18101.1.A1_at | --- | --- |
| Dr.7666.1.S1_at | uncharacterized LOC100537083 | LOC100537083 | Dr.11151.1.A1_at | --- | --- |
| Dr.11172.2.S1_at | uncharacterized LOC100537668 | LOC100537668 | Dr.25445.1.A1_at | --- | --- |
| Dr.17159.1.A1_at | zinc finger protein 84-like | LOC100537687 | Dr.25318.1.A1_at | --- | --- |
| Dr.16932.1.A1_at | c-Jun-amino-terminal kinase-interacting protein 1-like | LOC100537717 | Dr.5685.1.S1_at | --- | --- |
| Dr.26296.1.A1_s_at | c-Jun-amino-terminal kinase-interacting protein 1-like | LOC100537717 | Dr.15131.1.A1_at | --- | --- |
| Dr.14801.1.S1_at | probable G-protein coupled receptor 75-like \| probable G-protein coupled receptor 75-like | LOC100538001 \| LOC561668 | Dr.13475.1.A1_at | --- | --- |
| Dr.14069.1.A1_at | calsyntenin-3-like | LOC555627 | Dr.22720.1.A1_at | --- | --- |
| Dr.7441.1.A1_at | glycerophosphodiester phosphodiesterase domain-containing protein 5-like | LOC560645 | Dr.13933.1.A1_at | --- | --- |
| Dr.21875.1.A1_at | "novel protein similar to H.sapiens TTC9, tetratricopeptide repeat domain 9 (TTC9)" | LOC560706 | Dr.14352.1.A1_at | --- | --- |
| Dr.23663.1.S1_at | novel protein similar to vertebrate atrophin 1 (ATN1) | LOC561408 | Dr.17958.1.S1_at | --- | --- |
| Dr.19528.1.A1_at | uncharacterized LOC562053 | LOC562053 | Dr.26299.1.A1_at | --- | --- |
| Dr.6172.1.A1_at | uncharacterized LOC567192 | LOC567192 | Dr.7509.1.A1_at | --- | --- |
| Dr.12540.1.A1_at | "novel protein similar to H.sapiens GALNTL6, UDP-N-acetyl-alpha-D-galactosamine:polypeptide N-acetylgalactosaminyltransferase-like 6 (GALNTL6)" | LOC568915 | Dr.16794.2.A1_a_at | --- | --- |
| Dr.12472.1.A1_at | syntaxin-12-like | LOC569124 | Dr.16786.1.A1_at | --- | --- |
| Dr.11095.1.A1_at | serine incorporator 4-like | LOC570112 | Dr.12642.1.A1_at | --- | --- |
| Dr.15891.1.A1_at | zinc finger protein 836-like \| zgc:174928 | LOC572225 \| zgc:174928 | Dr.6999.1.A1_at | --- | --- |
| Dr.13261.1.A1_at | pleckstrin homology domain-containing family G member 5-like | LOC796404 | Dr.8944.1.A1_at | --- | --- |

| | | |
|---|---|---|
| Dr.12598.1.S1_at | neural cell adhesion molecule 2 | ncam2 |
| Dr.12107.1.A1_at | N-myc downstream regulated gene 1a | ndrg1a |
| Dr.827.1.A1_at | "NADH dehydrogenase (ubiquinone) 1 alpha subcomplex, 4" | ndufa4 |
| Dr.16132.1.A1_at | "neurofilament, medium polypeptide a" | nefma |
| Dr.22710.1.A1_at | NEL-like 2a | nell2a |
| Dr.22741.1.A1_at | NEL-like 2b | nell2b |
| Dr.556.1.S1_at | neurogenic differentiation | neurod |
| Dr.5740.1.S1_at | neurogenic differentiation 2 | neurod2 |
| Dr.8196.1.S1_at | neurogenic differentiation 6a | neurod6a |
| Dr.8197.1.S1_at | neurogenic differentiation 6b | neurod6b |
| Dr.4859.1.A1_at | nexilin (F actin binding protein) | nexn |
| Dr.12913.1.A1_at | neuroligin 4a | nlgn4a |
| Dr.9836.1.S1_at | NME/NM23 nucleoside diphosphate kinase 2a | nme2a |
| Dr.7457.1.A1_at | nicotinamide nucleotide adenylyltransferase 2 | nmnat2 |
| Dr.4949.1.A1_at | nitric oxide synthase interacting protein | nosip |
| Dr.20524.1.A1_at | neuronal PAS domain protein 4a | npas4a |
| Dr.23064.2.S1_at | "neural proliferation, differentiation and control, 1" | npdc1 |
| Dr.7065.2.A1_at | natriuretic peptide precursor C-like protein | nppcl |
| Dr.10103.1.A1_at | neuroplastin a | nptna |
| Dr.14062.1.A1_at | neuregulin 3 | NRG3 |
| Dr.12859.1.S1_at | "neurogranin (protein kinase C substrate, RC3) a" | nrgna |
| Dr.7206.1.A1_at | neurensin 1 | nrsn1 |
| Dr.10477.1.S1_at | NSA2 ribosome biogenesis homolog (S. cerevisiae) | nsa2 |
| Dr.3959.1.A1_at | "5'-nucleotidase, cytosolic II, like 1" | nt5c2l1 |
| Dr.6259.1.S1_at | nucleobindin 2b | nucb2b |
| Dr.2992.1.A1_at | optic atrophy 1 (human) | opa1 |
| Dr.8194.1.S1_at | "opsin 1 (cone pigments), short-wave-sensitive 1" | opn1sw1 |
| Dr.19427.2.S1_at | --- | OSBPL2A |
| Dr.7597.1.S1_at | oxytocin | oxt |
| Dr.11392.1.A1_at | "poly A binding protein, cytoplasmic 1 b" | pabpc1b |
| Dr.26364.1.A1_at | "platelet-activating factor acetylhydrolase, isoform Ib, alpha subunit b" | pafah1b1b |
| Dr.15254.1.A1_at | "platelet-activating factor acetylhydrolase 1b, catalytic subunit 2" | pafah1b2 |
| Dr.2836.1.A1_at | paralemmin 1b | palm1b |
| Dr.12635.1.A1_at | progestin and adipoQ receptor family member IIIa | paqr3a |
| Dr.1697.1.A1_at | "poly (ADP-ribose) polymerase family, member 12b" | parp12b |
| Dr.22774.1.A1_at | piccolo (presynaptic cytomatrix protein) b | pclob |
| Dr.17014.1.S1_at | Purkinje cell protein 4 like 1 | pcp4l1 |
| Dr.13800.1.A1_at | peroxisomal biogenesis factor 5 | pex5 |
| Dr.6819.1.A1_at | phosphoglycerate mutase 1b | pgam1b |
| Dr.24191.1.S1_at | "phosphoinositide-3-kinase, regulatory subunit 3b (gamma)" | pik3r3b |
| Dr.11329.1.A1_at | "protein kinase (cAMP-dependent, catalytic) inhibitor beta" | pkib |
| Dr.7101.1.S1_at | pro-melanin concentrating hormone-like protein | pmchl |
| Dr.11138.1.A1_at | podocalyxin-like 2 | podxl2 |
| Dr.13574.1.S1_at | "POU domain, class 4, transcription factor 2" | pou4f2 |
| Dr.24898.1.S1_at | pancreatic progenitor cell differentiation and proliferation factor a | ppdpfa |
| Dr.26034.1.A1_at | "protein kinase, cAMP-dependent, regulatory, type I, alpha (tissue specific extinguisher 1) a" | prkar1aa |
| Dr.8139.1.S1_at | prospero-related homeobox gene 1a | prox1a |
| Dr.22807.1.A1_at | "pleckstrin and Sec7 domain containing 3, like" | psd3l |
| Dr.17113.1.A1_a_at | phosphatase and tensin homolog A | ptena |
| Dr.12800.1.S1_at | "protein tyrosine phosphatase-like (proline instead of catalytic arginine), member b" | ptplb |
| Dr.13952.1.A1_at | "protein tyrosine phosphatase, non-receptor type 11, b" | ptpn11b |
| Dr.14147.1.S1_at | parvalbumin 6 | pvalb6 |
| Dr.16940.1.S1_at | "RAB11B, member RAS oncogene family, b" | rab11bb |
| Dr.16940.2.A1_at | "RAB11B, member RAS oncogene family, b" | rab11bb |
| Dr.13273.1.S1_at | "RAB15, member RAS oncogene family" | rab15 |
| Dr.25650.1.A1_at | "RAB, member of RAS oncogene family-like 2" | rabl2 |
| Dr.6811.1.A1_at | "ras-related C3 botulinum toxin substrate 3a (rho family, small GTP binding protein Rac3)" | rac3a |

| | | |
|---|---|---|
| Dr.13222.1.A1_at | --- | --- |
| Dr.26519.2.A1_at | --- | --- |
| Dr.13465.1.A1_at | --- | --- |
| Dr.26398.2.S1_at | --- | --- |
| Dr.936.1.A1_at | --- | --- |
| Dr.25884.1.A1_at | --- | --- |
| Dr.14167.1.S1_at | --- | --- |
| Dr.12823.1.A1_at | --- | --- |
| Dr.17976.1.S1_at | --- | --- |
| Dr.16582.1.A1_at | --- | --- |
| Dr.15208.1.A1_at | --- | --- |
| Dr.17141.1.A1_at | --- | --- |
| Dr.16506.1.A1_at | --- | --- |
| Dr.16373.1.S1_at | --- | --- |
| Dr.11232.1.A1_at | --- | --- |
| Dr.17756.1.A1_at | --- | --- |
| Dr.12943.1.A1_at | --- | --- |
| Dr.20400.1.A1_at | --- | --- |
| Dr.14762.1.A1_at | --- | --- |
| Dr.13441.1.A1_at | --- | --- |
| Dr.17920.1.A1_at | --- | --- |
| Dr.16737.1.S1_at | --- | --- |
| Dr.12738.1.A1_at | --- | --- |
| Dr.26429.1.A1_at | --- | --- |
| Dr.15403.1.A1_at | --- | --- |
| Dr.26085.1.A1_at | --- | --- |
| Dr.12075.1.A1_at | --- | --- |
| Dr.18656.1.S1_at | --- | --- |
| Dr.17757.1.A1_at | --- | --- |
| Dr.11143.1.A1_at | --- | --- |
| Dr.15196.1.S1_at | --- | --- |
| Dr.16081.1.S1_at | --- | --- |
| Dr.9998.1.A1_at | --- | --- |
| Dr.17577.1.A1_at | --- | --- |
| Dr.11049.1.A1_at | --- | --- |
| Dr.11079.1.A1_at | --- | --- |
| Dr.8650.1.S1_at | --- | --- |
| Dr.17398.1.A1_at | --- | --- |
| Dr.9417.1.A1_at | --- | --- |
| Dr.12871.1.A1_at | --- | --- |
| Dr.26254.1.A1_at | --- | --- |
| Dr.23433.1.A1_at | --- | --- |
| Dr.20440.1.A1_at | --- | --- |
| Dr.13616.1.A1_at | --- | --- |
| Dr.241.1.A1_at | --- | --- |
| Dr.12564.1.A1_at | --- | --- |
| Dr.16063.1.S1_at | --- | --- |
| Dr.7363.1.A1_at | --- | --- |
| Dr.10124.1.A1_at | --- | --- |
| Dr.14333.1.A1_at | --- | --- |
| Dr.22707.1.A1_at | --- | --- |
| Dr.26174.1.A1_at | --- | --- |
| Dr.15471.1.A1_at | --- | --- |
| Dr.17957.1.A1_at | --- | --- |
| Dr.19556.2.S1_at | --- | --- |
| Dr.13529.1.A1_at | --- | --- |
| Dr.11376.1.A1_at | --- | --- |

| | | | | | | |
|---|---|---|---|---|---|---|
| Dr.14199.1.S1_at | uncharacterized LOC797202 | LOC797202 | | Dr.13538.1.S1_at | --- | --- |
| Dr.10186.1.A1_at | "phytanoyl-CoA dioxygenase, peroxisomal-like" | LOC798210 | | Dr.16719.1.A1_at | --- | --- |
| Dr.22391.1.A1_at | low density lipoprotein receptor-related protein 11 | lrp11 | | Dr.16311.1.A1_at | --- | --- |
| Dr.10168.2.S1_at | v-maf musculoaponeurotic fibrosarcoma (avian) oncogene homolog | maf | | Dr.14759.1.A1_at | --- | --- |
| Dr.4477.1.A1_at | "membrane associated guanylate kinase, WW and PDZ domain containing 1b" | magi1b | | Dr.20489.2.S1_at | --- | --- |
| Dr.16681.1.A1_at | MAM domain containing 1 | mamdc1 | | Dr.15245.1.A1_at | --- | --- |
| Dr.26245.1.A1_at | mitogen-activated protein kinase kinase kinase kinase 3 | MAP4K3 (1 of 2) | | Dr.22364.1.A1_at | --- | --- |
| Dr.16118.1.A1_at | microtubule-associated protein tau b | maptb | | Dr.11343.1.A1_at | --- | --- |
| Dr.20062.1.S1_at | transformed 3T3 cell double minute 4 homolog (mouse) | mdm4 | | Dr.11355.1.A1_at | --- | --- |
| Dr.8060.1.S1_at | mesoderm specific transcript | mest | | Dr.17052.1.A1_at | --- | --- |
| Dr.25285.1.S1_at | MID1 interacting protein 1b | mid1ip1b | | Dr.21822.2.A1_a_at | --- | --- |
| Dr.16892.1.A1_at | "membrane protein, palmitoylated 2b (MAGUK p55 subfamily member 2b)" | mpp2b | | Dr.24189.1.S1_at | --- | --- |
| Dr.17264.1.A1_at | musashi homolog 2b (Drosophila) | msi2b | | Dr.18354.2.A1_at | --- | --- |
| Dr.17628.1.A1_at | musashi homolog 2b (Drosophila) | msi2b | | Dr.14765.1.A1_at | --- | --- |
| Dr.5778.1.S1_at | myostatin b | mstnb | | Dr.5685.2.S1_at | --- | --- |
| Dr.15836.1.A1_at | metadherin a | mtdha | | Dr.26158.1.A1_at | --- | --- |
| Dr.18683.1.S1_at | myocilin | myoc | | Dr.14358.1.A1_at | --- | --- |

### Supplemental Figure 5: List of SAM negative genes

Knockdown of nhsb causes changes in gene expression in the retina. SAM analysis identified significanly downregulated genes in nhsb MO injected embryos compared to control
